# Symbiont replacement and subsequent parallel genome erosion reshape a dual obligate symbiosis in the aphid *Lachnus tropicalis*

**DOI:** 10.1101/2025.09.22.677894

**Authors:** Tomonari Nozaki, Yuuki Kobayashi, Mika Ikeda, Shuji Shigenobu

**Author notes:** Author for correspondence: Tomonari Nozaki.

## Abstract

Many insects rely on obligate microbial symbioses, often involving multiple partners. Although symbiont replacement is well-documented, how newly acquired and resident obligate symbionts adapt after such events remains unclear. Here, we investigate the dual obligate symbiosis of the aphid *Lachnus tropicalis*, where an ancestral *Serratia* lineage was replaced by a newly acquired another *Serratia* lineage while the primary symbiont *Buchnera* remained. Our metagenomic sequencing yielded complete genomes of *Buchnera* (0.42 Mb) and *Serratia* (2.8 Mb), revealing developing metabolic complementarity. Although the *Serratia* genome retained abundant gene sets for amino acid synthesis, it also contained pseudogenes in leucine and methionine pathways, which would be compensated for by *Buchnera* or the host. Comparison with *L. roboris*, which harbors the ancestral *Serratia* lineage, showed that the newly acquired *Serratia* in *L. tropicalis* exhibits identical tissue localization and vertical transmission pattern, suggesting the smooth succession of the prior’s microniche. Notably, *Buchnera* in *L. tropicalis* exhibited a slightly more degenerated genome than its counterpart in *L. roboris*, indicating that symbiont replacement can accelerate gene loss even in ancient symbionts. Overall, our findings provide new insights into the dynamics of novel mutualism establishment and highlight symbiont replacement as a driver of host-symbiont co-evolution.

## 1. Introduction

Many insects have established highly integrated symbiotic relationships with microorganisms such as bacteria and fungi (1, 2, 3). To date, a large number of highly integrated insect–microbe symbiotic systems have been discovered, highlighting microbial symbiosis as an evolutionary driver (4, 5). Some mutualistic insect–bacterial relationships are ancient, implying enduring partnerships (6, 7, 8). However, “symbiont replacements,” in which older relationships are abandoned for newly acquired microbial partners, have been documented as evolutionary outcomes in many insect groups, including those harboring ancient symbionts (4, 9). Whether this phenomenon results from selection and adaptive evolution, or simply an escape from the “symbiont rabbit hole” to compensate for overly degenerated genomes, remains unresolved (5, 8, 10, 11). Comparative studies focusing on before and after replacement, including the states of the replaced and replacing symbionts, the host, any remaining co-symbionts, and the mechanisms by which new stable symbiotic systems are established and maintained, are crucial for advancing this debate.

Before symbiont replacement, existing partners may show signs of instability, such as genomic fragmentation (12), which can precede replacement by new partners, as observed in cicadas (13). After replacement, newly integrated symbionts typically undergo rapid microniche adaptation, including genome degeneration and restricted localization owing to bottlenecks and isolation (8, 11, 14, 15, 16). This process is also hypothesized to accelerate genomic degeneration in ancient co-resident symbionts, particularly if the new partner has broader nutritional capacities, leading to microniche reallocation (16, 17). Despite these emerging insights, the full extent of the genomic consequences and their validation across diverse systems remains unresolved. Moreover, it is unclear how host mechanisms for symbiosis (such as localization restriction, vertical transmission, and nutrient exchange across host–symbiont membranes) change post-replacement, or how new symbionts integrate into established systems. Indeed, newly acquired symbionts exhibit varied localization and transmission modes, suggesting flexibility in both host and symbiont (18, 19, 20). Answering these questions requires detailed comparative studies across recently replaced and closely related symbiotic systems.

Aphid (Hemiptera: Aphididae) symbiotic systems and their functions have been well-studied (21, 22), and growing research has detailed independently evolved, complex symbioses involving multiple obligate symbionts across subfamilies (23, 24, 25, 26, 27). Within the subfamily Lachninae, the widespread association with *Serratia symbiotica* and comprehensive symbiont diversity studies have led to specific symbiont replacement hypotheses (26, 28, 29, 30). Detailed genome-based studies of *Cinara* spp. and related clades suggest a dynamic process: the common ancestor of Lachninae established a complex symbiosis with both *Buchnera* and *Serratia* (30, 32, 33). During subsequent diversification, *Buchnera* was maintained, but the ancestral *Serratia* was repeatedly replaced by diverse bacterial lineages, including *Sodalis*, *Fukatsuia*, *Erwinia,* and other *Serratia* strains (30, 31). However, the precise events before and after these replacements remain unclear. For example, understanding host changes upon new symbiont entry, how new partners adapt to the insect body and novel bacteriome niches, and how resident *Buchnera* are impacted requires detailed comparative analyses of closely related species.

Within the aphid subfamily Lachninae, the genus *Lachnus* represents a unique case of symbiont replacement. Previous studies have indicated a co-obligate symbiotic system in more basal *Lachnus* species, such as *Lachnus roboris*, comprising *Buchnera* alongside a *Serratia symbiotica* lineage known as “Clade B” (Figure 1A, S1, S2, Supplementary Information Chapter 1; 26, 28, 29, 30). This *Serratia* lineage is characterized by coccoid morphology and reduced genomes, a pattern observed in strains from *Cinara cedri* and *Tuberolachnus salignus* (32, 33). Conversely, *Lachnus tropicalis* (Figure 1B) and other East and Southeast Asian species harbor a different *Serratia* lineage, Clade A (rod-shaped and retaining relatively large genomes), as the partner symbiont alongside *Buchnera* (Figure 1A, S1). Based on the symbiont replacement hypothesis in *Lachnus* (Figure 1A, Supplementary Information Chapter 1; 30), these Clade A *Serratia* strains are considered relatively newly integrated partners. Therefore, the genus *Lachnus* is an ideal system for comprehensively describing and comparing pre- and post-replacement symbiotic systems, offering crucial insights into the factors driving symbiont replacement and its evolutionary consequences.

**Figure 1.**
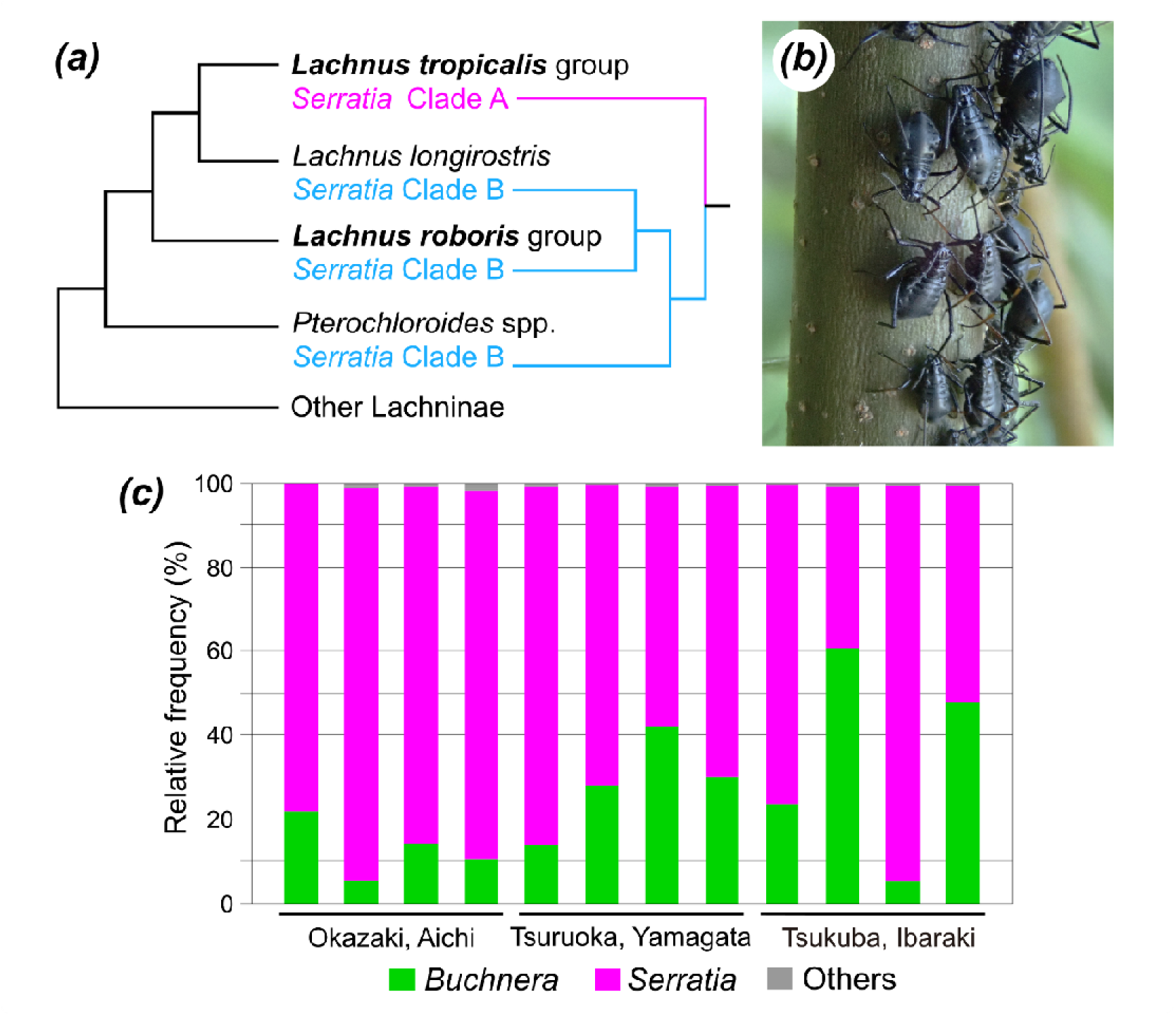
**(a)** Simplified phylogenetic tree of *Lachnus* and related species, showing the associated *Serratia* clades (“A” or “B”) harbored by aphids in addition to *Buchnera.* This tree suggests a symbiont replacement event from *Serratia* Clade B to Clade A in the *L. tropicalis* group (29, 30). **(b)** Colony of the black chestnut aphid, *Lachnus tropicalis* (Okazaki, Aichi). **(c)** Microbial diversity associated with *L*. *tropicalis* from different populations in Japan, based on amplicon sequencing of the hypervariable V3/V4 region of the 16S rRNA gene.

In this study, we first comprehensively characterized the symbiotic system of *L. tropicalis*, the great chestnut aphid. We used amplicon and metagenomic sequencing to describe the microbiome and obtain the complete genomes of its bacterial symbionts, *Buchnera aphidicola* (*Buchnera* Lt) and *Serratia symbiotica* (*Serratia* Lt). As expected, our analysis revealed that *Serratia* Lt exhibited the characteristics of a recently acquired symbiont and was well-integrated into the pre-existing symbiosis with the aphid and *Buchnera*. We combined these genomic insights with detailed observations of symbiont localization and vertical transmission to elucidate how *Serratia* Lt took over the microniche of its ancestral counterpart. This description was then contextualized by comparison with the symbiotic system of *L. roboris*, which harbors ancestral *Serratia*, based on a thorough review of the existing literature (26, 30, 34). Furthermore, using the recently sequenced *L. roboris Buchnera* genome (35), we investigated the evolutionary consequences for *Buchnera* following symbiont replacement.

## 2. Materials and methods

### (a) Characterization of the symbiotic system in *L. tropicalis*

#### **(i)** Sample collection

Between 2021 and 2024, we collected *L. tropicalis* from three localities in Japan: Okazaki (Aichi Prefecture), Tsuruoka (Yamagata Prefecture), and Tsukuba (Ibaraki Prefecture). When aphid colonies were observed on twigs of Fagaceae trees, individuals were carefully collected using aspirators or forceps. Upon collection, they were morphologically identified as *L. tropicalis* based on the distinct forewing pattern of winged adults and further confirmed by sequencing the mitochondrial cytochrome c oxidase subunit I gene region, commonly used for aphid species identification. Aphid samples were preserved in 99.5% ethanol at 4 °C or at room temperature (approximately 20–28 °C) until DNA extraction for amplicon sequencing. For metagenomic sequencing and imaging analyses, fresh samples were collected from the National Institute for Basic Biology (NIBB) Campus (Okazaki, Aichi).

#### (ii) 16S ribosomal DNA (rDNA) amplicon sequencing for *L. tropicalis* microbiome

To clarify the bacterial diversity associated with *L. tropicalis*, we conducted amplicon sequencing of the hypervariable V3/V4 region of the bacterial 16S ribosomal RNA (rRNA) gene using 12 individuals from the three geographically distinct localities mentioned above. Libraries were constructed according to the 16S rRNA Metagenomic Sequencing Guide provided by Illumina (USA) and sequenced on the Illumina MiSeq platform. Detailed methods are provided in the Supplementary Information (Supplementary Information Chapter 2). Raw reads were deposited in the DDBJ database under accession numbers DRR709987-DRR709998 (PRJDB35790). Raw paired-end reads were analyzed using QIIME 2 (version 2020.8) (37) with the plugin “dada2” (37) for quality filtering, trimming of read length, merging paired reads, and removing chimeric sequences. We excluded dada2-derived amplicon sequence variants (ASVs) with <100 reads and manually combined those with >99% sequence identity. The four resulting ASVs were manually assigned to genus-level taxa using the BLAST function in the National Center for Biotechnology Information (NCBI) database.

#### (iii) Metagenomic sequencing of the endosymbionts in *L. tropicalis*

To characterize the genomic features of *L. tropicalis* endosymbionts (*Buchnera* Lt and *Serratia* Lt), we performed metagenomic sequencing and subsequently assembled bacterial genomes using a hybrid assembly approach. Briefly, high-quality genomic DNA was extracted from fresh aphid samples; Nanopore and Illumina libraries were then prepared and sequenced using the R9.4.1 flow cell on the GridION system and the Illumina HiSeq X Ten platform, respectively. The total number of raw Nanopore reads was 2,038,881. The raw Nanopore reads were deposited in the DDBJ database under the accession number DRR718948 (PRJDB35790). The total number of raw Illumina paired-end reads obtained was 203,514,513. The raw Illumina reads were deposited in the DDBJ database under accession number DRR718949. To generate complete symbiont genomes from hologenomic samples, we carried out a hybrid assembly strategy involving long-read backbone assembly and short-read error correction. Detailed experimental procedures and assembly strategies are described in the Supplementary Information (Supplementary Information Chapter 3). The assembled genome and plasmid sequences were deposited in the DDBJ database under accession numbers: AP043953 and AP043954 for *Buchnera* Lt, and AP043955-AP043957 for *Serratia* Lt. Annotation of the corrected genomes was performed using DDBJ Fast Annotation and Submission Tool (DFAST) version 1.2.18 (38). We manually corrected the DFAST annotation for some genes, re-evaluating pseudogenes that appeared functional based on comparisons with related genomes. To characterize the gene repertoires of both symbionts in *L. tropcialis*, we examined the Cluster of Orthologous Genes (COG) category tags (39). This information was compared with that of related genomes (see Supplementary Information Chapter 3). The genome was visualized using Proksee (40), and the resulting images were processed with Inkscape version 1.4 (86a8ad7, 2024-10-11) (41). Metabolic pathways of the two symbiont genomes were reconstructed using the Kyoto Encyclopedia of Genes and Genomes Mapper tool (42), considering pseudogenization information obtained from the DFAST results (for details, see Supplementary Information Chapter 4).

#### **(iv)** Phylogenomic analysis

To infer the phylogeny of *Buchnera* and *Serratia* in *L. tropicalis* based on their genomic information, separate phylogenetic trees were constructed for each bacterial genus using GToTree v1.8.14 (43) and its prepackaged single-copy gene set for Gammaproteobacteria (172 targets). Briefly, protein-coding genes were predicted from input genome FASTA files (listed in Tables S7 and S8; details in Supplementary Information Chapter 3) using Prodigal v2.6.3 (44). Targeted single-copy genes were identified with HMMER3 v3.4 (45), individually aligned using MUSCLE v5.1 (46), and trimmed with TrimAl v1.5. rev0 (47), and concatenated. Phylogenetic trees were subsequently estimated using IQ-TREE v2.4.0 (48). The resulting tree files were visualized using Interactive Tree of Life (iTOL) v6 (49) and manually refined with Inkscape.

#### (v) Histological observations of *L. tropicalis* bacteriome and symbiont localization

To characterize the cellular features of the *L. tropicalis* bacteriome, we conducted morphological observations of the dissected bacteriomes with 4’,6-diamidino-2-phenylindole (DAPI) and Phalloidin staining for nuclei/DNA and F-actin, respectively. Furthermore, to visualize the tissue localization and vertical transmission of both symbionts in *L. tropicalis*, we conducted fluorescence in situ hybridization (FISH) on the bacteriome and viviparous embryos, using symbiont-specific probes targeting the 16S rRNA gene sequences (50). All stained or hybridized samples were observed under a confocal microscope FV1000 (Olympus, Japan). Detailed procedures and definitions of the embryonic developmental stages are described in the Supplementary Information (Supplementary Information Chapter 5).

### (b) Comparative analysis of *L. roboris* and *L. tropicalis* symbiotic systems

To compare symbiotic systems before and after symbiont replacement, we surveyed existing information on *L. roboris*. This European and Middle Eastern species forms a clade distinct from its Asiatic counterparts, including *L. tropicalis* (51, 52). We used the symbiotic system of *L. roboris*, which harbors both *Buchnera* and an ancestral lineage of *Serratia symbiotica* (Clade B), as a model for the pre-replacement state (Supplementary Information Chapter 6). This *Serratia* is closely related to symbionts found in *Cinara cedri* and *Tuberolachnus salignus* (Figure S2; 29, 30, 32) (see Supplementary Information Chapter 1).

We reviewed bacteria taxa consistently detected in the microbiome (26, 28, 29) and re-analyzed the recently sequenced *Buchnera* draft genome (35). We downloaded the *L. roboris Buchnera* genome (Zenodo: https://zenodo.org/records/10513209) and re-annotated it using DFAST. To infer genomic differences between *Buchnera* from *L. roboris* and *L. tropicalis*, we used Proksee for visualization and sequence comparison, along with BLAST+ v2.16.0 (53) and FastANI v1.34 (54). To detect the presence or absence of genes in both *Buchnera* genomes, we performed an orthology analysis using OrthoFinder v3.1.0 (55, with *Buchnera* APS (GCF_000009605) as the outgroup.

Notably, detailed histological observations exist for *L. roboris* (under its synonym *Pterochlorus roboris*) concerning symbiont tissue localization and vertical transmission (34). We thoroughly reviewed this work and compared its features with those of the *L. tropicalis* symbiotic system (see Supplementary Information Chapter 5). As specific microbial species such as *Buchnera* and *Serratia* were not formally described in 1927, we first correlated Klevenhausen’s descriptions with our current understanding of symbiont identities (see Results).

## 3. Results

### (a) Description of the symbiotic system in *L. tropicalis*

#### (i) Microbiome of *L. tropicalis*

In our 16S rDNA amplicon sequencing analysis of *L. tropicalis* collected from three geographically distinct localities in Japan (Table S3), we consistently detected *Buchnera* and *Serratia* across all sampled localities and individuals (Figure 1C). Both symbionts were abundant within the *L. tropicalis* microbiome (mean ± standard deviation: *Buchnera* 25.5% ± 17.4%; *Serratia* 73.9% ± 17.3%). The *Serratia* partial sequence (427 bp) showed a 100% match with the deposited 16S rDNA sequences of *Serratia* symbionts from *L. tropicalis*, *L. siniquercus,* and *L. yunlongensis* (FJ655545, KP866556, KP866555, KP866552, and KF751207). Therefore, *Serratia* detected in the analysis belongs to *Serratia symbiotica* “Clade A,” which contains pathogenic, facultative, and obligate symbionts in aphids (56, 57). This result was consistent with phylogenetic analysis using the nearly full-length 16S rDNA extracted from the *Serratia* genome assembled in this study (Figure S2).

#### (ii) Genomic characterization of *Buchnera* and *Serratia* in *L. tropicalis*

We performed shotgun hologenome sequencing of *L. tropicalis*, and our metagenomic assembly yielded complete genomes for two bacterial symbionts: *Buchnera* Lt and *Serratia* Lt (Figures 2 and S3, Table 1; for more details, see Supplementary Information Chapter 3).

**Figure 2.**
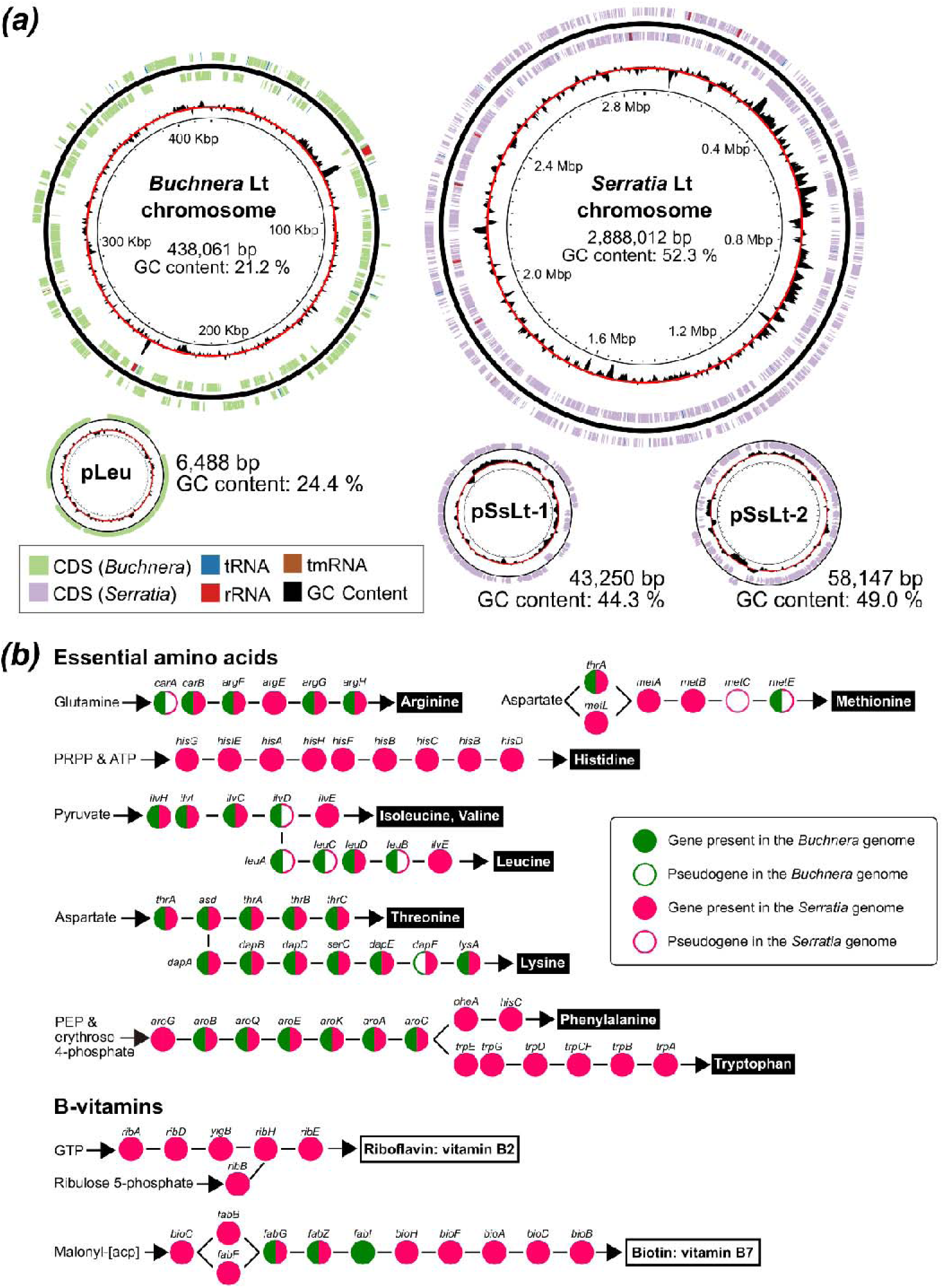
**(a)** Circular maps of *Lachnus tropicalis* symbiont genomes. The chromosomes of the primary symbiont *Buchnera aphidicola* Lt and *Serratia symbiotica* Lt, as well as their plasmids: pLeu for *Buchnera*, and pSsLt-1 and pSsLt-2 for *Serratia*. The rings, from outermost to innermost, represent: (i) predicted protein-coding genes, tRNAs, tmRNAs, and rRNAs on the plus strand; (ii) genome backbone; (iii) predicted protein-coding genes, tRNAs, tmRNAs, and rRNAs on the minus strand; (iv) GC content (deviation from the average); and (v) genome coordinates in megabases. Both chromosomes are oriented with the reading frame of the *dnaA* gene as the first CDS on the forward strand. The *Buchnera* plasmid pLeu is oriented at *repA*, while the *Serratia* plasmids begin at arbitrary positions. **(b)** Metabolic complementation for the biosynthesis of 10 essential amino acids (EAAs) and B-vitamins (B2 and B7) in the *L. tropicalis* endosymbiotic system. The *Serratia* Lt genome shows a relatively broad capacity for EAA synthesis, but pseudogenization was detected in the methionine (*metC* and *metE*) and leucine (*leuA*, *leuC*, and *leuB*) pathways. These functions are presumably complemented by *Buchnera* (for *metE*) or the aphid host (for *metC*). All leucine genes are located on the *Buchnera* plasmid pLeu, underscoring the complementary nature of the two genomes. Conversely, the synthesis of histidine and tryptophan is entirely dependent on *Serratia*.

**Table 1.**
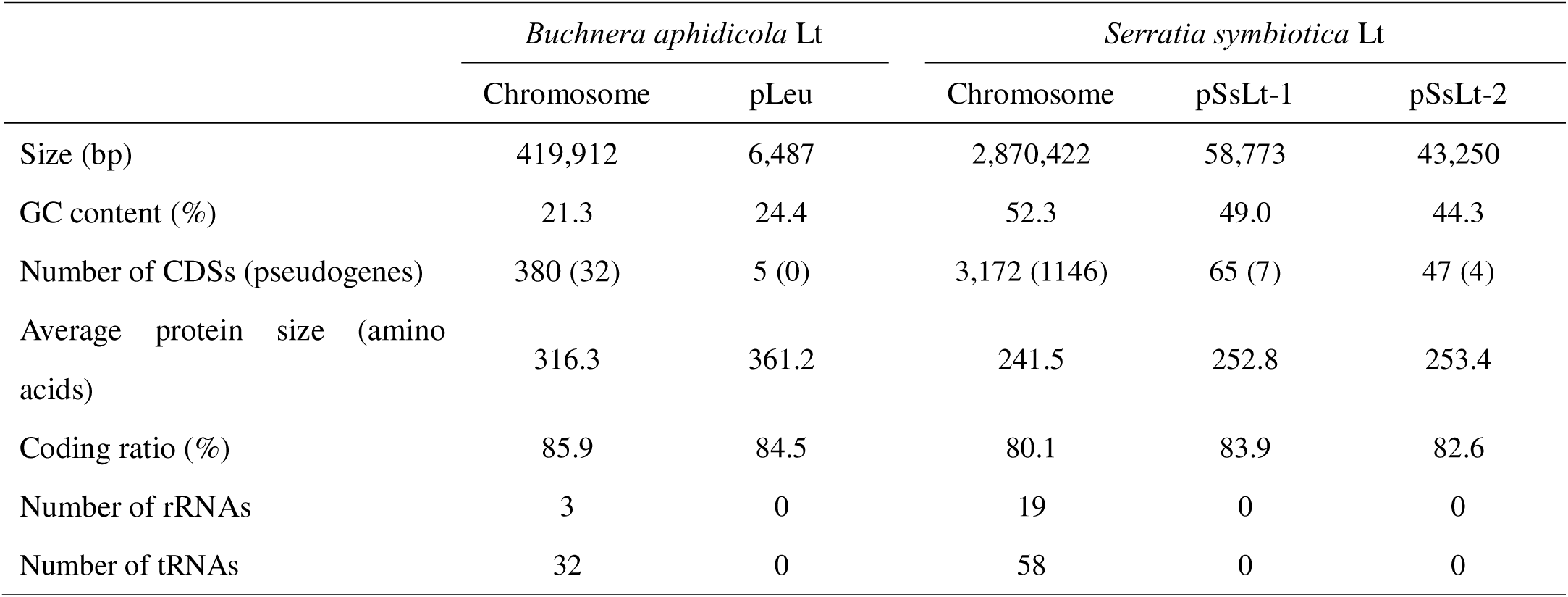
Summary of assembled genomes of *Buchnera* and *Serratia* in *Lachnus tropicalis*.

The *Buchnera* Lt genome comprised a 419,912 bp circular chromosome (21.3% GC content, 85.9% coding density) and a 6,487 bp pLeu plasmid encoding five leucine biosynthesis genes (Figure 2A, Table 1). Although considerably smaller than the mono-symbiotic *Buchnera* genomes (approximately 600 kb) (58) found in many aphid species (6, 22, 59), this reduction is typical for *Buchnera* in Lachninae aphids (23, 31, 32, 33), all of which co-exist with another obligate symbiont (*S. symbiotica* or other bacteria; Table S4). DFAST annotation revealed that the *Buchnera* Lt chromosome encodes 380 protein-coding genes (CDSs), three rRNAs, 32 transfer RNAs (tRNAs), and 32 pseudogenes. Notably, the pTrp plasmid, common in most *Buchnera* lineages, was not detected. We confirmed this absence through three independent analyses: plasmid assembly using “plassembler” (60), long-read assembly (no reads exceeding 5kb), and short-read mapping on pTrp of *Buchnera* APS and *Buchnera* from *Cinara cedri* using “bbmap.” All analyses failed to detect pTrp, indicating that *Buchnera* Lt lacks the tryptophan plasmid. COG analysis showed *Buchnera* Lt had gene repertoires similar to *Buchnera* in *Cinara cedri* (Figure S4). However, *Buchnera* Lt exhibited more pseudogenes (9.07%) than *Buchnera* genomes from pea aphids (0% in APS) and *Cinara cedri* (0.82%) (Figure S4, Table S4), suggesting ongoing genomic degeneration.

The *Serratia* Lt genome consisted of a 2,870,422 bp circular chromosome (52.3% GC content, 80.1% coding density) and two plasmids, pSsLt-1 and pSsLt-2 (Figure 2A, Table 1). Its total genome size (2.97 Mb) is comparable to that of *Serratia* in *Acyrthosiphon pisum* (2.82 Mb), a facultative symbiont (61), and *Serratia* in *Peryphyllus lyropictus* (3.15 Mb), a recently acquired co-obligate partner (25). Overall, the *Serratia* Lt genome encoded 3,172 proteins, including 1,146 pseudogenes, 19 rRNAs, and 58 tRNAs. Its normal GC content, typical number of rRNA operons (five full sets and two partial sets of 16S and 23S rRNA), and high pseudogene count indicate a relatively recent establishment as an aphid endosymbiont (Table S5). Furthermore, COG analysis revealed that *Serratia* in *L. tropicalis* retained as many genes as *Serratia* strains IS and CWBI-2 and did not exhibit the functional degeneration observed in *Serratia* from *C. cedri* (Figure S4). Our phylogenomic analysis, utilizing the newly sequenced genomes together with deposited related species (Tables S6 and S7), confirmed the phylogenetic placement of *L. tropicalis Buchnera* and *Serratia*. *Buchnera* Lt clustered within Lachninae and was most closely related to *Buchnera* in *L. roboris* (Figure S5). *Serratia* Lt was placed in Clade A of *S. symbiotica*, with its closest relative being *Serratia* in *Cinara tujaphilina* (Figure S6).

We then compared the gene repertoires of *Buchnera* Lt and *Serratia* Lt to determine their co-obligate associations with host aphids. This analysis revealed complementary metabolic pathways for essential amino acids (EAAs) and B vitamins, demonstrating a clear metabolic complementarity between the two symbionts (Figures 2B, S7, S8; Tables S8, S9; Supplementary Information Chapter 4). Notably, the *Buchnera* Lt genome lacked histidine biosynthesis genes and the tryptophan plasmid, representing the most degraded EAA synthesis capacity observed in this bacterial clade. Although *Serratia* in *L. tropicalis* possessed abundant EAA gene sets, its genome contained pseudogenes for leucine biosynthesis (compensated by *Buchnera*’s plasmid) and for later steps in methionine production (likely covered by *Buchnera* and the host aphid) (Figure 2B, S7). For cofactor biosynthesis, including B vitamins, *Serratia* Lt retained nearly all genes required to complete these pathways (Figures 2B, S8). To infer the evolutionary stage of the symbionts, we also examined genes related to cell wall synthesis, cell division, and cellular motility in *Serratia Lt* and *Buchnera* Lt. As expected, *Buchnera* Lt exhibited extensive genome degradation typical of *Buchnera* in Lachninae, Chaitophorinae, and Hormaphidinae (23, 24, 25, 26, 27). In contrast, *Serratia* Lt showed an ongoing process of functional degeneration in these pathways, suggesting that it is transitioning toward a more highly integrated symbiotic lifestyle (Tables S10, S11, S12; Supplementary Information Chapter 4).

#### **(iii)** Symbiont localization, vertical transmission, and bacteriome formation in *L. tropicalis*

Aphid symbiotic organ (bacteriome) generally consists of two types of cells: bacteriocytes, which house the obligate symbiont *Buchnera*, and sheath cells, which often harbor facultative symbionts (34, 62, 63, 64). Bacteriocytes are large polyploid cells (approximately 256 ploidy in *Acyrthosiphon pisum*) that serve as interfaces for nutritional interactions. Although some aphids with “multiple obligate symbionts,” such as *Cinara cedri* and *Ceratovacuna japonica*, harbor each symbiont in distinct bacteriocytes, sheath cells are typically either free of symbionts or house facultative ones (20, 24, 65) (Figure S9). Our morphological observations and FISH revealed that both *Buchnera* and *Serratia* were localized within the bacteriome of *L. tropicalis* (Figure 3). Consistent with other aphids, the primary aphid symbiont, *Buchnera*, resided in large polyploid bacteriocytes. In contrast, *Serratia* was localized in smaller sheath cells. We also observed that both symbionts exhibited distinct modes of maternal vertical transmission (Figures 4A–I, S10, and S11). Detailed observations of embryos from viviparous individuals showed that, following transmission, the two symbionts were distributed separately into embryonic bacteriome cells in a complex and integrated manner (Supplementary Information Chapter 5).

**Figure 3.**
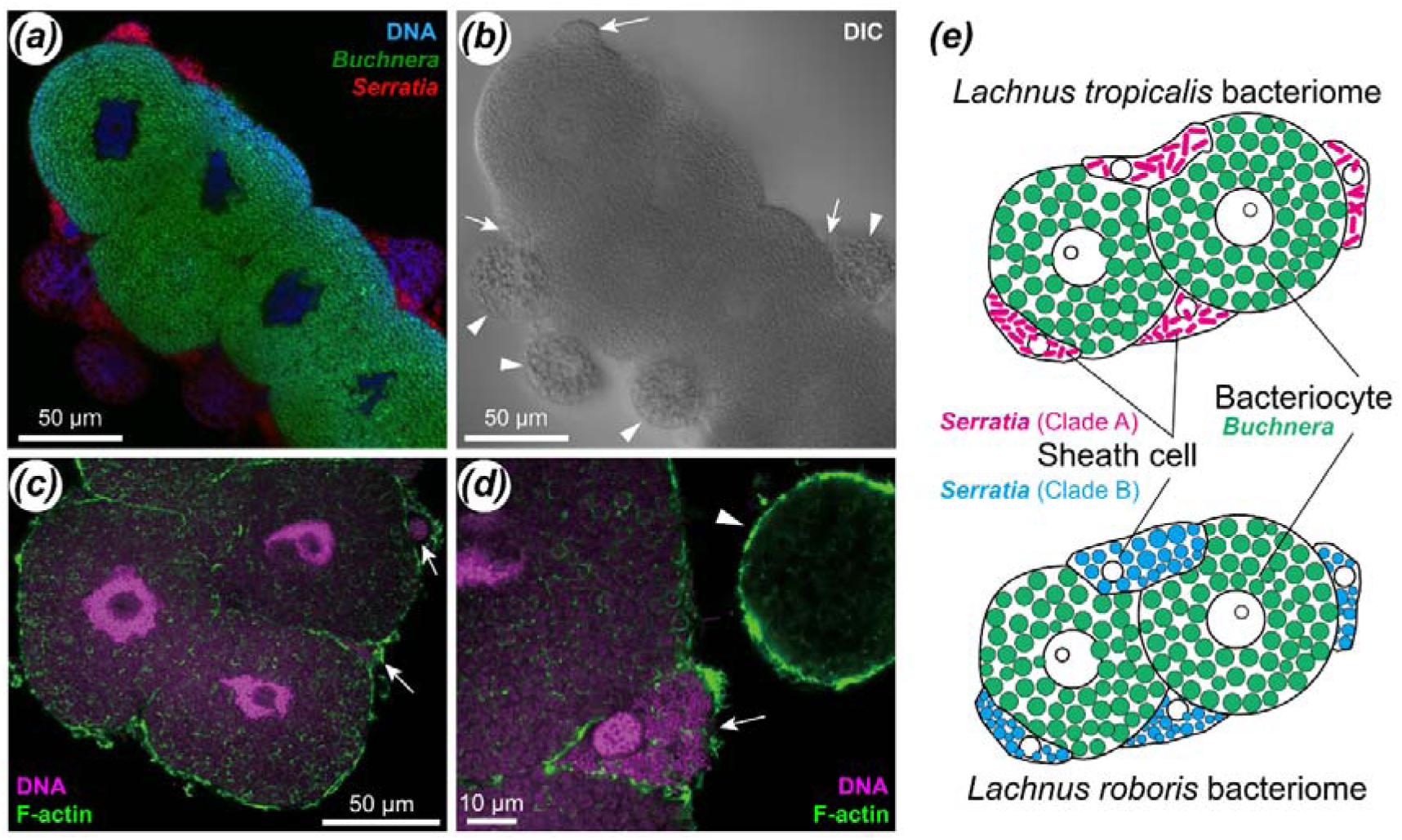
Symbiont localization and bacteriome structure in *Lachnus tropicalis* and *L. roboris*. **(a–d)** Symbiont localization and bacteriome structure in *Lachnus tropicalis*. **(a)** Fluorescent in situ hybridization (FISH) visualization of *Buchnera* (green) and *Serratia* (red) *in vivo*. *Buchnera* is harbored in large, polyploid bacteriocytes, whereas *Serratia* occupies flattened sheath cells. **(b)** Differential interference contrast image of the same tissue shown in (A). Arrows indicate sheath cells, and arrowheads denote fat cells. **(c, d)** DAPI-Phalloidin staining of the bacteriome. DNA (magenta) and F-actin (green) highlight cellular structures. *Buchnera* (round-shaped) and *Serratia* (rod-shaped) cells were confirmed to be housed in the bacteriocytes and sheath cells (arrows), respectively. Fat cells (arrowheads) contain no symbionts. **(e)** Comparison of bacteriome structure and symbiont localization between *L. tropicalis* (this study) and *L. roboris* (34). In both species, *Buchnera* is localized within bacteriocytes, while *Serratia* symbionts are confined to sheath cells. Notably, these observations revealed no major differences between the two species, except for the distinct cellular morphology of *Serratia* symbionts.

**Figure 4.**
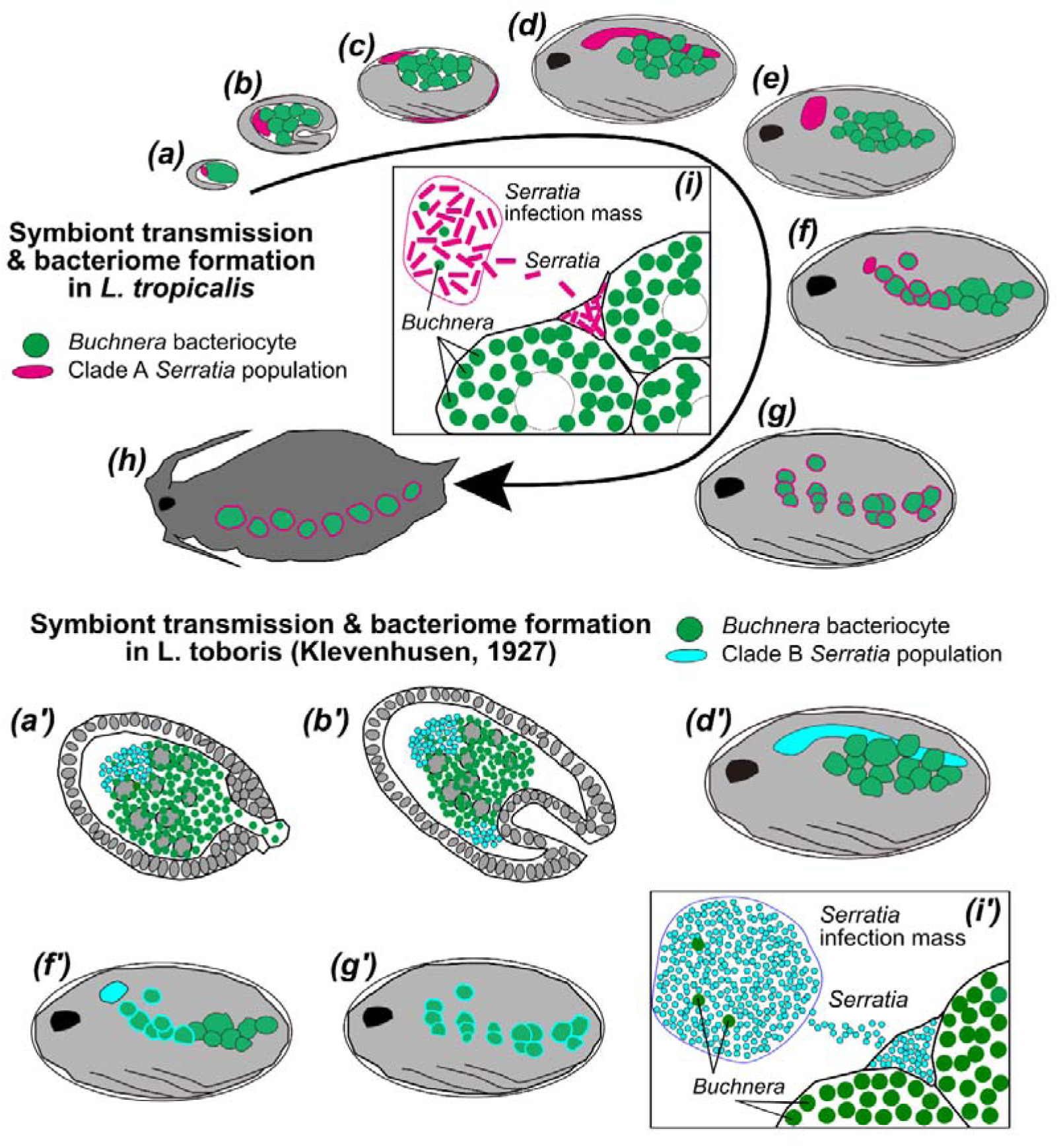
Vertical transmission, bacteriome formation, and symbiont distribution during viviparous embryogenesis in *Lachnus tropicalis* and *L. roboris*. **(a–h)** Schematic illustrations of *L. tropicalis* embryonic development, showing symbiont localization. *Buchnera* is indicated in green, and *Serratia* (Clade A, rod-shaped) in magenta. **(a)** Transmission stage. **(b)** Invagination (anatorepsis) stage. **(c)** Flip (katatrepsis) stage. **(d–g)** Final growth stage, during which *Serratia* cells sequentially infect sheath cells from a dorsally located infection mass. **(h)** Reconstruction of the bacteriome arrangement in adults (based on anatomical observations), reflecting the organization established during embryogenesis. **(i)** Schematic illustration of the *Serratia* infection mass and subsequent infection of sheath cells, showing the presence of several *Buchnera* cells within the infection mass. More detailed descriptions are provided in Supplementary Information Chapter 4, Figures S10 and S11. **(a’,b’, d’, f’, g’, i’)** Illustrations of the corresponding stages in *L. roboris*, based on 34. Each primed letter (such as a’) corresponds to the equivalent stage observed in *L. tropicalis*. *Buchnera* and bacteriocytes are shown in green, while *Serratia* (Clade B, round-shaped), its infection mass, and *Serratia*-containing sheath cells are depicted in cyan. Notably, no observable differences were found between *L. roboris* and *L. tropicalis*, except for the distinct cellular morphology of *Serratia*.

### (b) Comparison of symbiotic systems of *L. tropicalis* and *L. roboris*

We re-examined the existing literature on the symbiotic system of *L. roboris,* the species with the most documented research within the genus (Supplementary Information Chapter 6). This included a review of early histological observations by Klevenhausen (34), who first described three morphologically distinct symbionts of the aphid: a common round-shaped symbiont, a small round-shaped symbiont, and a rod-shaped symbiont. Building on this histological groundwork, recent phylogenetic analyses and amplicon sequencing have definitively reinterpreted these findings. The symbionts have been identified as *Buchnera* and *Serratia* belonging to Clade B, along with guest symbionts such as *Wolbachia* (Figures S1, S2; 26, 28, 29, 30). *Serratia* is characterized by its round shape, unique tissue localization, vertical transmission mode, and sophisticated distribution in sheath cells during embryonic development (Figure 4; 34).

We first compared the anatomical features, including tissue localization, vertical transmission of symbionts, and bacteriome formation during embryogenesis, between the *Buchnera/Serratia* (Clade B) symbiosis in *L. roboris* and the *Buchnera*/*Serratia* (Clade A) symbiosis in *L. tropicalis* (Table 2). We found no significant differences in symbiont localization (Figure 3E). In both aphid species, only *Buchnera* occupied the bacteriocytes, the main cell type comprising the symbiotic organs of aphids. In contrast, both *Serratia* lineages were localized in the sheath cells, which are flattened cells surrounding the bacteriocytes. Vertical transmission modes were nearly identical between the two *Lachnus* species (Figure 4). *Buchnera* cells were first transferred to infect early-stage embryos. Subsequently, *Serratia* cell populations were transmitted, splitting the *Buchnera* mass in half. After transmission, the mass of *Serratia* cells formed a dome-like structure positioned over the *Buchnera* mass at the anterior pole of the embryo.

**Table 2.** Summary of comparison of symbiotic system in two *Lachnus* species

|  | *Lachnus tropicalis* | *Lachnus roboris* |
| --- | --- | --- |
| Distribution | East and Southeast Asia, including India and East Siberia | Europe, North Africa, the Middle East, Central Asia |
| Host plants | *Quercus* spp., *Lithocarpus* spp.,<br>*Castanea* spp., *Casuarina equisetifolia* | *Quercus* spp., *Castanea sativa* |
| Primary obligate symbiont | *Buchnera aphidicola* | *Buchnera aphidicola* |
| Genome size of *Buchnera* | 0.42 Mb | 0.42 Mb |
| EAA synthesis pathways | Arg, Met, Val, Leu, Ile and Thr | Arg, Met, Val, **Leu**, Ile, Thr and **Trp** |
| *Buchnera* localization | Bacteriocytes | Bacteriocytes |
| Secondary obligate symbiont | *Serratia symbiotica* (clade A) | *Serratia symbiotica* (clade B) |
| Genome size of *Serratia* | 2.9 Mb | 0.6 ~ 2.0 Mb? |
| EAA synthesis pathways* | Arg, Met, His, Lys, Phe and Trp | Arg, Met, His, Lys and Phe |
| *Serratia* localization | Sheath cells | Sheath cells, trachea and fat cells |
| Shape of *Serratia* | Rod-shape | Small round shape |
| Vertical transmission mode | Almost same as each other; see Fig. 4 |  |

\* Information on *Serratia* in *L. roboris* is a parsimonious inference from its *Buchnera* genome.

Bacteriome formation and symbiont distribution were also strikingly similar across both species (Figure 3, Table 2). *Buchnera* populations proliferated and cellularized as bacteriocytes, consistent with observations in the pea aphid and cedar bark aphid, *Cinara cedri* (20, 63, 65 66). *Serratia* cells, conversely, proliferated but remained as a cohesive symbiont mass (“secondary infection mass” in Klevenhausen [34]) during the early stages of embryonic development (Figure 4). As embryonic development progressed, the *Serratia* mass temporarily divided and elongated. The definitive symbiotic localization was then established as *Serratia* cells invaded the sheath cells. Initially, they invaded the sheath cells immediately adjacent to the *Serratia* mass. Subsequently, the entire *Serratia* infection mass ruptured, leading to the infection of all sheath cells. Notably, we observed the same phenomenon in *L. tropicalis* as previously reported in *L. roboris*, where a portion of the *Buchnera* population became engulfed within the *Serratia* mass, subsequently degenerated, and was expelled from the aphid (Figure 4I, I’). Although the functional significance of this phenomenon remains unclear, these consistent findings underscore that both aphid species follow nearly identical mechanisms for symbiont distribution, proliferation within the host, and arrangement within the symbiotic organs.

Furthermore, we compared the genomic features of *Buchnera* in *L. roboris* (*Buchnera* Lr) and *L. tropicalis* (*Buchnera* Lt) (Figure 5). First, FastANI was performed to assess genomic relatedness; the average nucleotide identity (ANI) value was 83.0, indicating an intragenus-level relationship despite evolutionary divergence. Reciprocal mapping between them indicated that many regions and their orders were evolutionarily conserved, with 119 of 141 ontologically query fragments ontologically matched (Figure 5A). Gene content and number were largely conserved between the two *Buchnera* genomes, with *Buchnera* Lr and Lt containing 380 and 386 CDSs, respectively. Notably, the histidine synthesis pathway was also not detected in *Buchnera* Lr, suggesting that its loss is not exclusively tied to symbiont replacement events. Our orthological analysis of the genomes of *Buchnera* Lr, Lt, and an outgroup, *Buchnera* APS, identified 349 orthogroups shared among all three strains and 20 groups exclusive to Lr and Lt. When comparing only *Buchnera* Lr and Lt, the analysis revealed that each orthogroup generally corresponded to a single gene in both *Lachnus Buchnera* genomes. A single exception was the 16S rRNA methyltransferase gene in *Buchnera* Lt, which was pseudogenized into two separate CDSs but remained within one orthogroup. A large number of shared genes were detected, with 369 genes in *Buchnera* Lr and 370 genes in *Buchnera* Lt belonging to the common 369 orthogroups. The remaining genes consisted of four genes detected only in *Buchnera* Lr and nine in *Buchnera* Lt (all hypothetical genes), as well as seven orthogroups (containing seven genes) shared exclusively with APS in each strain. These numbers fully account for the total CDS counts predicted by DFAST for both *Buchnera* Lr (380 CDSs) and Lt (386 CDSs). A notable difference in gene content was that *Buchnera* Lr lacked genes encoding elongation factor P (*efp*), leucine synthesis (*leuA–lepD*), and *repA*, whereas *Buchnera* Lt retained them. Conversely, *Buchnera* Lt had lost the tryptophan synthesis pathway genes, along with the heat shock protein *ibpB*, phosphofructokinase *pfkA*, and ribose-phosphate pyrophosphokinase *prs*. The presence of chromosomal tryptophan synthesis genes (*trpA*–*trpD*) in *Buchnera* Lr highlights its capacity to synthesize this amino acid, a function lacking in *Buchnera* Lt (Table 2). We assumed that both pLeu (*leuA*–*lepD*) and pTrp (*trpE* and *trpG*) plasmids are present in the *Buchnera* Lr genome but were not detected due to assembly limitations, which are common in short-read-only assemblies (Figure 5B and Supplementary Information Chapter 6). These differences in gene content, particularly the loss of the tryptophan pathway in *Buchnera* Lt, highlight the distinct evolutionary trajectories of the two *Lachnus* lineages and raise intriguing questions regarding the factors that facilitated these changes.

**Figure 5.**
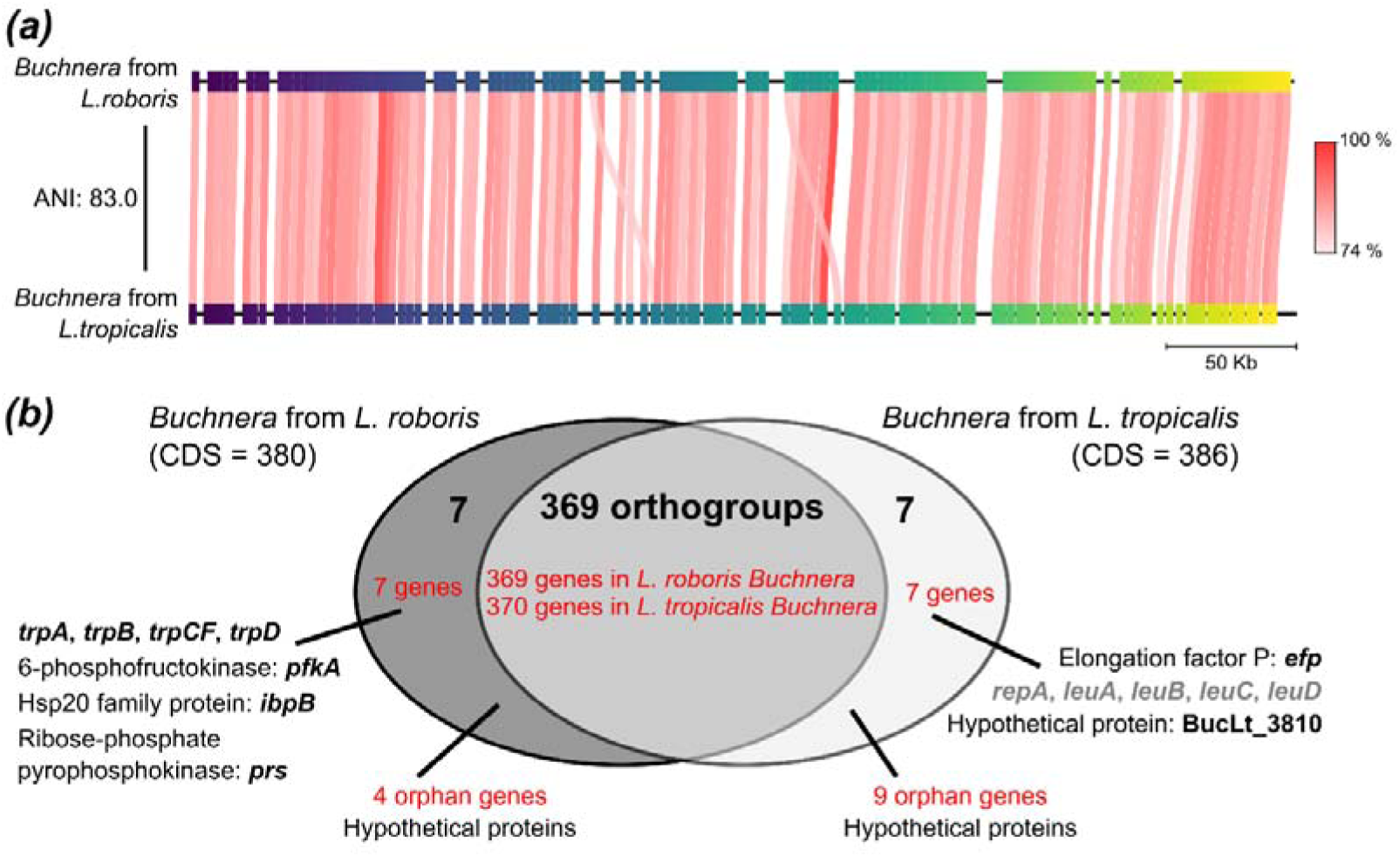
Genomic comparison of *Buchnera* from *Lachnus roboris* (*Buchnera* Lr) and *L. tropicalis* (*Buchnera* Lt). **(a)** FastANI analysis. Red bands represent reciprocal mapping between the *Buchnera* Lr (query) and *Buchnera* Lt (reference) genomes, indicating conserved regions. The average nucleotide identity (ANI) of 83.0% suggests intragenus-level similarity. **(b)** Orthologous gene analysis. Orthology was analyzed between *Buchnera* Lr and Lt, using *Buchnera* APS as the outgroup. The Venn diagram illustrates shared and unique gene sets. The large overlapping region contains 369 orthogroups, each corresponding to a single gene, accounting for the majority of genes common to both *Buchnera* in *Lachnus*. The only exception is the 16S rRNA methyltransferase gene in *Buchnera* Lt, which was pseudogenized into two CDSs but still grouped into one orthogroup. This results in 370 shared genes in *Buchnera* Lt and 369 in *Buchnera* Lr, out of total CDS counts of 386 and 380, respectively. The remaining genes are unique to each strain: 16 in *Buchnera* Lt and 11 in *Buchnera* Lr. Notably, *Buchnera* Lr retains chromosomal tryptophan synthesis genes (*trpA*–*trpD*), which are absent in *Buchnera* Lt. Genes on the pLeu plasmid (*leuA*–*leuD*) in *Buchnera* Lt are shown in gray. It is assumed that both pLeu and pTrp (*trpE* and *trpG*) plasmids are also present in the *Buchnera* Lr genome but were not detected presumably due to technical limitations (see Results and Supplementary Information Chapter 6).

## 4. Discussion

In this study, we investigated the evolutionary consequences of symbiont replacement within the aphid genus *Lachnus*, comparing the ancestral symbiotic system of *L. roboris* with the derived system of *L. tropicalis* (Figures S1, S2). *L. roboris*, a European species, harbors both *Buchnera* and *Serratia symbiotica* belonging to a more symbiotically advanced lineage (Clade B, Figure S2). In contrast, *L. tropicalis* and its sister Asian species have acquired a distinct *Serratia* lineage (Clade A, see Supplementary Information Chapter 1 for more details), which we hypothesized would replace the ancestral *Serratia* (29, 30). We characterized the symbiotic system of *L. tropicalis*, encompassing its ancient aphid symbiont *Buchnera* and the relatively recently acquired *Serratia symbiotica*. Microbiome and genomic analyses revealed that both symbionts were deeply metabolically integrated into the aphid host, indicating their obligate symbiotic nature. Genomic characterization suggests that *Serratia symbiotica* in *L. tropicalis* (*Serratia* Lt) is a recently integrated symbiont, as evidenced by its large genome size (ca. 3.0 Mb), abundant pseudogenes, neutral GC content, and a high number of rRNA and tRNA genes (Figure S4, Table S6). In contrast, the genome of *Buchnera* in *L. tropicalis* (*Buchnera* Lt) exhibited features similar to those of related *Buchnera* species (Table S5). We also performed histological observations to determine the tissue localization and vertical transmission modes of the symbionts in *L. tropicalis* (Figures 3 and 4; for details, see Figures S10 and S11, Supplementary Information Chapter 5).

Next, we compared the symbiotic systems of *L. tropicalis* and *L. roboris* by integrating existing knowledge of *L. roboris*, including microbiome analyses (26, 28, 29, 30), the sequence of the *Buchnera* genome (35), and detailed historical histological descriptions of symbiont localization and vertical transmission during embryogenesis (34). We found that, consistent with almost all other aphid species, *Buchnera* was housed in bacteriocytes in both aphid species, and *Serratia* was localized to the sheath cells in both species (Figure 3E, S9). Furthermore, no substantial differences were observed in the vertical transmission modes of *Buchnera* and *Serratia*, or in their allocation to the newly formed embryonic bacteriome between the two *Lachnus* species (Figure 4). These histological findings suggest that despite the distinct clades of *Serratia* (Clade B vs. Clade A) and their differing bacterial morphologies (cocci vs. bacilli), there are remarkably few histological differences between the ancestral and newly acquired *Serratia* symbionts. This implies that the new *Serratia* has effectively utilized, or “taken over,” the microniche of the ancestral *Serratia*. Although further research is needed to understand the mechanisms underlying this successful niche succession, the consistent presence of *Serratia* Clade A across diverse phylogenetic backgrounds (Figures S2, S6B) strongly suggests that its inherent adaptability and flexibility likely play crucial roles in its ability to take over these established symbiotic niches. Future histological studies, such as those in *Cinara* (where ancestral *Serratia* was often replaced by a different genus; 31, 67) or *Ceratovacuna* (where ancestral *Arsenophonus* symbionts were replaced by other lineages; 22, 27), could further elucidate the mechanisms of this niche succession.

Finally, to investigate the genomic consequences of symbiont replacement on primary and remaining symbionts, we compared the *Buchnera* genome sequences of *L. roboris* and *L. tropicalis* (Figure 5). We observed no major differences in genome size, CDS numbers, or syntenic relationships. Indeed, their ANI values were relatively high (>80%, indicative of intragenus-level relationships). However, *Buchnera* Lt had lost the tryptophan synthesis pathway genes on both the plasmid and chromosome, a function now compensated for by *Serratia* in *L. tropicalis*. In contrast, the histidine synthesis pathway (also covered by *Serratia* in *L. tropicalis*) was absent in *L. roboris Buchnera*. These results strongly suggest that, even for *Buchnera*, an ancient and deeply integrated symbiont, replacement events involving the acquisition of new symbionts with broader capacities can facilitate further genomic degeneration. Although the concept of symbiont replacement as a driver of symbiont genome degeneration has been proposed in previous studies (16, 17), our case provides a particularly compelling and well-documented example that uniquely demonstrates genomic consequences for the ancient symbiont *Buchnera*.

Our genomic analyses of *L. tropicalis* further elucidated the ongoing metabolic integration between its ancient (*Buchnera aphidicola*) and recently acquired (*Serratia symbiotica* Clade A) symbionts (Figures 2B, S7, S8, and Supplementary Information Chapter 4). Although *Serratia* broadly contributes to EAA and cofactor synthesis (such as histidine, phenylalanine, tryptophan, the initial steps of methionine, and nearly all B-vitamin pathways), a complementary, non-redundant relationship is clearly emerging. For instance, *Serratia* Lt abandoned parts of the methionine and leucine pathways, whereas the tryptophan pathway, originally maintained by *Buchnera*, was taken over by *Serratia*. This is noteworthy because other Clade A *Serratia* strains, such as CWBI-2.3 and 24.1, still possess the complete methionine and leucine pathways (57). This suggests that *Serratia* Lt lost these genes during its integration as a symbiont, likely because the *Buchnera* pathway, historically preserved in *L. roboris* and other Lachninae species (Figure 5, 32, 33), was a more efficient option. Similar dynamics were likely to develop for valine, leucine, and isoleucine. On the other hand, many redundant genes persisted in the arginine, threonine, lysine, and chorismate pathways. It is therefore reasonable to predict that, in future evolutionary processes, these overlapping pathways will be streamlined for efficiency. Further genomic analyses of *Buchnera* and *Serratia* in other *Lachnus* species, combined with gene expression studies of both symbionts and the aphid host, could reveal broader patterns of gene replacement by *Serratia* and retention by *Buchnera*. Such expression studies could also detect cases where redundant genes are retained but largely inactive, representing a pre-loss state and potentially indicative of genetic assimilation.

Symbiont replacement is a pivotal evolutionary event; however, its genomic antecedents and immediate drivers remain challenging to decipher (8). Although our research provides a unique example within the aphid genus *Lachnus*, a key limitation is the absence of genomic data for *Serratia* Clade B in *L. roboris*, leaving the precise state of ancestral *Serratia* prior to replacement unknown. Drawing parallels with other Sternorrhynchans, in which ancient symbionts such as *Hodgkinia* in cicadas (68) and certain *Sulcia* lineages in planthoppers show highly fragmented subgenomes (17, 69), it remains unclear whether *Serratia* Clade B in *Lachnus* exhibited similar genomic erosion before its replacement. Alternatively, factors such as host plant shifts or range expansion may have favored *Serratia* Clade A as relatively more efficient, leading to selective uptake (8). It is tempting to speculate that this successful replacement contributed to the adaptive radiation of *Lachnus*, particularly the rapid diversification of *L. tropicalis* and other *Serratia* Clade A-harboring species in East Asia (28, 29, 30, 51, 52). Further investigations of these ecological implications are crucial for a comprehensive understanding of the broader evolutionary impacts of replacement events. Future research should prioritize sequencing symbiont genomes from related *Lachnus* species and integrating life-history data to critically analyze the events surrounding this replacement.

## Author Contributions

T.N. contributed to conceptualization, data curation, formal analysis, funding acquisition, investigation, methodology, project administration, resources, validation, visualization, writing the original draft, and writing—review and editing. Y.K. contributed to data curation, investigation, methodology, resources, writing the original draft, and writing—review and editing. M.I. contributed to methodology, resources, and writing—review and editing. S.S. contributed to conceptualization, funding acquisition, investigation, project administration, writing the original draft, and writing—review and editing. All authors approved the final version of the manuscript and agreed to be accountable for this work.

## Acknowledgements

We thank Shunta Yorimoto for critical discussions and Wen Hsin-I and Katushi Yamaguchi for technical support with genomic library preparation and sequencing. We are also grateful to Jinyoung Choi, Filip Husnik, Ryuichi Koga, Takema Fukatsu, Akari, and Arisa Nozaki for sample collection and helpful advice. Finally, we thank Hiraku Yamada and the other members of the Laboratory of Evolutionary Genomics at the NIBB for their assistance with the experiments. Computational resources were provided by the Data Integration and Analysis Facility, NIBB.

## Generative AI disclosure

The authors verify and take full responsibility for the use of generative AI in the preparation of this manuscript. Generative AI (Google, 2025), specifically the Gemini model, was used as a writing assistant to improve grammar and sentence flow in parts of the manuscript. The scientific content, data analysis, and overall structure were developed solely by the authors.

## Data Accessibility

All datasets generated and analyzed in this study are publicly available. Raw sequencing reads, including 16S rRNA amplicon sequences (DRR709987–DRR709998), Nanopore long reads (DRR718948), and Illumina short reads (DRR718949), have been deposited in the DDBJ Sequence Read Archive (DRA) under BioProject ID PRJDB35790. The primary raw sequencing data are further associated with BioSample ID: SAMD01601843. The assembled genome sequences of *Buchnera aphidicola* isolate Lt-NIBB (BioSample ID: SAMD01609681, Accession numbers: AP043953 and AP043954) and *Serratia symbiotica* isolate Lt-NIBB (BioSample ID: SAMD01609682, Accession numbers: AP043955-AP043957) from *L. tropicalis* are also available in the DDBJ/GenBank/ENA database under BioProject ID PRJDB35790. All other relevant data supporting these findings are provided in this article and supplementary material.

