## Supplementary material for "Symbiont replacement and subsequent parallel genome erosion reshape a dual obligate symbiosis in the aphid *Lachnus tropicalis*": SI

*Author for correspondence:

ORCID:

https://orcid.org/0000-0003-4358-8118 (TN)

https://orcid.org/0000-0003-1061-3639 (YK)

https://orcid.org/0000-0003-4640-2323 (SS)

**This file includes;**

**Chapter 1: Symbiont replacement hypothesis in the genus *Lachnus***

**Chapter 2: Detailed microbiome analysis of *L. tropicalis***

**Chapter 3: Detailed genomic analysis of *L. tropicalis* symbionts**

**Chapter 4: Detailed metabolic insights into *L. tropicalis* symbionts**

**Chapter 5: Detailed protocols and observations for *L. tropicalis* symbionts**

**Chapter 6: A review of the symbiotic system in *L. roboris***

**Chapter 1: Symbiont replacement hypothesis in the genus *Lachnus***

**Background and methods for phylogenetic analysis**

To elucidate the evolutionary dynamics of “symbiont replacement” within the genus *Lachnus* (Burmeister, 1835), we reviewed previous studies describing the diversity of symbiotic bacteria in *Lachnus* (1, 2, 3, 4) and conducted phylogenetic analyses using available 16S ribosomal RNA (rRNA) gene sequences from *Buchnera aphidicola* and secondary symbionts (including *Serratia symbiotica*) obtained from aphid species within the tribe Lachnini, specifically *Lachnus* and its sister genera: *Pterochloroides*, *Maculolachnus*, and *Longistigma*. Detailed information on the sequences of *Buchnera* and secondary symbionts (*Serratia* and *Fukatsuia*) is provided in Tables S1 and S2, respectively.

To infer the host aphid phylogeny and facilitate the mapping of secondary symbiont associations, we constructed a *Buchnera* phylogenetic tree. Although concordance between *Buchnera* and the host phylogeny is generally accepted, despite some debate (5, 6), we prioritized *Buchnera* sequences for this purpose. Only *Buchnera* sequences detected in individual host samples that also contained secondary symbionts were included in the analysis (Table S1). Sequences were downloaded from the National Center for Biotechnology Information (NCBI) GenBank, manually inspected, and aligned using the ClustalX algorithm (7). *Longistigma*, identified as the most basal genus within Lachnini (8), was chosen as the outgroup for *Lachnus*, *Pterochloides*, and *Maculolachnus*, based on the established Lachnini phylogenetic relationships. The 16S rRNA sequence extracted from the *Buchnera* genome from *Lachnus tropicalis* (van der Goot, 1916) sequenced in this study (see main text) was also included. Phylogenetic analysis involving 23 nucleotide sequences was performed in MEGA X (9). Evolutionary history was inferred using the Maximum Likelihood method and the Hasegawa-Kishino-Yano model, with a discrete gamma distribution (five categories) to model evolutionary rate differences among sites. The final dataset comprised 1,286 nucleotide positions. Following the construction of this *Buchnera* phylogenetic tree, the types of secondary symbionts detected in the samples were manually mapped.

Within the genus *Lachnus*, *Serratia* is the only secondary symbiont observed to date. To clarify the phylogenetic relationship between intra-generic Clade A and Clade B *Serratia* lineages (2, 3), we constructed a phylogenetic tree using all available *Serratia symbiotica* 16S rRNA gene sequences (Table S2). Sequences were manually reviewed, and only those of sufficient length (>1,000 bases) were included in the analysis. The selected sequences were aligned using the ClustalX algorithm. As outgroups, we included 16S rDNA sequences from *Serratia marcescens (S. marcescens)*, *S. odorifera*, and *S. fonticola*. In addition, a *Serratia* symbiont detected in *Stomaphis* (SMLSS; *Serratia marcescens*-like secondary symbiont), closely related to *S. marcescens* (1, 2), was added as an outgroup. The 16S rRNA sequence extracted from the *L. tropicalis* *Serratia* genome sequenced in this study (see main text) was also included. Phylogenetic analysis of 48 nucleotide sequences was conducted in MEGA X. Evolutionary history was inferred using the Maximum Likelihood method and the Kimura 2-parameter model, with a discrete gamma distribution (five categories) to model evolutionary rate differences among sites. The final dataset comprised 1,216 nucleotide positions.

**Reconstruction of symbiont evolutionary scenarios in the tribe Lachnini**

A thorough review of the literature (1, 2, 3, 4), combined with phylogenetic analysis using deposited sequence data, revealed the placement of *Serratia symbiotica* as a presumed co-obligate symbiont within the tribe Lachnini, including the genus *Lachnus*. In *Longistigma liquidambarus*, *Serratia* Clade A was identified, whereas *Maculolachnus sumbacula* harbored both *Serratia* Clade A and *Fukatsuia*. In *Pterochloroides*, the sister group to *Lachnus*, *Serratia* Clade B was consistently found to as a co-obligate with *Buchnera* (with one reported case of *Serratia* Clade B infection lacking *Buchnera* sequence data [LT600341.1]) (Table S1). Within the genus *Lachnus*, *Serratia* Clade A was detected in derived, monophyletic species such as *L. tropicalis*, *L. takahashii*, *L. siniquercus*, *L. shiicola*, and *L. yunlongensis*, whereas *Serratia* Clade B was prevalent in basal species, including *L. roboris* (Linnaeus, 1758) and *L. quercihabitans* (Figure S1).

Sequence data for Lachnini remain less comprehensive than that for other Lachninae tribes (6, 8), leaving the exact timing of *Serratia* Clade B acquisition unresolved—whether it occurred in the common ancestor of *Pterochloroides* and *Lachnus* or independently in each genus (3, 11). Nevertheless, our updated phylogenetic analyses (Figures S1, S2; Table S1) allow for a conservative evolutionary scenario. Ancestrally, *Serratia* Clade B established a symbiotic association within *Lachnus* (Figure S1). Subsequently, in a subset of derived species (the *L. tropicalis* group), *Serratia* Clade B was replaced by Clade A. Notably, the acquisition of *Serratia* Clade A appears to have occurred only once during the speciation of the *L. tropicalis* group (Figure S2). This suggests a relatively recent replacement event—later than the origin of *Lachnus* but predating diversification within this group. Given its regional specificity (1, 2, 3), the *L. tropicalis* group may have undergone rapid speciation in East/Southeast Asia following the acquisition of *Serratia* Clade A.

Phylogenetic relationships of *Serratia* in *Pterochloroides* and *Lachnus* suggest that Clade B symbionts in these sister genera were acquired independently, as they do not form a monophyletic group (Figure S2). Conversely, the distribution of Clade A in *Longistigma* and *Maculolachnus* does not suggest monophyly, indicating that *Serratia* Clade A was unlikely widespread in the common ancestor of the Lachnini tribe. To fully resolve the dynamics of symbiont replacement across Lachnini, more extensive sampling and detailed analyses are needed.

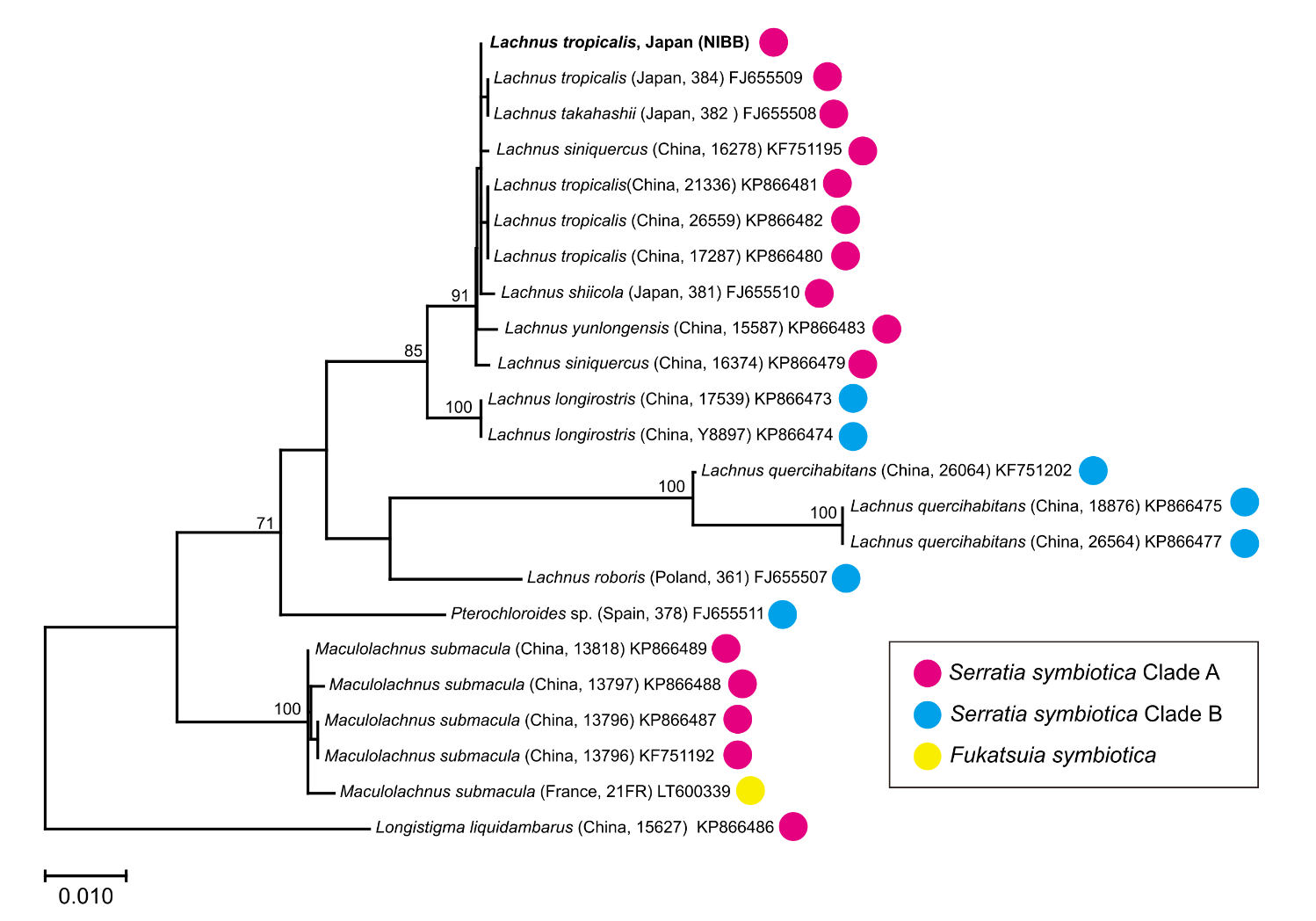
**Figure S1.** Phylogenetic relationship of *Buchnera*, reflecting the host aphid phylogeny, and their co-obligate partners in Lachnini. The tree was inferred using the Maximum Likelihood method based on 23 *Buchnera* 16S rRNA gene sequences from Lachnini aphids for which secondary symbionts were also detected. *Buchnera* from *Longistigma liquidambarus* was used as the outgroup. Bootstrap support values (≥70%, 500 replicates) are shown on the branches. *Lachnus tropicalis* (NIBB), newly sequenced in this study, is shown in **bold**. Aphid host species names are indicated at the tips of the tree, followed by country of collection and sample code (in parentheses), and the corresponding GenBank accession number. Branch lengths represent the number of substitutions per site. The final dataset included 1,286 nucleotide positions after removal of all gaps and missing data.

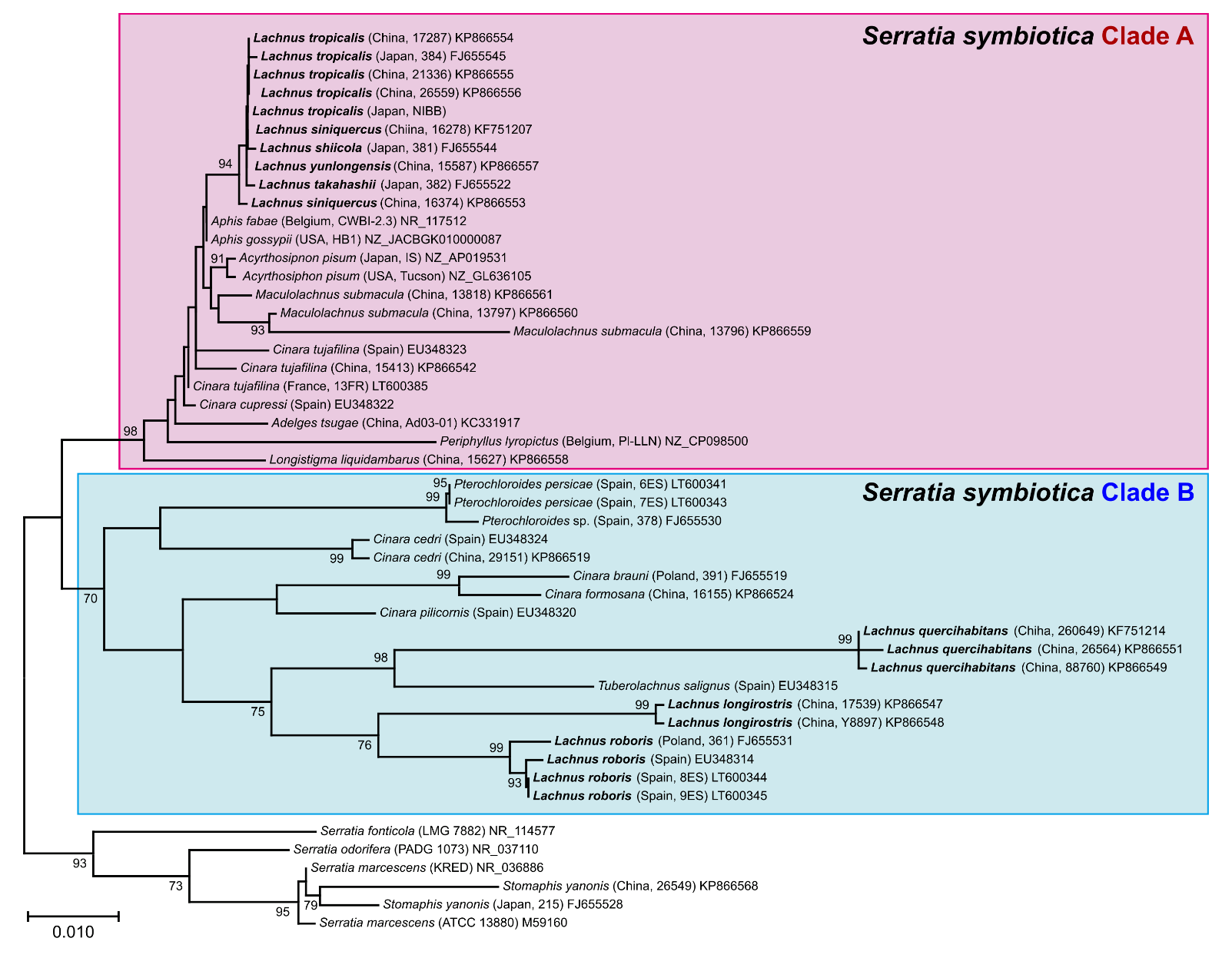
**Figure S2.** Phylogenetic relationships of *Serratia symbiotica* Clades A and B. The tree was inferred using the Maximum Likelihood method based on 42 *Serratia symbiotica* and six additional *Serratia* 16S rRNA gene sequences. *S. marcescens*, *S. fonticola*, *S. odorifera*, and the *Serratia* symbiont from *Stomaphis* (SMLSS) were used as the outgroup. Bootstrap support values (≥70%, 500 replicates) are shown on the branches. For *S. symbiotica*, host aphid species names are indicated at the tree tips, followed by country of collection and sample code (in parentheses), and the corresponding GenBank accession number. *Lachnus tropicalis* (NIBB), newly sequenced in this study, is highlighted in **bold**. The tree is drawn to scale, with branch lengths representing the number of substitutions per site. The final dataset included 1,216 nucleotide positions after removal of all gaps and missing data.

**Table S1.** List of symbiotic bacteria detected in *Lachnus* and related genera, including 16S rRNA sequences used for *Buchnera* phylogenetic analysis and associated symbiont information (related to Figure S1).

| ID | Host aphid | Accession no. of *Buchnera* | Sequence length (bp) | Associated symbiont | Accession no. of another symbiont | Isolate/voucher code | Collection locality |
| --- | --- | --- | --- | --- | --- | --- | --- |
| 1 | *Lachnus roboris* | **FJ655507.1** | 1519 | *Serratia symbiotica* (Clade B) | FJ655531.1 | *Lachnus roboris* isolate 361 | Poland |
| 2 | *Lachnus roboris* | AJ296756.1 | 1506 | - | - | - | Unknown |
| 3 | *Lachnus roboris* | - | - | *Serratia symbiotica* (Clade B) | EU348314 | - | Spain |
| 4 | *Lachnus roboris* | - | - | *Serratia symbiotica* (Clade B) | LT600344.1 | 8ES | Spain |
| 5 | *Lachnus roboris* | - | - | *Serratia symbiotica* (Clade B) | LT600345.1 | 9ES | Spain |
| 6 | *Lachnus quercihabitans* | **KF751202.1** | 1504 | *Serratia symbiotica* (Clade B) | KF751214.1 | isolate 26064 | China |
| 7 | *Lachnus quercihabitans* | **KP866475.1** | 1348 | *Serratia symbiotica* (Clade B) | KP866549.1 | voucher 18876 | China |
| 8 | *Lachnus quercihabitans* | **KP866477.1** | 1348 | *Serratia symbiotica* (Clade B) | KP866551.1 | voucher 26564 | China |
| 9 | *Lachnus longirostris* | **KP866473.1** | 1347 | *Serratia symbiotica* (Clade B) | KP866547.1 | voucher 17539 | China |
| 10 | *Lachnus longirostris* | **KP866474.1** | 1347 | *Serratia symbiotica* (Clade B) | KP866548.1 | voucher Y8897 | China |
| 11 | *Lachnus tropicalis* | **AP043953** | 1540 | *Serratia symbiotica* (Clade A) | AP043955 | This study | Japan |
| 12 | *Lachnus tropicalis* | **FJ655509.1** | 1326 | *Serratia symbiotica* (Clade A) | FJ655545.1 | *Lachnus tropicalis* isolate 384 | Japan |
| 13 | *Lachnus tropicalis* | **KP866480.1** | 1347 | *Serratia symbiotica* (Clade A) | KP866554.1 | voucher 17287 | China |
| 14 | *Lachnus tropicalis* | **KP866481.1** | 1347 | *Serratia symbiotica* (Clade A) | KP866555.1 | voucher 21336 | China |
| 15 | *Lachnus tropicalis* | **KP866482.1** | 1347 | *Serratia symbiotica* (Clade A) | KP866556.1 | voucher 26559 | China |
| 16 | *Lachnus tropicalis* | JX998110.1 | 1392 | - | - | ZMIOZ 14528 | China |
| 17 | *Lachnus siniquercus* | **KP866479.1** | 1347 | *Serratia symbiotica* (Clade A) | KP866553.1 | voucher 16374 | China |
| 18 | *Lachnus siniquercus* | **KF751195.1** | 1500 | *Serratia symbiotica* (Clade A) | KF751207.1 | isolate 16278 | China |
| 19 | *Lachnus shiicola* | **FJ655510.1** | 1319 | *Serratia symbiotica* (Clade A) | FJ655544.1 | *Lachnus shiicola* isolate 381 | Japan |
| 20 | *Lachnus takahashii* | **FJ655508.1** | 1346 | *Serratia symbiotica* (Clade A) | FJ655522.1 | *Lachnus takahashii* isolate 382 | Japan |
| 21 | *Lachnus yunlongensis* | **KP866483.1** | 1347 | *Serratia symbiotica* (Clade A) | KP866557.1 | voucher 15587 | China |
| 22 | *Pterochloroides persicae* | - | *-* | *Serratia symbiotica* (Clade A) | LT600341.1 | 6ES | Spain |
| 23 | *Pterochloroides persicae* | LT600342.1 | 636 | *Serratia symbiotica* (Clade B) | LT600343.1 | 7ES | Spain |
| 24 | *Pterochloroides sp.* | **FJ655511.1** | 1523 | *Serratia symbiotica* (Clade B) | FJ655530.1 | *Pterochloroides* sp. isolate 378 | Spain |
| 25 | *Maculolachnus submacula* | **KF751192.1** | 1508 | *Serratia symbiotica* (Clade A) | KF751204.1 | isolate 13796 | China |
| 26 | *Maculolachnus submacula* | **KP866487.1** | 1347 | *Serratia symbiotica* (Clade A) | KP866559.1 | voucher 13796 | - |
| 27 | *Maculolachnus submacula* | **KP866488.1** | 1347 | *Serratia symbiotica* (Clade A) | KP866560.1 | voucher 13797 | - |
| 28 | *Maculolachnus submacula* | **KP866489.1** | 1347 | *Serratia symbiotica* (Clade A) | KP866561.1 | voucher 13818 | - |
| 29 | *Maculolachnus submacula* | **LT600339.1** | 1450 | *Fukatsuia symbiotica* | LT600340.1 | 21FR | France |
| 30 | *Maculolachnus submacula* | AJ296755.1 | 1508 | - | - | - | - |
| 31 | *Maculolachnus submacula* | - | *-* | *Fukatsuia symbiotica* | EU348312 | - | Spain |
| 32 | *Longistigma liquidambarus* | **KP866486.1** | 1354 | *Serratia symbiotica* | KP866558.1 | voucher 15627 | China |

Note: Accession numbers for *Buchnera* shown in **bold** were used for phylogenetic analysis. These represent instances where both *Buchnera* and its partner symbiont were co-detected from a single sample. LT600342 was excluded due to its limited sequence length.

**Table S2.** List of 16S rRNA sequences of *Serratia symbiotica* and related species, used for phylogenetic analysis (related to figure S2).

| ID | Accession no. | Species name | Host aphid | Isolate/voucher code | Collection locality |
| --- | --- | --- | --- | --- | --- |
| 1 | NR_117512.1 | *Serratia symbiotica* (Clade A) | *Aphis fabae* | CWBI-2.3 | Belgium |
| 2 | NZ_JACBGK010000087.1 | *Serratia symbiotica* (Clade A) | *Aphis gossypii* | HB1 87 | USA |
| 3 | NZ_AP019531.1 | *Serratia symbiotica* (Clade A) | *Acyrthosipnon pisum* | Strain IS | Japan |
| 4 | NZ_GL636105.1 | *Serratia symbiotica* (Clade A) | *Acyrthosiphon pisum* | Strain Tucson | USA |
| 5 | EU348322.1 | *Serratia symbiotica* (Clade A) | *Cinara cupressi* | - | Spain |
| 6 | KP866542.1 | *Serratia symbiotica* (Clade A) | *Cinara tujafilina* | voucher 15413 | China |
| 7 | LT600385.1 | *Serratia symbiotica* (Clade A) | *Cinara tujafilina* | isolate 13FR | France |
| 8 | EU348323.1 | *Serratia symbiotica* (Clade A) | *Cinara tujafilina* | - | Spain |
| 9 | KP866560.1 | *Serratia symbiotica* (Clade A) | *Maculolachnus submacula* | voucher 13797 | China |
| 10 | KP866561.1 | *Serratia symbiotica* (Clade A) | *Maculolachnus submacula* | voucher 13818 | China |
| 11 | KP866559.1 | *Serratia symbiotica* (Clade A) | *Maculolachnus submacula* | voucher 13796 | China |
| 12 | KP866558.1 | *Serratia symbiotica* (Clade A) | *Longistigma liquidambarus* | voucher 15627 | China |
| 13 | AP043955 | *Serratia symbiotica* (Clade A) | *Lachnus tropicalis* | This study | Japan |
| 14 | KP866554.1 | *Serratia symbiotica* (Clade A) | *Lachnus tropicalis* | voucher 17287 | China |
| 15 | KP866555.1 | *Serratia symbiotica* (Clade A) | *Lachnus tropicalis* | voucher 21336 | China |
| 16 | KP866556.1 | *Serratia symbiotica* (Clade A) | *Lachnus tropicalis* | voucher 26559 | China |
| 17 | FJ655545.1 | *Serratia symbiotica* (Clade A) | *Lachnus tropicalis* | isolate 384 | Japan |
| 18 | KP866557.1 | *Serratia symbiotica* (Clade A) | *Lachnus yunlongensis* | voucher 15587 | China |
| 19 | FJ655544.1 | *Serratia symbiotica* (Clade A) | *Lachnus shiicola* | isolate 381 | Japan |
| 20 | KF751207.1 | *Serratia symbiotica* (Clade A) | *Lachnus siniquercus* | isolate 16278 | China |
| 21 | KP866553.1 | *Serratia symbiotica* (Clade A) | *Lachnus siniquercus* | voucher 16374 | China |
| 22 | FJ655522.1 | *Serratia symbiotica* (Clade A) | *Lachnus takahashii* | isolate 382 | Japan |
| 23 | Z_CP098500.1 | *Serratia symbiotica* (Clade A) | *Periphyllus lyropictus* | Strain Pl-LLN | Belgium |
| 24 | KC331917.1 | *Serratia symbiotica* (Clade A) | *Adelges tsugae* (adelgid) | Ad03-01_GSB | China |
| 25 | EU348324 | *Serratia symbiotica* (Clade B) | *Cinara cedri* | Spain 16S | Spain |
| 26 | KP866519.1 | *Serratia symbiotica* (Clade B) | *Cinara cedri* | voucher 29151 | China |
| 27 | FJ655519.1 | *Serratia symbiotica* (Clade B) | *Cinara brauni* | isolate 391 | Poland |
| 28 | KP866524.1 | *Serratia symbiotica* (Clade B) | *Cinara formosana* | voucher 16155 | China |
| 29 | EU348320.1 | *Serratia symbiotica* (Clade B) | *Cinara pilicornis* | - | Spain |
| 30 | LT600341.1 | *Serratia symbiotica* (Clade B) | *Pterochloroides persicae* | isolate 6ES | Spain |
| 31 | LT600343.1 | *Serratia symbiotica* (Clade B) | *Pterochloroides persicae* | isolate 7ES | Spain |
| 32 | FJ655530.1 | *Serratia symbiotica* (Clade B) | *Pterochloroides* sp. | isolate 378 | Spain |
| 33 | EU348315.1 | *Serratia symbiotica* (Clade B) | *Tuberolachnus salignus* | - | Spain |
| 34 | LT600344.1 | *Serratia symbiotica* (Clade B) | *Lachnus roboris* | isolate 8ES | China |
| 35 | LT600345.1 | *Serratia symbiotica* (Clade B) | *Lachnus roboris* | isolate 9ES | China |
| 36 | FJ655531.1 | *Serratia symbiotica* (Clade B) | *Lachnus roboris* | isolate 361 | Poland |
| 37 | EU348314.1 | *Serratia symbiotica* (Clade B) | *Lachnus roboris* | - | Spain |
| 38 | KP866547.1 | *Serratia symbiotica* (Clade B) | *Lachnus longirostris* | voucher 17539 | China |
| 39 | KP866548.1 | *Serratia symbiotica* (Clade B) | *Lachnus longirostris* | voucher Y8897 | China |
| 40 | KP866549.1 | *Serratia symbiotica* (Clade B) | *Lachnus quercihabitans* | voucher 18876 | China |
| 41 | KF751214.1 | *Serratia symbiotica* (Clade B) | *Lachnus quercihabitans* | isolate 26064 | China |
| 42 | KP866551.1 | *Serratia symbiotica* (Clade B) | *Lachnus quercihabitans* | voucher 26564 | China |
| 43 | KP866568.1 | *Serratia marcescens* | *Stomaphis yanonis* | voucher 26549 | China |
| 44 | FJ655528.1 | *Serratia marcescens* | *Stomaphis yanonis* | - | Japan |
| 45 | M59160.1 | *Serratia marcescens* | - | ATCC 13880 | - |
| 46 | NR_036886.1 | *Serratia marcescens* | - | strain KRED | - |
| 47 | NR_037110.1 | *Serratia odorifera* | - | strain PADG 1073 | - |
| 48 | NR_114577.1 | *Serratia fonticola* | - | strain LMG 7882 | - |

**Chapter 2: Detailed microbiome analysis of *L. tropicalis***

**Detailed extraction and sequencing methods**

Total DNA was extracted as follows: single individuals preserved in 99.5% ethanol were rinsed with 70% ethanol, air-dried, and briefly rinsed with Buffer A (10 mM Tris [pH 8.0], 1 mM EDTA, and 25 mM NaCl). Each sample was placed in 100 μL of Buffer A containing 1 μL of proteinase K (400 μg/mL) and completely homogenized using BioMasher II (Nippi, Japan). The homogenates were incubated at 37 °C for 1 h, followed by heating at 98°C for 2 min.

The extracted DNA was used to construct sequencing libraries according to Illumina’s “16S Metagenomic Sequencing Library Preparation Guide (15044223 B JPN).” The V3/V4 region (ca. 460 bp) of the bacterial 16S rRNA gene was amplified using the primers 16S AmpF_IL and 16S AmpR_IL. Each 10 μL polymerase chain reaction (PCR) contained 2 μL DNA template, 1 μL of each primer (2 µM), 5 μL of 2x KAPA HiFi HotStart ReadyMix (KAPA Biosystems, USA), and 2 μL ultrapure water. The PCR program was: 95°C for 3 min; 25 cycles of 95°C for 30 s, 55°C for 30 s, and 72°C for 30 s; and a final extension at 72°C for 5 min. PCR products were purified using AMPure XP beads (Beckman Coulter, USA). Indexing was performed with the Nextera XT Index Kit (Nextera DNA UD Index Set B; Illumina). The quality of purified libraries was assessed using a TapeStation D1000 (Agilent, USA). Pooled libraries were sequenced on an Illumina MiSeq platform, generating 250 bp paired-end reads (Table S3).

| # | Insect stage (viviparous female) | Locality | No. of raw read^*^ | DRR Run No.  (PRJDB35790) | Detected bacteria^†^ |
| --- | --- | --- | --- | --- | --- |
| 1 | Adult female (apterous) | Okazaki, Aichi | 62,568 | DRR709987 | ***Buchnera***, ***Serratia*** |
| 2 | Adult female (winged) | Okazaki, Aichi | 25,104 | DRR709988 | ***Buchnera***, ***Serratia***, *Wolbachia, Rickettsia* |
| 3 | Adult female (winged) | Okazaki, Aichi | 22,953 | DRR709989 | ***Buchnera***, ***Serratia***, *Wolbachia, Rickettsia* |
| 4 | Adult female (winged) | Okazaki, Aichi | 17,862 | DRR709990 | ***Buchnera***, ***Serratia***, *Wolbachia, Rickettsia* |
| 5 | Adult female (apterous) | Tsuruoka, Yamagata | 21,575 | DRR709991 | ***Buchnera***, ***Serratia***, *Wolbachia, Rickettsia* |
| 6 | Adult female (apterous) | Tsuruoka, Yamagata | 24,064 | DRR709992 | ***Buchnera***, ***Serratia***, *Wolbachia, Rickettsia* |
| 7 | Late-stage nymph | Tsuruoka, Yamagata | 23,750 | DRR709993 | ***Buchnera***, ***Serratia***, *Wolbachia, Rickettsia* |
| 8 | Late-stage nymph | Tsuruoka, Yamagata | 23,714 | DRR709994 | ***Buchnera***, ***Serratia***, *Wolbachia, Rickettsia* |
| 9 | Adult female (apterous) | Tsukuba, Ibaraki | 20,633 | DRR709995 | ***Buchnera***, ***Serratia***, *Wolbachia, Rickettsia* |
| 10 | Adult female (apterous) | Tsukuba, Ibaraki | 40,067 | DRR709996 | ***Buchnera***, ***Serratia***, *Wolbachia, Rickettsia* |
| 11 | Late-stage nymph | Tsukuba, Ibaraki | 19,799 | DRR709997 | ***Buchnera***, ***Serratia***, *Wolbachia, Rickettsia* |
| 12 | Late-stage nymph | Tsukuba, Ibaraki | 20,910 | DRR709998 | ***Buchnera***, ***Serratia***, *Wolbachia, Rickettsia* |

**Table S3.** Detailed information on amplicon sequencing

^*^ 250 bp of paired-end reads generated by Illumina MiSeq.

^†^Bacterial taxa detected with 100 or more reads as ASV in at least one sample are listed. Those accounting for more than 1% of the total reads per sample are shown in **bold**.

**Chapter 3: Detailed genomic analysis of *Lachnus tropicalis* symbionts**

**Detailed methods for Nanopore and Illumina genomic library preparation**

For Nanopore long-read sequencing, we prepared high molecular weight DNA from five fresh young adults collected from the NIBB campus using frozen, powdery QIAGEN G2 buffer (Qiagen, Japan) and a QIAGEN Genomic-tip 20/G column (details in 12). The quantity of extracted DNA was measured using a Qubit dsDNA HS Assay Kit (Thermo Fisher Scientific, Waltham, MA, USA) and a Qubit 2.0 Fluorometer (Thermo Fisher Scientific). The integrity of the genomic DNA was assessed by pulsed-field gel electrophoresis using a CHEF Mapper (Bio-Rad). Using this DNA sample, which contained genomes derived from both aphids and symbionts, a Nanopore sequencing library was prepared with the SQL-LSK110 Ligation Sequencing Kit (Oxford Nanopore Technologies, UK) according to the manufacturer’s instructions and sequenced using an R9.4.1 flow cell on the GridION system. Reads were base-called using “super accuracy mode” implemented in Guppy (version 5.0.7).

For Illumina short-read sequencing, genomic DNA was extracted from five fresh young adults collected from the NIBB campus using the DNeasy Blood & Tissue Kit (Qiagen). The extracted DNA was fragmented into 200–500 bp fragments using a Covaris Focused-ultrasonicator M220. A library for whole-genome sequencing was prepared with the TruSeq DNA PCR-Free Library Prep Kit (Illumina) according to the manufacturer’s protocol. Library quality was validated using TapeStation HS D5000 (Agilent Technologies, USA). Library concentrations were quantified using an Applied Biosystems 7500 Real-Time PCR System (Applied Biosystems, Japan). Sequencing was performed on an Illumina HiSeq X Ten platform (Illumina) at Macrogen Japan (Tokyo, Japan) using a 2 × 150 bp paired-end sequencing protocol.

**Detailed strategies of genome assembly using long- and short-read sequences**

Initially, raw Nanopore reads were used to assemble the symbiont genome backbones. Following the removal of relatively short sequences (<5000 bp) using Seqkit (v0.8.1) (13), the remaining 386,257 reads underwent metagenomic assembly with Minimap2 (version 2.17-r941) (14) and Miniasm (version 0.4-r179) (15) (hereafter referred to as the initial long-read assembly). All 275 contigs from this assembly were subjected to a local BLAST search against the reference genomes of *Buchnera aphidicola* APS (GCF_000009605), *Serratia symbiotica* CWBI-2.3 (GCF_000821185), and *Serratia symbiotica* IS (GCF_008370165). This analysis identified a circularized *Buchnera* contig (chromosome: approximately 0.4 Mb) and a non-circularized *Serratia* contig (chromosome: approximately 2.9 Mb) from the initial long-read assembly. Mapping the original long reads to the *Buchnera* and *Serratia* contigs using Minimap2 and examining depth of coverage with SAMtools (version 1.16) (16) revealed an average depth of 336.8× for the *Buchnera* contig and 41.6× for the *Serratia* contig. The *Buchnera* contig was polished with Racon (version 1.4.20) (17) and Medaka (version 1.4.1, https://github.com/nanoporetech/medaka) to obtain consensus sequences.

Next, based on the mapping results of the 2.9 Mb *Serratia* contig, raw long reads were extracted. Primary mapped reads were selected using SAMtools (version 1.16) and Seqkit (v0.8.1). These 23,831 selected reads were then used for another round of *Serratia* genome assembly with the same parameters as the initial long-read assembly. Local BLAST analysis of this second-round assembly revealed a circularized *Serratia* contig (2.9 Mb). Similar to the *Buchnera* contig, polishing and consensus calling were performed using Racon (version 1.4.20) and Medaka (version 1.4.1).

Given that our local BLAST search of the initial long-read assembly did not detect plasmids—often maintained by both *Buchnera* (typically leucine and tryptophan plasmids, pLeu and pTrp, respectively) (18; Table S5) and *Serratia* *symbiotica* (up to two plasmids: one in IS [19], two in CWBI-2.3 [20]; Table S6)—we extracted 39 circular contigs, excluding the *Buchnera* chromosome. Subsequent web-based BLAST searches (blastn) identified three plasmids based on their best hits: *Buchnera* [*Cinara spledens*] plasmid pLeu [LR217723.1], which we designated *Buchnera* pLeu; *Serratia* IS plasmid pSsyis1 [NZ_AP019532.1], designated pSsLt-1; and *Serratia* CWBI-2.3, plasmid pSsAf2.3-1 [NZ_CP050856], designated pSsLt-2. Following the same polishing procedure used for the chromosomes, each plasmid was polished with Racon (version 1.4.20), and consensus sequences were generated with Medaka (version 1.4.1). In summary, the long-read assembly yielded five circular contigs representing the *Buchnera* chromosome (437,441 bp), *Buchnera* pLeu (6,456 bp), the *Serratia* chromosome (2,887,204 bp), pSsLt-1 (42,855 bp), and pSsLt-2 (58,491 bp).

To confirm the assembled contigs, Illumina raw reads were mapped using BWA (version 0.7.17-r1188) (21) to all contigs from the initial long-read assembly that had been polished with Racon (version 1.4.20) and Medaka (version 1.4.1). Adapter trimming and quality filtering were performed on raw Illumina paired-end reads using Trim Galore (version 0.6.10) (22). A blob plot was created based on sequencing depth with BlobTools (version 1.1.1) (23), which allowed us to identify contigs corresponding to the chromosomes of *Buchnera* and *Serratia* as well as their plasmids (Figure S3). Circularity was confirmed by mapping Nanopore reads to symbiont genome sequences using Minimap2 and checking for overlapping reads at both ends.

For final error correction with Illumina short reads, the cleaned Illumina reads (n = 203,128,459) were mapped to the assembled genomes and plasmids using BWA (version 0.7.17-r1188). Assembly polishing was performed with Pilon (version 1.24) (24), and this step was repeated three times.

**Clusters of orthologous genes (COG) analysis, gene repertoire, and comparisons with related genomes**

DDBJ Fast Annotation and Submission Tool (DFAST) annotations also included COG category tags (25). To characterize the genome content of *Buchnera* and *Serratia* symbionts in *L. tropicalis*, we compared their gene content with that of related bacteria using COG category tags. Genomes of the endosymbiont *Buchnera aphidicola* strain APS (GCF_000009605) and strain BCc (GCF_000090965.1), along with BCc’s tryptophan plasmid (EU660486.1), were retrieved from GenBank (accessed October 2024). Notably, the deposited genome of *Buchnera aphidicola* strain BCc contains a leucine plasmid but lacks a tryptophan plasmid (26), although the presence of the tryptophan has been confirmed (27). Similarly, genomes of *Serratia symbiotica* were obtained (strain CWBI-2.3, GCF_000821185.2; strain IS, GCF_008370165.1; strain ‘*Cinara cedri*,’ GCF_000238975.1). The genomes were processed using DFAST and compared with those of *Buchnera* and *Serratia* in *L. tropicalis*. The number of COG category tags was recorded, distinguishing between pseudogenes and intact genes.

**Taxon sampling for the phylogenomic analysis of *L. tropicalis,* *Serratia,* and *Buchnera***

For the *Buchnera* genome analysis, 53 genomes were included, comprising 50 *Buchnera* genomes (including one sequenced in this study) and three outgroup species: *Escherichia coli*, *Ishikawaella capsulata*, and *Wiggleworthia glossinidia*. This analysis was based on a concatenated alignment of 145 genes, yielding a total length of 29,425 amino acids, generated with GToTree. The final maximum-likelihood tree search with IQ-TREE was completed after 102 iterations, and nodal support was assessed with 1,000 ultrafast bootstrap replicates. For the *Serratia* genome analysis, 29 genomes were used, including 25 *Serratia symbiotica* genomes (one sequenced in this study) and four outgroup *Serratia* lineages (*Serratia ficaria*, *S. entomophila*, and two *S. marcescens* genomes). A total of 172 target genes were used, producing a concatenated alignment of 38,172 amino acids. Similar to the *Buchnera* analysis, the IQ-TREE run completed its final maximum-likelihood tree search after 102 iterations, and 1,000 ultrafast bootstrap replicates were generated to evaluate nodal support.

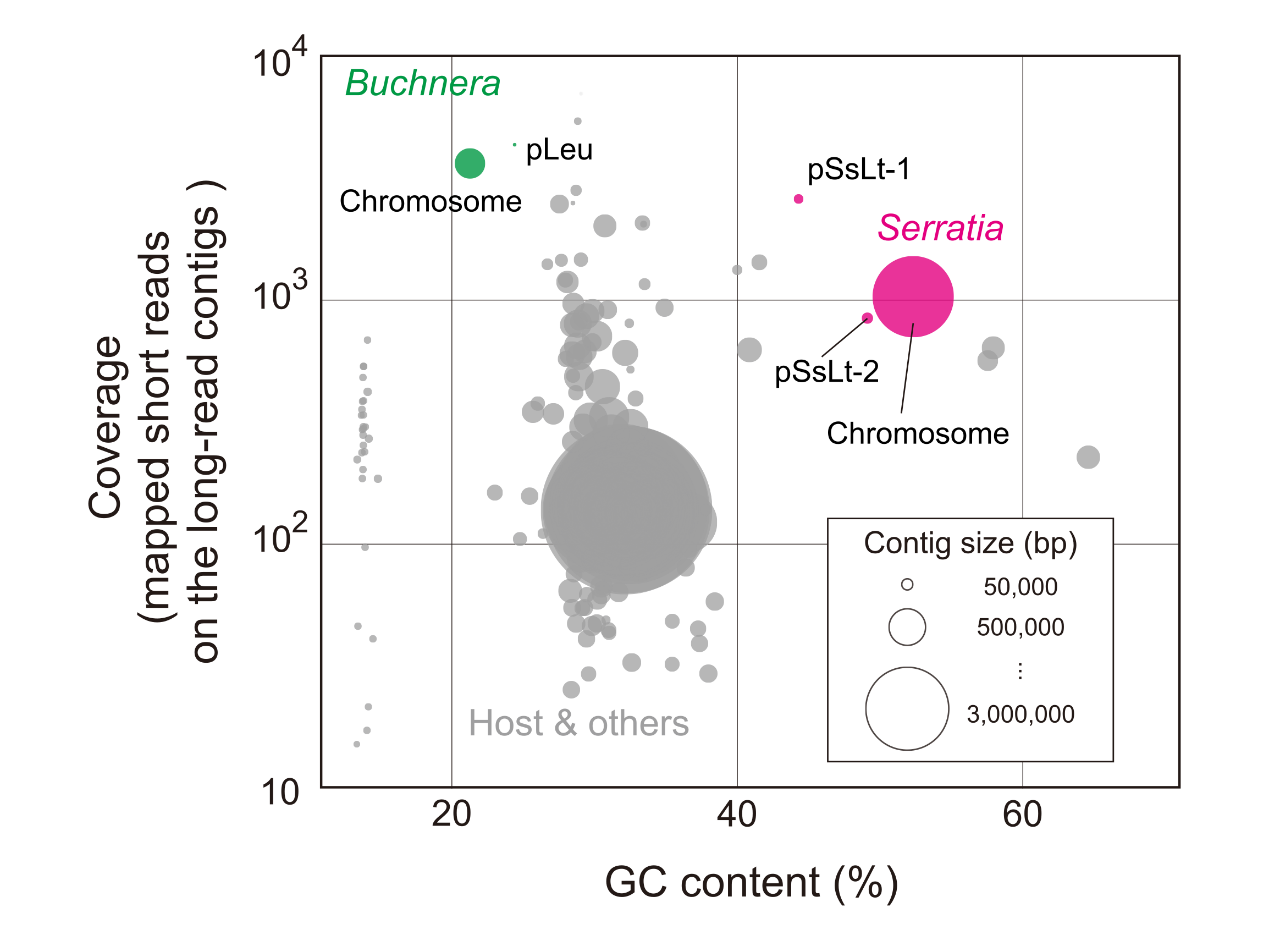
**Figure S3.** **Blobplot: GC-coverage plot for metagenomic assembly.** The plot displays GC content versus coverage for contigs obtained from the metagenomic assembly. Coverage was calculated by mapping Illumina short reads to the consensus assembly generated using Nanopore reads. Each blob represents a contig in the assembly. *Buchnera* contigs are colored green, and *Serratia* contigs are colored magenta, with chromosomes and plasmids identified based on local BLAST analysis. “Host & others” indicates contigs that did not match *Buchnera* or *Serratia*. The size of each blob corresponds to the contig length. All assembled contigs showed high coverage (>10^3^), indicating the high quality of the final genomes obtained in this study.

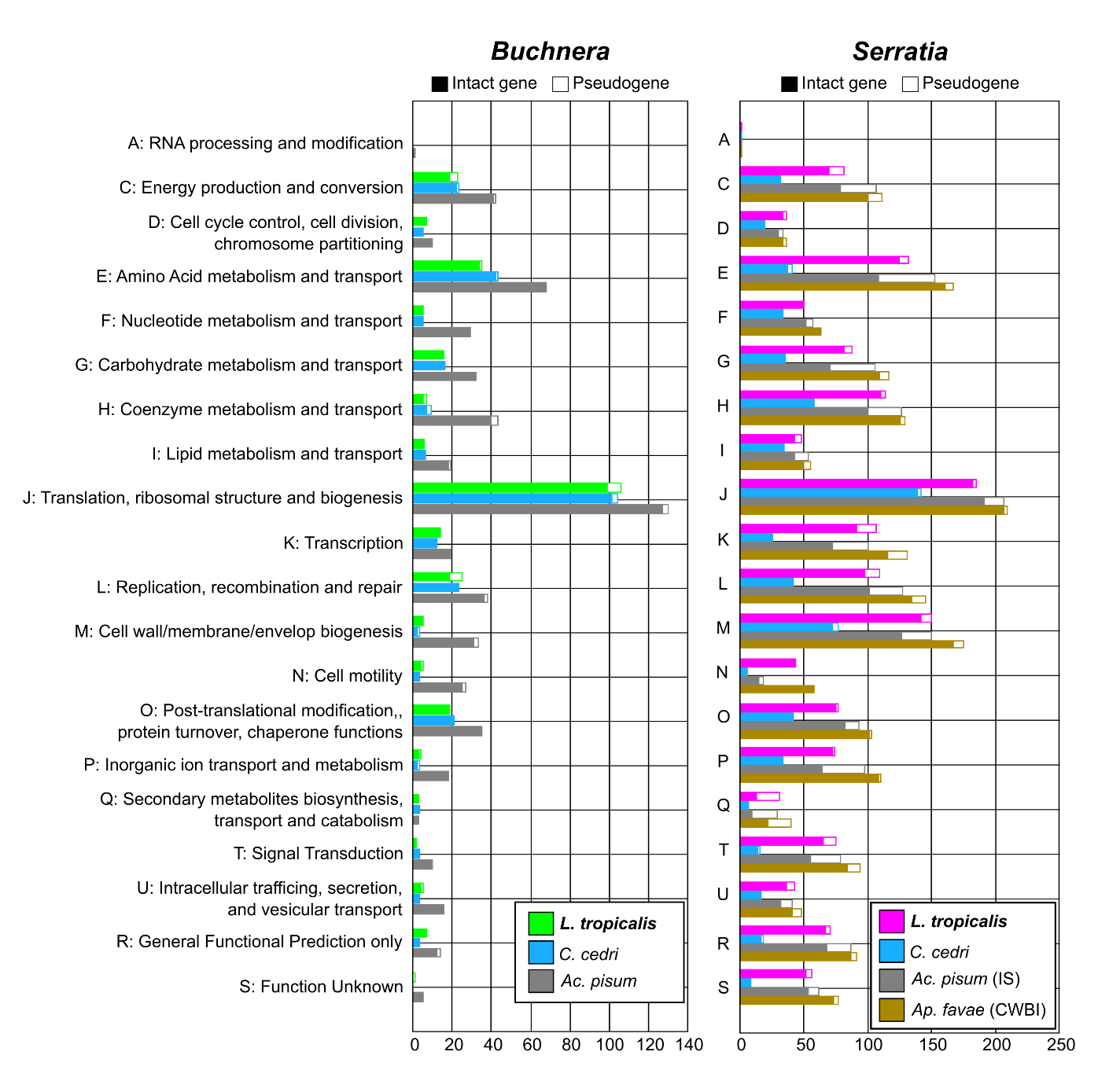
**Figure S4. COG classification of protein-coding genes in *Buchnera* and *Serratia* genomes.** Gene content classification based on COG categories, derived from DFAST annotation, was visualized for *Buchnera* and *Serratia* genomes. For *Buchnera* in *Lachnus tropicalis* (*Buchnera* Lt), its COG profile was compared with those of *Buchnera* in *Cinara cedri* (GCA_000090965.1) and *Acyrthosiphon pisum* (GCA_000009605.1). For *Serratia* in *L. tropicalis* (*Serratia* Lt), comparisons were made with *Serratia* from *Cinara cedri* (GCA_000238975.1), *Acyrthosiphon pisum* (GCA_000009605.1), and *Aphis fabae* (GCA_000821185.2). Pseudogene information, also obtained from DFAST results, was integrated into this analysis.

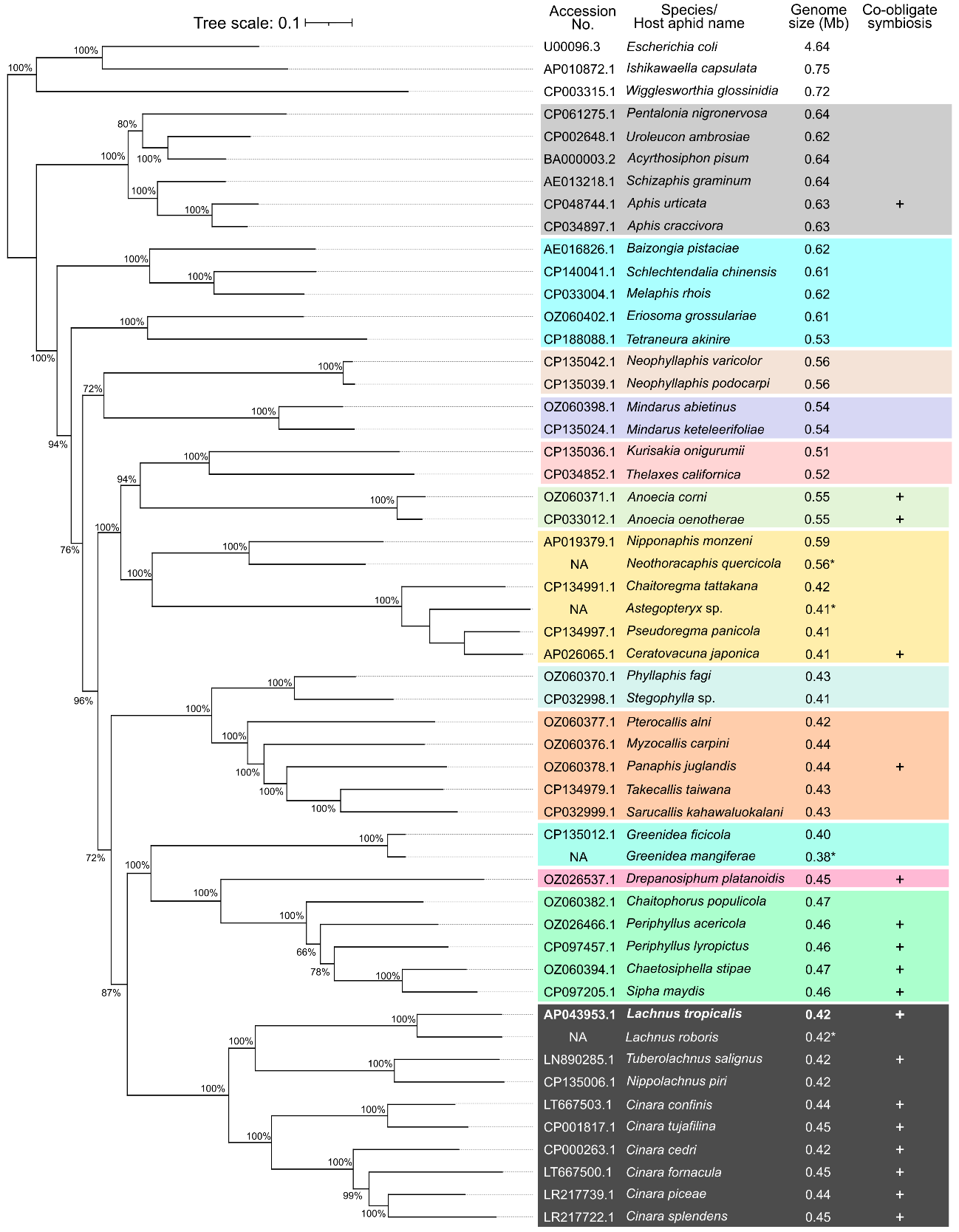
**Figure S5.** Phylogenomic tree estimated using IQ-TREE based on a concatenated alignment of 145 core orthologous genes (29,425 amino acids) generated with GToTree. The final maximum-likelihood tree search was completed after 102 iterations. Nodal support was assessed with 1,000 ultrafast bootstrap replicates; bootstrap values ≥70 are shown. Genome sizes marked with an asterisk (*) indicate draft assemblies. A plus sign (+) denotes species for which co-obligate symbiosis has been suggested in previous studies.

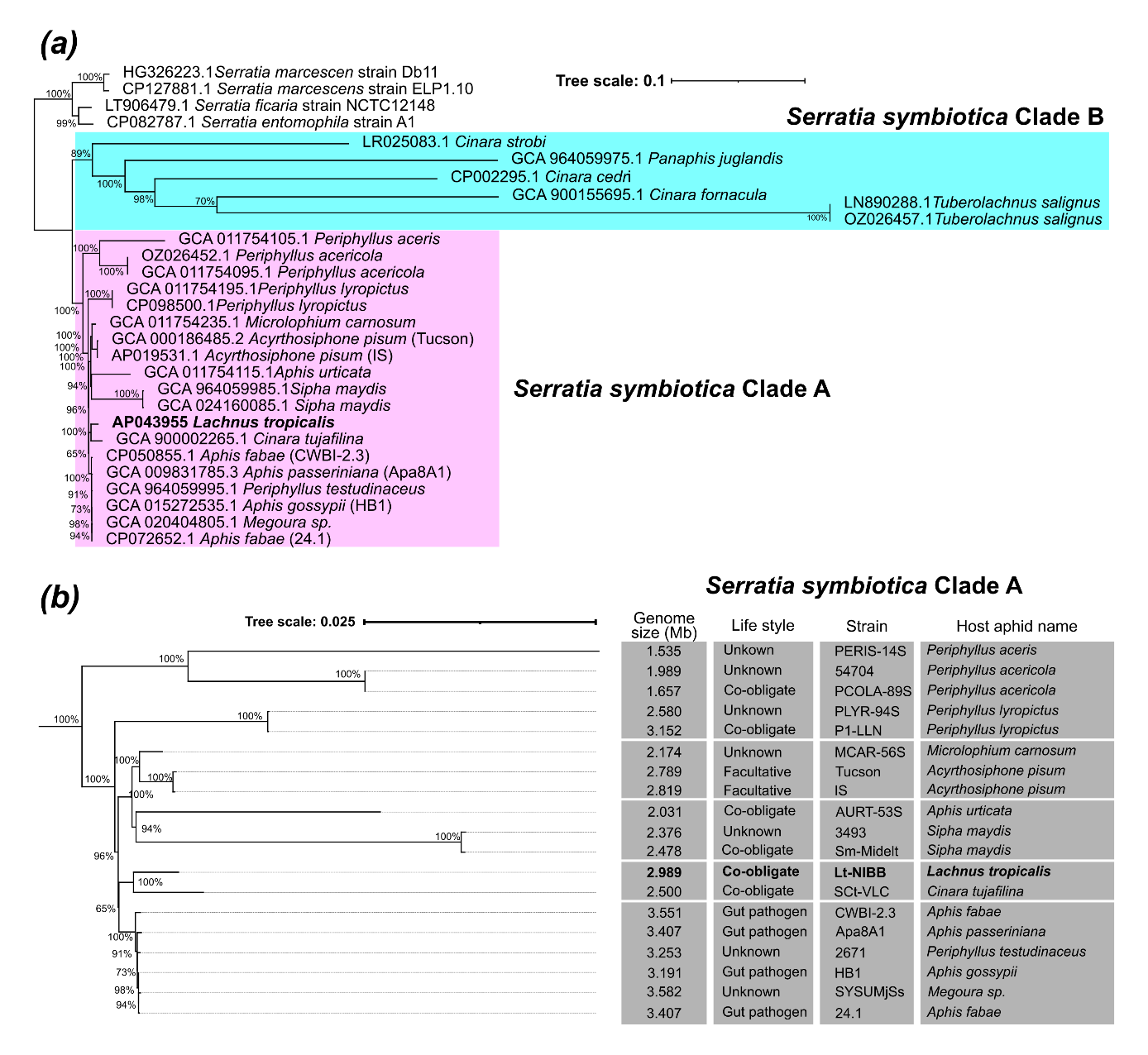
**Figure S6.** **(a)** Phylogenomic tree estimated using IQ-TREE based on a concatenated alignment of 172 core orthologous genes (38,172 amino acids) generated with GToTree. The final maximum-likelihood tree search was completed after 102 iterations. Nodal support was assessed with 1,000 ultrafast bootstrap replicates; bootstrap values ≥70 are shown. **(b)** Enlarged view of *Serratia symbiotica* Clade A. Associated lifestyle and genome size information, from either prior studies or this study, is provided.

**Table S4.** General features of sequenced *Buchnera* genomes of *Lachnus tropicalis* and the related aphid species

| Host aphid species | *Lachnus tropicalis* | *Cinara cedri* | *C. tujaphilina* | *Tuberolachnus salignus* | *Acyrthosiphon pisum* |
| --- | --- | --- | --- | --- | --- |
| Symbiont status | Obligate | Obligate | Obligate | Obligate | Obligate |
| Genome size (bp) | 426,395 | 425,229 | 452,994 | 430,464 | 655,725 |
| Chromosome size (bp) | 419,908 | 416,380 | 444,925 | 421,426 | 640,681 |
| Number of plasmids | 1 | 2 | 1 | 2 | 2 |
| Number of contigs | 2 | 3 | 2 | 3 | 3 |
| GC content (%) | 21.3 | 20.1 | 23.1 | 21.6 | 26.4 |
| Number of CDS (pseudogenes) | 380 (35) | 362 (3) | 407 (44^*1^) | 387 (19) | 574 (0) |
| Number of rRNAs | 3 | 3 | 3 | 3 | 3 |
| Number of tRNAs | 32 | 31 | 30 | 32 | 32 |
| Coding ratio (%) | 85.9 | 85.3 | 80.9 | 87.0 | 87.2 |
| GenBank assembly accession no. | This study | GCA_000090965.1^*2^ | GCA_000217635.1^*3^ | GCA_900016785.1 | GCA_000009605.1 |

^*1^ DFAST initially annotated the plasmid-borne *trpG* gene as a pseudogene (partial). However, by comparing it to *trpG* sequences from other *Buchnera* strains, we concluded it to be intact.

^*2^ The deposited assembly did not include the plasmid pTrp (EU660486). We manually added its FASTA sequence to the assembly before running DFAST annotation.

^*3^ The deposited assembly did not include the fused plasmid pTrpLeu (AY438024). We manually added its FASTA sequence to the assembly before running DFAST annotation.

**Table S5.** General features of sequenced *Serratia symbiotica* genomes of *Lachnus tropicalis* and the related aphid species

| Host aphid species | *L. tropicalis* | *C. cedri* | *C. tujaphilina* | *T. salignus* | *Ac. pisum* | *Aphis fabae* |
| --- | --- | --- | --- | --- | --- | --- |
| Symbiont status | Obligate | Obligate | Obligate | Obligate | Facultative | Gut pathogen |
| *Serratia* clade | Clade A | Clade B | Clade A | Clade B | Clade A | Clade A |
| Strain name | Lt-NIBB | Str. ‘Cinara cedri’ | SCt-VLC | STs | IS | CWBI-2.3 |
| Genome size (bp) | 2,972,445 | 1,762,765 | 2,500,326^*^ | 650,317 | 2,818,957 | 3,550,913 |
| Chromosome size (bp) | 2,870,422 | 1,762,765 | NA | 650,317 | 2,736,352 | 3,351,526 |
| Number of plasmids | 2 | Unknown | Unknown | Unknown | 1 | 2 |
| Number of contigs | 3 | 1 | 32 | 1 | 2 | 3 |
| GC content (%) | 52.1 | 29.2 | 52.1 | 20.9 | 51.9 | 52.0 |
| Number of CDS (pseudogenes) | 3,306 (1160) | 782 (48) | 2,850 (1120) | 500 (16) | 3,155 (1178) | 3,480 (598) |
| Number of rRNAs | 19 | 3 | 12 | 3 | 21 | 22 |
| Number of tRNAs | 57 | 36 | 49 | 33 | 61 | 70 |
| Coding ratio (%) | 80.3 | 39.9 | 78.7 | 77.4 | 79.8 | 81.2 |
| GenBank assembly accession no. | This study | GCA_000238975.1 | GCA_000217635.1 | GCA_900016785.1 | GCA_000009605.1 | GCA_000821185.2 |

^*^ Genome size is reported as the total size of the assembled scaffolds for unclosed genomes

**Table S6.** Sequence (chromosome) list used for *Buchnera* phylogenomic analysis

| ID | Species | Life-style | Host insect | Aphid subfamily | Accession number | Notes (stain name, etc.) |
| --- | --- | --- | --- | --- | --- | --- |
| 1 | *Escherichia coli* | free-living | NA | NA | U00096.3 | Str. K-12 substr. MG1655 |
| 2 | *Ishikawaella capsulata* | obligative | *Megacopta punctatissima* | NA | AP010872.1 | Strain Mpkobe |
| 3 | *Wigglesworthia glossinidia* | obligative | *Glossina morsitans* | NA | CP003315.1 | Isolate WGM |
| 4 | *Buchnera aphidicola* | obligative | *Acyrthosiphon pisum* | Aphidinae | BA000003.2 | Str. APS |
| 5 | *Buchnera aphidicola* | obligative | *Schizaphis graminum* | Aphidinae | AE013218.1 | Str. Sg |
| 6 | *Buchnera aphidicola* | obligative | *Aphis craccivora* | Aphidinae | CP034897.1 | Strain Acr |
| 7 | *Buchnera aphidicola* | obligative | *Aphis urticata* | Aphidinae | CP048744.1 | Isolate AURT-53B |
| 8 | *Buchnera aphidicola* | obligative | *Pentalonia nigronervosa* | Aphidinae | CP061275.1 | Isolate Ba4 |
| 9 | *Buchnera aphidicola* | obligative | *Uroleucon ambrosiae* | Aphidinae | CP002648.1 | Str. Ua |
| 10 | *Buchnera aphidicola* | obligative | *Schlechtendalia chinensis* | Eriosomatinae | CP140041.1 | Isolate R1 |
| 11 | *Buchnera aphidicola* | obligative | *Melaphis rhois* | Eriosomatinae | CP033004.1 | Strain Mrh |
| 12 | *Buchnera aphidicola* | obligative | *Baizongia pistaciae* | Eriosomatinae | AE016826.1 | Str. Bp |
| 13 | *Buchnera aphidicola* | obligative | *Pemphigus populi* | Eriosomatinae | OZ060372.1 | Strain Pepopuli-3700 |
| 14 | *Buchnera aphidicola* | obligative | *Eriosoma grossulariae* | Eriosomatinae | OZ060402.1 | Strain Ergrossulariae-3692 |
| 15 | *Buchnera aphidicola* | obligative | *Tetraneura akinire* | Eriosomatinae | CP188088.1 | Isolate Buch_TA_S2 |
| 16 | *Buchnera aphidicola* | obligative | *Neophyllaphis varicolor* | Neophyllaphidinae | CP135042.1 | Strain Neova26 |
| 17 | *Buchnera aphidicola* | obligative | *Neophyllaphis podocarpi* | Neophyllaphidinae | CP135039.1 | Strain Neopo25 |
| 18 | *Buchnera aphidicola* | obligative | *Mindarus abietinus* | Mindarinae | OZ060398.1 | Strain Miabietinus-3704 |
| 19 | *Buchnera aphidicola* | obligative | *Mindarus keteleerifoliae* | Mindarinae | CP135024.1 | Strain Minke18 |
| 20 | *Buchnera aphidicola* | obligative | *Thelaxes californica* | Thelaxinae | CP034852.1 | Strain Tca |
| 21 | *Buchnera aphidicola* | obligative | *Kurisakia onigurumii* | Thelaxinae | CP135036.1 | Strain Kuron24 |
| 22 | *Buchnera aphidicola* | obligative | *Anoecia corni* | Anoeciinae | OZ060371.1 | Strain Ancorni-2928 |
| 23 | *Buchnera aphidicola* | obligative | *Anoecia oenotherae* | Anoeciinae | CP033012.1 | Strain Aoe |
| 24 | *Buchnera aphidicola* | obligative | *Nipponaphis monzeni* | Hormaphidinae | AP019379.1 | Strain Nmo |
| 25 | *Buchnera aphidicola* | obligative | *Neothoracaphis quercicola* | Hormaphidinae | NA | https://zenodo.org/records/10513209 |
| 26 | *Buchnera aphidicola* | obligative | *Chaitoregma tattakana* | Hormaphidinae | CP134991.1 | Strain Chata04 |
| 27 | *Buchnera aphidicola* | obligative | *Astegopteryx sp.* | Hormaphidinae | NA | https://zenodo.org/records/10513209 |
| 28 | *Buchnera aphidicola* | obligative | *Ceratovacuna japonica* | Hormaphidinae | AP026065.1 | Strain CjNOSY1 |
| 29 | *Buchnera aphidicola* | obligative | *Pseudoregma panicola* | Hormaphidinae | CP134997.1 | Strain Psepa06 |
| 30 | *Buchnera aphidicola* | obligative | *Phyllaphis fagi* | Phyllaphidinae | OZ060370.1 | Strain Phfagi-3703 |
| 31 | *Buchnera aphidicola* | obligative | *Stegophylla sp.* | Phyllaphidinae | CP032998.1 | Strain Ssp |
| 32 | *Buchnera aphidicola* | obligative | *Pterocallis alni* | Calaphidinae | OZ060377.1 | Strain Ptalni-3691 |
| 33 | *Buchnera aphidicola* | obligative | *Myzocallis carpini* | Calaphidinae | OZ060376.1 | Strain Mycarpini-3696 |
| 34 | *Buchnera aphidicola* | obligative | *Panaphis juglandis* | Calaphidinae | OZ060378.1 | Strain Pajuglandis-2786 |
| 35 | *Buchnera aphidicola* | obligative | *Takecallis taiwana* | Calaphidinae | CP134979.1 | Strain Takta17 |
| 36 | *Buchnera aphidicola* | obligative | *Sarucallis kahawaluokalani* | Calaphidinae | CP032999.1 | Strain Ska |
| 37 | *Buchnera aphidicola* | obligative | *Greenidea ficicola* | Greenideinae | CP135012.1 | Strain Grefi11 |
| 38 | *Buchnera aphidicola* | obligative | *Greenidea mangiferae* | Greenideinae | NA | https://zenodo.org/records/10513209 |
| 39 | *Buchnera aphidicola* | obligative | *Drepanosiphum platanoidis* | Drepanosiphinae | OZ026537.1 | Isolate 54803 |
| 40 | *Buchnera aphidicola* | obligative | *Chaitophorus populicola* | Chaitophorinae | OZ060382.1 | Strain Chpopulicola-3609 |
| 41 | *Buchnera aphidicola* | obligative | *Periphyllus lyropictus* | Chaitophorinae | CP097457.1 | Strain Pl-LLN |
| 42 | *Buchnera aphidicola* | obligative | *Periphyllus acericola* | Chaitophorinae | OZ026466.1 | Isolate 54701 |
| 43 | *Buchnera aphidicola* | obligative | *Chaetosiphella stipae* | Chaitophorinae | OZ060394.1 | Strain Chstipae_setosa-2659 |
| 44 | *Buchnera aphidicola* | obligative | *Sipha maydis* | Chaitophorinae | CP097205.1 | Strain Sm Midelt |
| 45 | ***Buchnera aphidicola*** | **obligative** | ***Lachnus tropicalis*** | **Lachninae** | **AP043953** | **Isolate Lt-NIBB (this study)** |
| 46 | *Buchnera aphidicola* | obligative | *Lachnus roboris* | Lachninae | NA | https://zenodo.org/records/10513209 |
| 47 | *Buchnera aphidicola* | obligative | *Tuberolachnus salignus* | Lachninae | LN890285.1 | Strain BTs |
| 48 | *Buchnera aphidicola* | obligative | *Nippolachnus piri* | Lachninae | CP135006.1 | Strain Nippi09 |
| 49 | *Buchnera aphidicola* | obligative | *Cinara confinis* | Lachninae | LT667503.1 | Strain BCiconfinis |
| 50 | *Buchnera aphidicola* | obligative | *Cinara tujafilina* | Lachninae | CP001817.1 | Isolate BCtu |
| 51 | *Buchnera aphidicola* | obligative | *Cinara cedri* | Lachninae | CP000263.1 | Strain BC |
| 52 | *Buchnera aphidicola* | obligative | *Cinara fornacula* | Lachninae | LT667500.1 | Strain BCifornacula |
| 53 | *Buchnera aphidicola* | obligative | *Cinara piceae* | Lachninae | LR217739.1 | Strain BuCipiceae |
| 54 | *Buchnera aphidicola* | obligative | *Cinara splendens* | Lachninae | LR217722.1 | Strain BuCisplendens |

**Table S7.** Sequence (chromosome) list used for *Serratia* phylogenomic analysis

| ID | Species | Life-style | Host insect | Aphid subfamily | Accession number^*^ | Notes (stain name, etc.) |
| --- | --- | --- | --- | --- | --- | --- |
| 1 | *Serratia ficaria* | free-living | NA | NA | LT906479.1 | Strain NCTC12148 |
| 2 | *Serratia marcescens* | free-living | *Drosophila melanogaster* | NA | HG326223.1 | Strain Db11 |
| 3 | *Serratia marcescens* | free-living | NA | NA | CP127881.1 | Strain ELP1.10 |
| 4 | *Serratia entomophila* | pathogenic | *Costelytra zealandica* | NA | CP082787.1 | Strain A1 |
| 5 | *Serratia symbiotica* | gut pathogen | *Aphis passeriniana* | Aphidinae | GCA_009831785.3 | Strain Apa8A1 |
| 6 | *Serratia symbiotica* | gut pathogen | *Aphis fabae* | Aphidinae | CP050855.1 | Strain CWBI-2.3 |
| 7 | *Serratia symbiotica* | gut pathogen | *Aphis gossypii* | Aphidinae | GCA_015272535.1 | Strain HB1 |
| 8 | *Serratia symbiotica* | gut pathogen | *Aphis fabae* | Aphidinae | CP072652.1 | Strain 24.1 |
| 9 | *Serratia symbiotica* | facultative | *Acyrthosiphone pisum* | Aphidinae | GCA_000186485.2 | Strain Tucson |
| 10 | *Serratia symbiotica* | facultative | *Acyrthosiphone pisum* | Aphidinae | AP019531.1 | Strain IS |
| 11 | *Serratia symbiotica* | unknown | *Megoura* sp. | Aphidinae | GCA_020404805.1 | Strain SYSUMjSs |
| 12 | *Serratia symbiotica* | unknown | *Microlophium carnosum* | Aphidinae | GCA_011754235.1 | Strain MCAR-56S |
| 13 | *Serratia symbiotica* | obligative | *Aphis urticata* | Aphidinae | GCA_011754115.1 | Strain AURT-53S |
| 14 | *Serratia symbiotica* | unknown | *Panaphis juglandis* | Calaphidinae | GCA_964059975.1 | Strain Pajuglandis-2786 |
| 15 | *Serratia symbiotica* | unknown | *Periphyllus testudinaceus* | Chaitophorinae | GCA_964059995.1 | Strain Petestudinaceus-2671 |
| 16 | *Serratia symbiotica* | obligative | *Periphyllus lyropictus* | Chaitophorinae | CP098500.1 | Strain Pl-LLN |
| 17 | *Serratia symbiotica* | obligative | *Periphyllus lyropictus* | Chaitophorinae | GCA_011754195.1 | Strain PLYR-94S |
| 18 | *Serratia symbiotica* | obligative | *Periphyllus acericola* | Chaitophorinae | OZ026452.1 | Isolate 54704 |
| 19 | *Serratia symbiotica* | obligative | *Periphyllus acericola* | Chaitophorinae | GCA_011754095.1 | Strain PCOLA-89S |
| 20 | *Serratia symbiotica* | unknown | *Periphyllus aceris* | Chaitophorinae | GCA_011754105.1 | Strain PERIS-14S |
| 21 | *Serratia symbiotica* | obligative | *Sipha maydis* | Chaitophorinae | GCA_024160085.1 | Strain Sm-Midelt-1713498 |
| 22 | *Serratia symbiotica* | obligative | *Sipha maydis* | Chaitophorinae | GCA_964059985.1 | Strain Simaydis-3493 |
| 23 | ***Serratia symbiotica*** | **obligative** | ***Lachnus tropicalis*** | **Lachninae** | **AP043955** | **Isolate Lt-NIBB (this study)** |
| 24 | *Serratia symbiotica* | obligative | *Tuberolachnus salignus* | Lachninae | LN890288.1 | Strain STs |
| 25 | *Serratia symbiotica* | obligative | *Tuberolachnus salignus* | Lachninae | GCA_964019645.1 | Strain TubSali1 |
| 26 | *Serratia symbiotica* | obligative | *Cinara strobi* | Lachninae | LR025083.1 | Strain SeCistrobi |
| 27 | *Serratia symbiotica* | obligative | *Cinara fornacula* | Lachninae | GCA_900155695.1 | Strain Scifornacula |
| 28 | *Serratia symbiotica* | obligative | *Cinara tujaphilina* | Lachninae | GCA_900002265.1 | Strain SCt-VLC |
| 29 | *Serratia symbiotica* | obligative | *Cinara cedri* | Lachninae | CP002295.1 | Str. ‘Cinara cedri’ |

^*^ For assemblies that are not complete chromosomes, we included their assembly accession numbers.

**Chapter 4: Detailed metabolic insights into *L. tropicalis* symbionts**

**Essential amino acid (EAA) synthesis (related to Figures 2B and S7)**

“Symbiotic metabolism” in aphids is best understood through the well-characterized partnership between the pea aphid *Acyrthosiphon pisum* and its obligate symbiont *Buchnera aphidicola* (reviewed in 28). This system clearly demonstrates that metabolic collaboration, particularly in amino acid biosynthesis, operates at the genomic level. Here, we detail the metabolic collaboration among *Buchnera* Lt, *Serratia* Lt, and the aphid *L. tropicalis*, based on the symbiont genomes obtained in this study. Due to the wealth of experimental data on *A. pisum* and the previous unavailability of *L. tropicalis* genomic information, we referenced *A. pisum* (28) for comparative purposes.

Insects generally require 10 EAAs: arginine (Arg), methionine (Met), histidine (His), isoleucine (Ile), valine (Val), leucine (Leu), threonine (Thr), lysine (Lys), phenylalanine (Phe), and tryptophan (Trp) (29). These EAAs are typically scarce in the plant phloem sap consumed by aphids. For instance, in *Vicia faba*, a host plant of the pea aphid, EAAs constitute <1% of the total amino acid content in phloem sap, whereas non-EAAs, such as aspartic acid/asparagine and glutamic acid/glutamine, can make up 90% of the total (30, 31). The amino acid composition and concentration in the phloem exudates of *Quercus dentata*, one of host trees of *L. tropicalis*, have also been investigated (32, 33). Similar to *V. faba*, EAAs were limited, whereas several non-EAAs were relatively abundant: aspartic acid (0.021 ± 0.022 nmol ml^-1^, Mean ± SD), serine (0.031 ± 0.046), asparagine (0.065 ± 0.134), glutamic acid (0.039 ± 0.072), glutamine (0.012 ± 0.019), alanine (0.035 ± 0.051), and proline (0.015 ± 0.022). In contrast, many EAAs were very low or undetected, including threonine (0.007 ± 0.009), glycine (0.005 ± 0.005), valine (0.005 ± 0.009), cysteine (0.001 ± 0.002), isoleucine (0.003 ± 0.004), leucine (0.004 ± 0.005), tyrosine (0.004 ± 0.006), phenylalanine (0.003 ± 0.003), tryptophan (0.001 ± 0.002), lysine (0.003 ± 0.005), histidine (0.001 ± 0.002), and arginine (0.004 ± 0.007). Notably, Methionine was not detected. These data are crucial for inferring the metabolic compensation within the *L. tropicalis* symbiotic system. Based on our symbiont genomic data, we provide a detailed description and discussion of each EAA biosynthetic pathway.

***Arginine biosynthesis pathway***

Our analysis revealed that *Serratia* Lt possesses a nearly complete arginine biosynthesis pathway (Figures 2B, S7; Table S8.1), with only the *carA* gene (encoding the small subunit of glutamine-hydrolyzing carbamoyl phosphate synthase) being pseudogenized due to a frameshift mutation. This enzyme is crucial for the conversion of glutamine into carbamoyl phosphate, which is an early step in arginine production. In contrast, the *Buchnera* Lt genome contains intact *carA* and *carB*, indicating that it can produce carbamoyl phosphate. However, *Buchnera* Lt lacks the *argE* gene, which is required for the later steps of arginine biosynthesis. Collectively, these findings suggest that *Serratia* Lt and *Buchnera* Lt complement each other to complete the arginine biosynthesis pathway. *Buchnera* likely provides the initial carbamoyl phosphate synthesis, whereas *Serratia* completes the subsequent steps. Although the contribution of aphid host genes cannot be completely ruled out, the established model for the pea aphid-*Buchnera* system indicates that arginine biosynthesis is largely symbiont-driven (28).

**Table S8.1.** Genes involving arginine synthesis

| Gene name | Enzyme name | *Buchnera* | *Serratia* |
| --- | --- | --- | --- |
| *carA* | Glutamine-hydrolyzing carbamoyl-phosphate synthase small subunit | present | pseudo |
| *carB* | Glutamine-hydrolyzing carbamoyl-phosphate synthase large subunit | present | present |
| *argF* | Ornithine carbamoyltransferase | present | present |
| *argE* | Acetylornithine deacetylase | absent | present |
| *argG* | Argininosuccinate synthase | present | present |
| *argH* | Argininosuccinate lyase | present | present |

***Methionine biosynthesis pathway***

Methionine biosynthesis in *L*. *tropicalis* is a complex process that relies on a tripartite metabolic collaboration between the symbionts and the host. Although both symbiotic bacteria contribute, the pathway contains critical genetic gaps (Figures 2B, S7; Table S8.2). The pathway begins with the conversion of aspartate into homoserine. This crucial initial step requires enzymes, such as aspartokinase and homoserine dehydrogenase, which are often encoded by genes such as *thrA* or *metL*. Both symbiont genomes possess an intact *thrA*, indicating their capacity to synthesize homoserine. However, the subsequent steps toward homocysteine present several critical bottlenecks. *Serratia* Lt possesses *metA* and *metB*, enabling the synthesis of cystathionine from homoserine and cysteine. However, *metC* is pseudogenized (frameshifted), preventing the direct conversion of cystathionine to homocysteine. In contrast, *Buchnera* lacks *metA*, *metB*, and *metC*, making *de novo* homocysteine synthesis via this route unlikely within *Buchnera*.

Given these genetic gaps, the ability of aphids to supply homocysteine is crucial. The host’s role in this pathway is relevant: while *Serratia* robustly synthesizes cysteine (*cysD* to *cysK* genes are present), which is vital as phloem sap contains only trace amounts of cysteine (32, 33), the subsequent steps for homocysteine production are intricate. Rather than converting cysteine to homocysteine via a forward transsulfuration pathway (which is not typically found in animals), the aphid host is now understood to be involved in producing homocysteine from cystathionine, drawing upon previous findings. Russell et al. (34) demonstrated that homocysteine can be synthesized from cystathionine in the pea aphid system. This suggests that Serratia’s *metC* likely became pseudogenized because the aphid host had already been efficiently performing the cystathionine-β-lyase-equivalent role of converting cystathionine to homocysteine. This preexisting host capability would have rendered *Serratia*’s *metC* redundant, removing the evolutionary pressure to maintain its functionality.

Ultimately, the generated homocysteine, likely through the aphid host’s metabolic machinery utilizing cystathionine (potentially supplied by *Serratia* after *metB* activity), was supplied to *Buchnera*. *Buchnera* possesses the *metE* gene and performs the final step of converting homocysteine to methionine, which is then provided back to the aphid host. Notably, *Serratia*’s *metE* gene is also pseudogenized (frameshift), further highlighting *Buchnera*’s indispensable role in this terminal methionine synthesis step. In summary, the direct pathway from aspartate to homocysteine appears incomplete in the symbionts, owing to specific gene losses or pseudogenization. Instead, this system orchestrates a more flexible and robust metabolic network for the supply of sulfur-containing amino acids, leveraging the aphid host’s ability to complement symbiotic genetic deficiencies. This crucial metabolic detour through the aphid host machinery ensures a continuous supply of this EAA.

**Table S8.2.** Genes involving methionine synthesis

| Gene name | Enzyme name | *Buchnera* | *Serratia* |
| --- | --- | --- | --- |
| *thrA* | Bifunctional aspartate kinase/homoserine dehydrogenase I | present | present |
| *metL* | Bifunctional aspartokinase/homoserine dehydrogenase II | absent | present |
| *metA* | Homoserine O-succinyltransferase | absent | present |
| *metB* | Cystathionine gamma-synthase | absent | present |
| *metC* | Cystathionine beta-lyase | absent | pseudo |
| *metE* | 5-methyltetrahydropteroyltriglutamate--homocysteine methyltransferase | present | pseudo |
| *cysD* | Sulfate adenylyltransferase small subunit | absent | present |
| *cysN* | Sulfate adenylyltransferase large subunit | absent | present |
| *cysC* | Adenylyl-sulfate kinase | absent | present |
| *cysH* | Phosphoadenylyl-sulfate reductase | absent | present |
| *cysI* | NADPH-dependent sulfite reductase heme-binding subunit | absent | present |
| *cysJ* | NADPH-dependent sulfite reductase flavoprotein subunit | absent | present |
| *cysE* | Serine O-acetyltransferase | absent | present |
| *cysK* | Methionine adenosyltransferase | absent | present |

***Histidine biosynthesis pathway***

*Serratia* Lt possesses a complete set of genes (*hisG*–*hisD*; Table S8.3) required for the full biosynthesis of histidine from PRPP and ATP. This suggests that *Serratia* can synthesize histidine *de novo*, serving as the primary source of this EAA for aphids. In contrast, *Buchnera* Lt lacks almost all the genes in the histidine biosynthetic pathway, rendering it incapable of independent histidine synthesis. This absence is likely a consequence of *Buchnera*’s extreme genome reduction. Notably, this pathway is maintained in *Buchnera* strains associated with other Lachninae species, such as *Cinara cedri* and *Tuberolachnus salignus* (11, 26), indicating a more advanced state of genome degeneration in *Buchnera* in *L. tropicalis*.

This pattern demonstrates adaptations associated with a newly established multiple-partner symbiosis. Although *Buchnera* has lost its histidine synthesis capability, *Serratia* fully compensates, ensuring a reliable supply of histidine. This represents an excellent example of individual symbiotic bacteria specializing in specific functions, where *Buchnera* has streamlined its genome by relinquishing certain metabolic pathways; *Serratia* steps in to fill that crucial role.

**Table S8.3.** Genes involving histidine synthesis

| Gene name | Enzyme name | *Buchnera* | *Serratia* |
| --- | --- | --- | --- |
| *hisG* | ATP phosphoribosyltransferase | absent | present |
| *hisIE* | Bifunctional phosphoribosyl-AMP cyclohydrolase/phosphoribosyl-ATP diphosphatase | absent | present |
| *hisA* | 1-(5-phosphoribosyl)-5-[(5-phosphoribosylamino) methylideneamino] imidazole-4-carboxamide isomerase | absent | present |
| *hisH* | Imidazole glycerol phosphate synthase subunit | absent | present |
| *hisF* | Imidazole glycerol phosphate synthase subunit | absent | present |
| *hisB* | Bifunctional histidinol-phosphatase/imidazoleglycerol-phosphate dehydratase | absent | present |
| *hisC* | Histidinol-phosphate transaminase | absent | present |
| *hisD* | Histidinol dehydrogenase | absent | present |

***Valine and leucine biosynthesis pathways***

Our analysis revealed that *Buchnera* Lt possesses the functional *ilvH* (*ilvB*, *ilvG*), *ilvI* (*ilvN*, *ilvM*), and *ilvC* (Table S8.4) genes. This indicates that *Buchnera* produces the early intermediate 2-oxoisopentanoate. Although *Serratia* Lt also carries these initial genes, its dihydroxy-acid dehydratase (*ilvD* gene) is pseudogenized (frameshifted), suggesting that *Serratia* cannot independently complete the conversion to 2-oxoisopentanoate and thus relies on *Buchnera* for this critical intermediate.

This pathway subsequently diverges toward valine and leucine synthesis. The final amination step, catalyzed by branched-chain amino acid aminotransferase (*ilvE*), is absent in *Buchnera* Lt. However, the pea aphid possesses an enzyme capable of converting the keto acid intermediates into valine and leucine (28), suggesting that the host itself plays a direct role in this crucial final step, also in this case. For leucine-specific steps, *Serratia* Lt carries several pseudogenized *leu* genes (*leuA*: stop codon and frameshift; *leuC*: frameshift; and *leuB*: frameshift), indicating that this pathway is relinquishing. This pattern aligns with the presence of plasmid-encoded *leu* genes in *Buchnera* Lt, which are typically highly expressed to support efficient amino acid production. Although theoretically *Serratia* Lt could perform the final amination using *ilvE* if supplied with keto acid precursors from *Buchnera* Ltd, the established role of the aphid host in related systems and *Serratia*’s gene losses suggest that the host remains the primary contributor to this step.

In summary, branched-chain amino acid synthesis in *L. tropicalis* involves a coordinated division of labor: *Buchnera* Lt performs early and leucine-specific steps, *Serratia* Lt contributes partially to early steps, and the aphid host executes the critical final amination. This intricate metabolic collaboration ensures a robust and reliable supply of these EAAs, leveraging the complementary strengths of its symbiotic partners and their metabolic capabilities.

**Table S8.4.** Genes involving valine and leucine synthesis

| Gene name | Enzyme name | *Buchnera* | *Serratia* |
| --- | --- | --- | --- |
| *ilvH* (*ilvB*, *ilv*G) | Acetolactate synthase large subunit | present | present |
| *ilvI* (*ilvN*, *ilvM*) | Acetolactate synthase small subunit | present | present |
| *ilvC* | Ketol-acid reductoisomerase | present | present |
| *ilvD* | Dihydroxy-acid dehydratase | present | pseudo |
| *ilvE* | Branched-chain amino acid transaminase | absent | present |
| *leuA* | 2-isopropylmalate synthase | present* | pseudo |
| *leuC* | 3-isopropylmalate dehydratase large subunit | present* | pseudo |
| *leuD* | 3-isopropylmalate dehydratase small subunit | present* | present |
| *leuB* | 3-isopropylmalate dehydrogenase | present* | pseudo |

Note: the asterisk (*) indicates that the gene is located on a plasmid

***Isoleucine biosynthesis pathway***

Our genomic analysis revealed that neither *Buchnera* nor *Serratia* possesses a complete pathway for *de novo* isoleucine synthesis (Table S8.5). Although *Serratia* retains the initial *ilvA* gene starting from threonine and the final *ilvE* gene for amination, its *ilvD* gene is pseudogenized (frameshifted), preventing the production of the necessary intermediates. Conversely, *Buchnera* has functional *ilvH/I*, *ilvC*, and *ilvD* genes, allowing it to process intermediates but lacking both the initial *ilvA* and the final *ilvE*. This creates a critical gap in metabolism. This intricate dependency suggests a complex metabolic exchange between symbionts; however, it strongly implicates the aphid host as a key player in bridging this gap. Although *Serratia* initiates the pathway and *Buchnera* processes intermediates, the aphid likely contributes the missing enzymatic steps, potentially providing the initial threonine or a necessary intermediate, or even performing parts of the synthesis through its own metabolic machinery to ensure a continuous supply of this EAA.

**Table S8.5.** Genes involving isoleucine synthesis

| Gene name | Enzyme name | *Buchnera* | *Serratia* |
| --- | --- | --- | --- |
| *ilvA* | L-threonine dehydratase biosynthetic | absent | present |
| *ilvH* (*ilvB*, *ilv*G) | Acetolactate synthase large subunit | present | present |
| *ilvI* (*ilvN*, *ilvM*) | Acetolactate synthase small subunit | present | present |
| *ilvC* | ketol-acid reductoisomerase | present | present |
| *ilvD* | Dihydroxy-acid dehydratase | present | pseudo |
| *ilvE* | Branched-chain amino acid transaminase | absent | present |

***Threonine and lysine biosynthesis pathways***

The biosynthesis of threonine and lysine, both EAAs derived from aspartate, revealed a sophisticated metabolic division of labor between *Buchnera* Lt and *Serratia* Lt within *L*. *tropicalis* (Table S8.6). For threonine, the pathway from aspartate to the final product appeared fully functional in both symbionts, as evidenced by the presence of all necessary genes (*thrA*, *asd*, *thrB*, and *thrC)*. Notably, *Serratia* possesses two copies of the *asd* gene, potentially enhancing its capacity for this early step in both threonine and lysine pathways. This redundancy suggests a robust and resilient supply of threonine to aphids from both partners.

Lysine biosynthesis, however, presents a more intricate picture that relies largely on *Serratia*. Although the initial steps from aspartate as well as *dapA*, *dapB*, and *dapD,* are present in both symbionts, a key enzyme—*dapC* (succinyldiaminopimelate transaminase)—is absent in both *Buchnera* Lt and *Serratia* Lt. Despite this, both harbor *serC* (3-phosphoserine/phosphohydroxythreonine transaminase). As highlighted by Lal et al. (35), the N-acetylornithine aminotransferase activity required for lysine synthesis can be performed by the product of *serC*. Thus, *serC* can functionally substitute for *dapC*, effectively bridging this enzymatic gap.

Further along the pathway, both symbionts encode *dapE* and *lysA*. However, a crucial difference lies in *dapF*: in *Buchnera*, *dapF* is a pseudogene (frameshifted), preventing it from completing lysine biosynthesis. Conversely, *Serratia* Lt retains functional *dapF,* and with its active *serC* (compensating for *dapC*), it emerges as the primary contributor to *de novo* lysine production. This intricate genetic distribution underscores how complementary gene sets and functional substitutions between the two symbionts orchestrate the provision of EAAss to their aphid host.

**Table S8.6.** Genes involving threonine and lysine synthesis

| Gene name | Enzyme name | *Buchnera* | *Serratia* |
| --- | --- | --- | --- |
| *thrA* | Bifunctional aspartate kinase/homoserine dehydrogenase I | present | present |
| *asd* | Aspartate-semialdehyde dehydrogenase | present | present (2) |
| *thrB* | Homoserine kinase | present | present |
| *thrC* | Threonine synthase | present | present |
| *dapA* | 4-hydroxy-tetrahydrodipicolinate synthase | present | present |
| *dapB* | 4-hydroxy-tetrahydrodipicolinate reductase | present | present |
| *dapD* | 2,3,4,5-tetrahydropyridine-2,6-dicarboxylate N-succinyltransferase | present | present |
| *dapC* | Succinyldiaminopimelate transaminase | absent | absent |
| *serC* | 3-phosphoserine/phosphohydroxythreonine transaminase | present | present |
| *dapE* | Succinyl-diaminopimelate desuccinylase | present | present |
| *dapF* | Diaminopimelate epimerase | pseudo | present |
| *lysA* | Diaminopimelate decarboxylase | present | present |

***Phenylalanine and tryptophan biosynthesis pathways***

The biosynthesis of phenylalanine and tryptophan, both EAAs, relied almost entirely on *Serratia* Lt within the *L*. *tropicalis* aphid system (Table S8.7). This pathway begins with two ubiquitous precursors: phosphoenolpyruvate (PEP), a key intermediate in glycolysis, and erythrose 4-phosphate, derived from the pentose phosphate pathway. Both are readily available from the central carbon metabolism within the host or symbiont. These precursors then feed into the shikimate pathway, leading to the formation of a common intermediate, chorismate.

The shikimate pathway highlights *Serratia*’s indispensable role from the very first step. Although most genes for chorismate synthesis (*aroB*, *aroQ*, *aroE*, *aroK*, *aroA*, and *aroC*) were present in both *Buchnera* Lt and *Serratia* Lt, *aroG*, which codes the crucial initial enzyme (3-deoxy-7-phosphoheptulonate synthase) was absent in *Buchnera* but present in *Serratia*. This means *Buchnera* Lt cannot initiate the shikimate pathway *de novo*, making *Serratia* Lt solely responsible for converting PEP and erythrose 4-phosphate into chorismate. This chorismate then served as a branching point for the synthesis of the final aromatic amino acids. For phenylalanine, *Serratia* Lt possesses the necessary genes, such as *pheA* (bifunctional chorismate mutase/prephenate dehydratase) and *hisC* (histidinol-phosphate transaminase), whereas *Buchnera* Lt lacks them. Similarly, for tryptophan, *Serratia* Lt harbors all the required genes, including *trpE*, *trpG*, *trpD*, *trpCF* (bifunctional), *trpB*, and *trpA*, all of which are absent in *Buchnera* Lt. This complete set of genes in *Serratia* Lt for both specific branches firmly establishes its role as the primary, if not exclusive, producer of both phenylalanine and tryptophan.

In summary, *Buchnea*’s extensive gene loss in aromatic amino acid biosynthesis makes it entirely reliant on *Serratia* Lt. This symbiotic arrangement reflects a clear metabolic specialization, where *Serratia* Lt acts as a dedicated factory for essential aromatic amino acids, leveraging readily available central carbon metabolites to ensure that the nutritional needs of the aphid host are met.

**Table S8.7.** Genes involving phenylalanine and tryptophane synthesis

| Gene name | Enzyme name | *Buchnera* | *Serratia* |
| --- | --- | --- | --- |
| *aroG* | 3-deoxy-7-phosphoheptulonate synthase | absent | present |
| *aroB* | 3-dehydroquinate synthase | present | present |
| *aroQ* | Type II 3-dehydroquinate dehydratase | present | present |
| *aroE* | Shikimate dehydrogenase | present | present |
| *aroK* | Shikimate kinase | present | present |
| *aroA* | 3-phosphoshikimate 1-carboxyvinyltransferase | present | present |
| *aroC* | Chorismate synthase | present | present |
| *pheA* | Bifunctional chorismate mutase/prephenate dehydratase | absent | present |
| *hisC* | Histidinol-phosphate transaminase | absent | present |
| *trpE* | Anthranilate synthase component 1 | absent | present |
| *trpG* | Gamma-glutamyl-gamma-aminobutyrate hydrolase family protein^*^ | absent | present |
| *trpD* | Anthranilate phosphoribosyltransferase | absent | present |
| *trpCF* | Bifunctional indole-3-glycerol-phosphate synthase & phosphoribosylanthranilate isomerase | absent | present |
| *trpB* | Tryptophan synthase subunit beta | absent | present |
| *trpA* | Tryptophan synthase subunit alpha | absent | present |

^*^Corresponding to anthranilate synthase component II in *Serratia symbiotica*　(WP_061770748.1)

**Figure S7.** Metabolic complementation for the biosynthesis of 10 essential amino acids (EAAs) in the *Lachnus tropicalis* endosymbiotic system. The *Serratia* Lt genome exhibits a relatively broad capacity for EAA synthesis, although pseudogenization was detected in the methionine pathway (*metC* and *metE*). These functions are presumably compensated by *Buchnera* (for *metE*) or the aphid host (for *metC*). In addition, *Serratia* Lt contains pseudogenes in the leucine pathway (*leuA*, *leuC*, and *leuB*). Notably, all of these leucine genes are located on the *Buchnera* plasmid, pLeu. Although *dapC* is absent in both symbiont genomes, an alternative pathway using *serC* is present.
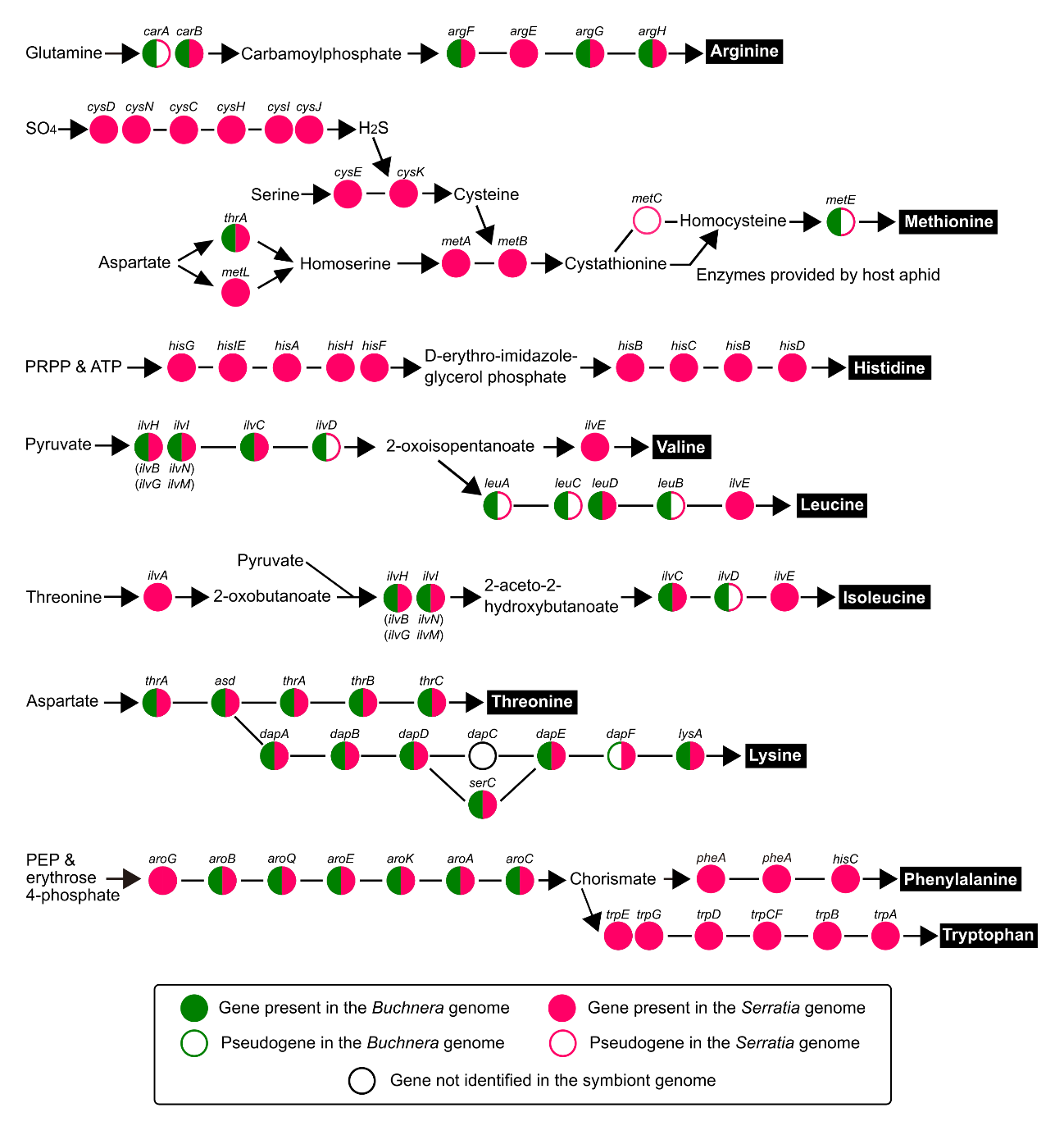

**Cofactor synthesis (related to Figure 2B, S8)**

In this study, we focused on eight cofactors that play essential roles in animal metabolism: riboflavin (vitamin B2), biotin (vitamin B7), folate (vitamin B9), pyridoxal 5’-phosphate (vitamin B6), pantothenate (vitamin B5), coenzyme A (CoA), nicotinamide adenine dinucleotide (NAD+), and thiamine (vitamin B1).

In *Buchnera-*monosymbiotic aphids, such as the pea aphid, B vitamins are thought to be synthesized through a combination of *Buchnera* and host genes (36). However, some *Buchnera* lineages in Lachninae, Hormaphidinae, and Chaitophorinae exhibit a somewhat reduced capacity for B-vitamin biosynthesis. Previous studies on multiplex symbiosis in aphids have primarily focused on how riboflavin (B2) and biotin (B7) are provisioned by symbionts (4, 12, 37, 38, 39, 40). Considering the gene repertoires in the *Serratia* genomes of *Cinara cedri*, *C. tujaphilina*, and *T. salignus*, these pathways appear to be complemented by these symbiont partners rather than *Buchnera* (11, 4). Although a relevant discussion remains regarding how these synthesis pathways are complemented and how they interact with the host (36), here, we organize how the synthesis pathways for these cofactors are potentially covered based on the genomic information of *Buchnera* and *Serratia* in *L. tropicalis*. Below, we present the results of our gene presence or absence analysis for specific pathways and discuss plausible metabolic complementation.

***Riboflavin: vitamin B2 biosynthesis pathway***

Our genomic analysis showed that *Buchnera* in *L. tropicalis* lacked all genes necessary for the *de novo* synthesis of riboflavin (Table S9.1). Therefore, *Buchnera* from *L. tropicalis* is unable to produce vitamin B2 on its own, similar to what has been observed in Lachninae aphids (*Cinara* and *Tuberolachnus*; Lamelas et al., 2011; 11). In contrast, *Serratia* in *L. tropicalis* possesses all genes required for the complete riboflavin biosynthesis pathway, indicating that *Serratia* retains the full genetic machinery to synthesize Vitamin B2. Therefore, it is strongly suggested that in *L. tropicalis*, the symbiont *Serratia* is the primary producer of riboflavin (vitamin B2) in the aphid host.

**Table S9.1.** Genes involving riboflavin synthesis

| Gene name | Enzyme name | *Buchnera* | *Serratia* |
| --- | --- | --- | --- |
| *ribA* | GTP cyclohydrolase II | absent | present |
| *ribD* | Bifunctional diaminohydroxyphosphoribosylaminopyrimidine deaminase/5-amino-6-(5-phosphoribosylamino) uracil reductase | absent | present |
| *yigB* | 5-amino-6-(5-phospho-D-ribitylamino) uracil phosphatase | absent | present |
| *ribH* | 6,7-dimethyl-8-ribityllumazine synthase | absent | present |
| *ribB* | 3,4-dihydroxy-2-butanone-4-phosphate synthase | absent | present |
| *ribE* | Riboflavin synthase | absent | present |

***Biotin: vitamin B7***

Based on our genomic analysis of *L. tropicalis*, the *Buchnera* symbiont lacks most of the genes essential for *de novo* biotin (vitamin B7) synthesis, including key enzymes such as *bioC*, *fabB*, *fabF*, *bioH*, *bioF*, *bioA*, *bioD*, and *bioB* (Table S9.2). Although *Buchnera* retains *fabG* and *fabZ*, the pathway remains incomplete. Notably, *Buchnera* also possesses *fabI*, which is absent in *Serratia* Lt. This suggests a potential complementation in which *Buchnera* could contribute *fabI* to the biotin synthesis pathway.

In contrast, *Serratia* Lt harbors nearly all the genes required for complete biotin biosynthesis, including *bioC*, *fabB*, *fabF*, *bioH*, *bioF*, *bioA*, *bioD*, and *bioB*. Although it lacks *fabI*, its overall comprehensive gene repertoire indicates a near-complete capacity to synthesize vitamin B7, potentially relying on *Buchnera* for the *fabI* step or an alternative pathway. Therefore, in *L. tropicalis*, *Serratia* Lt is strongly suggested to be the primary producer of biotin (vitamin B7), with *Buchnera* potentially complementing the pathway through the *fabI* gene. This highlights a complicated inter-symbiont metabolic cooperation rather than reliance on a single partner.

**Table S9.2.** Genes involving biotin synthesis

| Gene name | Enzyme name | *Buchnera* | *Serratia* |
| --- | --- | --- | --- |
| *bioC* | Malonyl-ACP O-methyltransferase | absent | present |
| *fabB* | Beta-ketoacyl-ACP synthase I | absent | present |
| *fabF* | Beta-ketoacyl-ACP synthase II | absent | present |
| *fabG* | 3-oxoacyl-ACP reductase | present | present |
| *fabZ* | 3-hydroxyacyl-ACP dehydratase | present | present |
| *fabI* | Enoyl-ACP reductase | present | absent |
| *bioH* | Pimeloyl-ACP methyl ester esterase | absent | present |
| *bioF* | 8-amino-7-oxononanoate synthase | absent | present |
| *bioA* | Adenosylmethionine--8-amino-7-oxononanoate transaminase | absent | present |
| *bioD* | Dethiobiotin synthase | absent | present |
| *bioB* | Biotin synthase | absent | present |

***Folate: vitamin B9***

Based on our data, the *Buchnera* symbiont lacks all the genes necessary for *de novo* folate (vitamin B9) synthesis (Table S9.3). This indicates that *Buchnera* from *L. tropicalis* is unable to produce vitamin B9 independently. In contrast, *Serratia* possesses nearly all of the genes required for the complete folate biosynthesis pathway, including *folE*, *folX*, *folB*, *folK*, *pabB*, *pabC*, *folP*, *folC*, and *folA*. Notably, *Serratia* lacks a *pabA*. However, it is known that when exogenous ammonia is available, *pabB* can potentially function in the absence of *pabA*. Indeed, certain *Serratia symbiotica* strains, such as *Serratia symbiotica* 24.1, also lack *pabA* but retain a class I glutamine amidotransferase, suggesting alternative mechanisms for ammonia provision or the presence of a divergent *pabA* homolog. Despite this single missing gene, the overall gene repertoire of *Serratia* strongly supports its capacity to synthesize vitamin B9. Therefore, in *L. tropicalis, Serratia* is most likely the primary—and possibly the sole—producer of folate (vitamin B9) for the aphid host.

**Table S9.3.** Genes involving folate synthesis

| Gene name | Enzyme name | *Buchnera* | *Serratia* |
| --- | --- | --- | --- |
| *folE* | GTP cyclohydrolase I | absent | present |
| *folX* | Dihydroneopterin triphosphate 2’-epimerase | absent | present |
| *folB* | Bifunctional dihydroneopterin aldolase/7,8-dihydroneopterin epimerase | absent | present |
| *folK* | 2-amino-4-hydroxy-6-hydroxymethyldihydropteridine diphosphokinase | absent | present |
| *pabA* | Glutamine amidotransferase subunit | absent | absent |
| *pabB* | Aminodeoxychorismate synthase component 1 | absent | present |
| *pabC* | Aminodeoxychorismate lyase | absent | present |
| *folP* | Dihydropteroate synthase | absent | present |
| *folC* | Bifunctional tetrahydrofolate synthase/dihydrofolate synthase | absent | present |
| *folA* | Type 3 dihydrofolate reductase | absent | present |

***Pyridoxal 5’-phosphate: vitamin B6***

Our analysis of *L. tropicalis* revealed that *Buchnera* Lt entirely lacked almost all genes necessary for *de novo* pyridoxal 5’-phosphate (vitamin B6) synthesis (Table S9.4). Specifically, *epd*, *pdxB*, *pdxA*, *pdxJ*, and *pdxH* were absent. Although *Buchnera* retained *serC*, this single gene was insufficient to complete the pathway, indicating that *Buchnera* from *L. tropicalis* cannot produce citamin B6 on its own. In contrast, *Serratia* Lt possessed most of the genes required for the vitamin B6 biosynthesis pathway, including *pdxB*, *serC*, *pdxA*, *pdxJ*, and *pdxH*.

However, both *Buchnera* and *Serratia* lacked *epd*. This suggests that neither symbiont maintained a fully independent pathway for vitamin B6 biosynthesis. The shared absence of *epd* in both symbionts strongly implies that the aphid host may supply the missing precursor normally generated by *epd*, or that an alternative, as-yet-uncharacterized pathway or gene compensates for its absence within the host or symbiont. Therefore, the provisioning of pyridoxal 5’-phosphate (vitamin B6) in *L. tropicalis* appears to rely on a complex metabolic collaboration. Although *Serratia* contributes the majority of the pathway genes, the absence of *epd* in both symbionts suggests a crucial role for host-derived precursors or alternative complementary mechanisms within this tripartite symbiotic system.

**Table S9.4.** Genes involving pyridoxal 5’-phosphate synthesis

| Gene name | Enzyme name | *Buchnera* | *Serratia* |
| --- | --- | --- | --- |
| *epd* | D-erythrose-4-phosphate dehydrogenase | absent | absent |
| *pdxB* | 4-phosphoerythronate dehydrogenase | absent | present |
| *serC* | 3-phosphoserine/phosphohydroxythreonine transaminase | present | present |
| *pdxA* | 4-hydroxythreonine-4-phosphate dehydrogenase | absent | present |
| *pdxJ* | Pyridoxine 5’-phosphate synthase | absent | present |
| *pdxH* | Pyridoxamine 5’-phosphate oxidase | absent | present |

***Pantothenate: vitamin B5 and CoA***

Our genomic analysis of *L. tropicalis* revealed a clear division of labor for pantothenate (vitamin B5) and CoA synthesis (Table S9.5). The *Buchnera* genome lacked nearly all the genes necessary for *de novo* synthesis of both pantothenate and CoA. Although it retained *ilvH*, *ilvI*, *ilvC,* and *ilvD* (which participate in an upstream pathway), it was missing essential pantothenate-specific genes such as *panB*, *panE*, *panD*, *panC*, as well as all downstream CoA synthesis genes (*coaA*, *coaBC*, *coaD*, *coaE*). Notably, *ilvE* was absent, suggesting that *Buchnera* from *L. tropicalis* is unable to produce either pantothenate or CoA on its own.

In contrast, *Serratia* in *L. tropicalis* possessed a comprehensive set of genes required for pantothenate and CaoA biosynthesis. These included *ilvH*, *ilvI*, *ilvC*, *ilvE*, *panE*, *panD*, *panC*, *coaA*, *coaBC*, *coaD*, and *coaE*. Importantly, *Serratia* carried two copies of *panB*. Although *ilvD* was a pseudogene (frameshift) in *Serratia* (but present in *Buchnera*), the extensive gene repertoire of *Serratia*, particularly the duplication of *panB*, strongly indicates that it retains almost complete genetic machinery for *de novo* synthesis of both vitamin B5 and CoA. Therefore, in *L. tropicalis*, *Serratia* is likely the primary (if not sole) producer of both pantothenate (vitamin B5) and CoA for the aphid host. This highlights *Serratia*’s vital role in supplying these essential cofactors, which are largely absent in *Buchnera*.

**Table S9.5.** Genes involving pantothenate and coenzyme A synthesis

| Gene name | Enzyme name | *Buchnera* | *Serratia* |
| --- | --- | --- | --- |
| *ilvH* (*ilvB*, *ilv*G) | Acetolactate synthase large subunit | present | present |
| *ilvI* (*ilvN*, *ilvM*) | Acetolactate synthase small subunit | present | present |
| *ilvC* | Ketol-acid reductoisomerase | present | present |
| *ilvD* | Dihydroxy-acid dehydratase | present | pseudo |
| *ilvE* | Branched-chain amino acid transaminase | absent | present |
| *panB* | 3-methyl-2-oxobutanoate hydroxymethyltransferase | absent | present (2) |
| *panE* | 2-dehydropantoate 2-reductase | absent | present |
| *panD* | Aspartate 1-decarboxylase | absent | present |
| *panC* | Pantoate--beta-alanine ligase | absent | present |
| *coaA* | Type I pantothenate kinase | absent | present |
| *coaBC* | Bifunctional phosphopantothenoylcysteine decarboxylase/phosphopantothenate--cysteine ligase | absent | present |
| *coaD* | Pantetheine-phosphate adenylyltransferase | absent | present |
| *coaE* | Dephospho-CoA kinase | absent | present |

***NAD+: nicotinamide adenine dinucleotide synthesis pathway***

Our genomic analysis of *L. tropicalis* revealed that neither *Buchnera* nor *Serratia* possesses a complete *de novo* NAD+ biosynthesis pathway (Table S9.6). The *Buchnera* genome lacks all the genes necessary for NAD+ synthesis, including *nadB*, *nadA*, *nadC*, *nadD*, *pncB*, and *nadE*, indicating that *Buchnera* from *L. tropicalis* is incapable of producing NAD+ independently.

Similarly, *Serratia* lacks a full *de novo* pathway. Although it retains *nadA*, *nadD*, and *pncB*, crucial genes such as *nadC* and *nadE* are absent. Furthermore, *nadB* has undergone pseudogenization (frameshift and internal stop codon), rendering it nonfunctional. The absence of *nadC* and *nadE*, together with the nonfunctional *nadB*, confirms that *Serratia* is unable to synthesize NAD+ *de novo*. These results suggest that the aphid host must obtain NAD+ or its precursors from external sources, most likely through direct dietary intake or an alternative, yet uncharacterized pathway in the host. This underscores a critical metabolic dependency of *L. tropicalis* on its diet or host physiology for maintaining sufficient levels of this essential cofactor.

**Table S9.6.** Genes involving NAD+ synthesis

| Gene name | Enzyme name | *Buchnera* | *Serratia* |
| --- | --- | --- | --- |
| *nadB* | L-aspartate oxidase | absent | pseudo |
| *nadA* | Quinolinate synthase | absent | present |
| *nadC* | Quinolinate phosphoribosyltransferase | absent | absent |
| *nadD* | Nicotinate-nucleotide adenylyltransferase | absent | present |
| *pncB* | Nicotinate phosphoribosyltransferase | absent | present |
| *nadE* | NAD+ synthetase | absent | absent |

***Thiamine: vitamin B1 biosynthesis pathway***

Genomic analysis of *L. tropicalis* showed that *Buchnera* completely lacks almost all genes necessary for *de novo* thiamine (vitamin B1) synthesis (Table S9.7). These include crucial genes such as *thiI*, *thiH*, *thiG*, *tenI*, *thiC*, *thiD*, *thiE*, and *rsgA*. Although *Buchnera* possesses *iscS* (cysteine desulfurase), it is classified within the IscS subfamily of cysteine desulfurase, which may indicate a broader role or a specialized variant. Nevertheless, *Buchnera*’s overall gene deficiency indicates that incapable of synthesizing vitamin B1 independently.

In contrast, *Serratia* possesses the majority of the genes necessary for a nearly complete thiamine biosynthesis pathway. These include *iscS*, *thiI*, *thiH*, *thiG*, *thiC*, *thiD*, *thiE*, and *rsgA*. However, both *Buchnera* and *Serratia* lack *tenI* (thiazole tautomerase). Although this enzyme can accelerate the tautomerization step in thiamine biosynthesis, the reaction can occur spontaneously in its absence, and thus its loss may not completely disrupt the pathway. Despite this shared absence, *Serratia*’s robust gene repertoire strongly indicates that it has the full genetic machinery to synthesize vitamin B1. Therefore, it is strongly suggested that, in *L. tropicalis*, the symbiont *Serratia* is the primary producer of thiamine (vitamin B1) in aphid hosts. The shared absence of *tenI* in both symbionts highlights a potential dependency on a spontaneous reaction or an alternative mechanism within the symbiotic system; however, *Serratia* clearly possesses the vast majority of the required machinery.

**Table S9.7.** Genes involving thiamine synthesis

| Gene name | Enzyme name | *Buchnera* | *Serratia* |
| --- | --- | --- | --- |
| *iscS* | Cysteine desulfurase | present | present |
| *thiI* | tRNA 4-thiouridine(8) synthase | absent | present |
| *thiH* | 2-iminoacetate synthase | absent | present |
| *thiG* | Thiazole synthase | absent | present |
| *tenI* | Thiazole tautomerase | absent | absent |
| *thiC* | Phosphomethylpyrimidine synthase | absent | present |
| *thiD* | Bifunctional hydroxymethylpyrimidine kinase/phosphomethylpyrimidine kinase | absent | present |
| *thiE* | Thiamine phosphate synthase | absent | present |
| *rsgA* | Small ribosomal subunit biogenesis GTPase | absent | present |

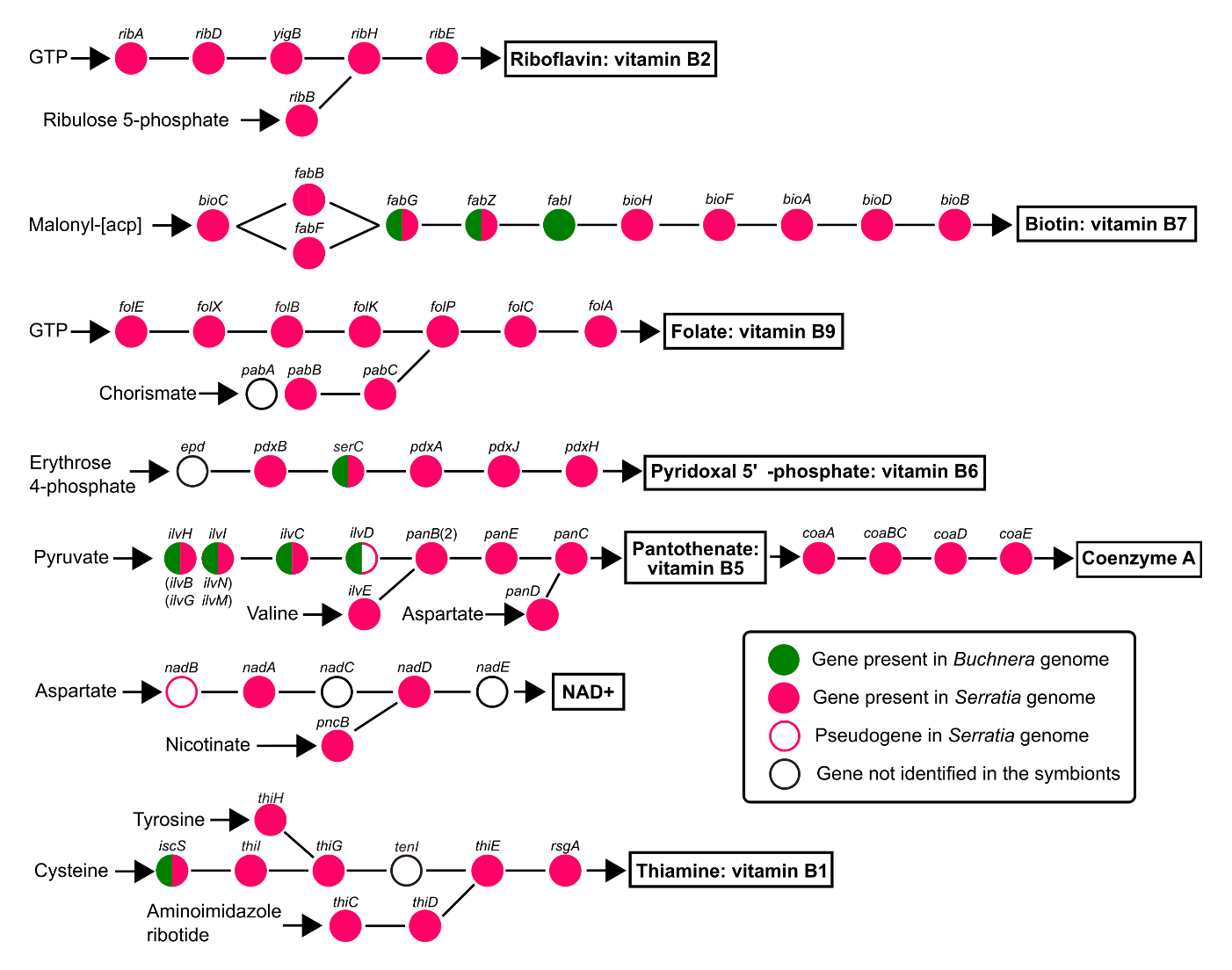
**Figure S8.** Metabolic complementation for cofactor biosynthesis in the *Lachnus tropicalis* endosymbiotic system. *Serratia* Lt exhibits a broad capacity for cofactor synthesis. Notably, *Serratia* Lt can completely synthesize riboflavin (vitamin B2), a pathway entirely being absent in *Buchnera* Lt. In contrast, biotin (vitamin B7) biosynthesis is likely cooperatively compensated by both symbionts.

**Other genes related to free-living lifestyles: cell wall synthesis, cell division, and cell motility**

In addition to EAA and cofactor biosynthesis, other fundamental bacterial characteristics crucial for free-living lifestyles, such as cell wall synthesis, cell division, and motility, often undergo substantial changes or losses in symbiotic organisms (28, 41, 42). These genes are frequently lost or pseudogenized in symbiotic owing to reduced selective pressure within the stable host environment. Therefore, we investigated these genes to characterize symbiotic adaptations and genome streamlining in *L. tropicalis* symbionts, using *E. coli* as a reference (K-12, MG1655; GCF_000005845.2).

***Cell wall proteins***

It is well established that insect endosymbionts often lose their ability to synthesize cell walls (41), which is considered an adaptation to a symbiotic lifestyle. Many insect endosymbionts, including *Buchnera*, as well as some pathogenic bacteria with parasitic lifestyles, lack a complete pathway for cell wall synthesis and often do not possess a cell wall (42). These functional losses affect cell morphology, division mechanisms, host interactions (such as immune recognition), and growth rates. From the host’s perspective, avoiding the costly process of cell wall synthesis in a stable intracellular environment can be considered energy-saving and efficient. In the current study, we observed that *Buchnera* and *Serratia* in *L. tropicalis* exhibited distinct morphologies: the former was coccoid (as typical of *Buchnera*), whereas the latter was rod-shaped (Figure 3). As shown in Table S10, we analyzed genes involved in peptidoglycan metabolism.

*Buchnera* Lt, similar to other *Buchnera* species with small genomes (approximately 0.4 Mb; such as Lachninae: *Cinara cedri*, 26; Hormaphidinae: *Ceratovacuna japonica*, 12), lacked almost all genes in this category, with the exception of the lipid II flippase (*murJ*). This strongly suggests that *Buchnera* Lt has completely lost the ability to synthesize peptidoglycans *de novo*. Notably, *Buchnera* species of Aphidinae, such as those from pea-aphids, retain relatively more of these genes (although in many cases *murF* is absent, suggesting that they cannot synthesize a cell wall independently, or only maintain a very limited one) (42). The complete loss of cell wall synthesis capability in *Buchnera* Lt can be viewed as one of the further functional reductions underlying the evolution of dual symbiosis.

In contrast, *Serratia* Lt, similar to *E. coli*, retains the majority of core genes involved in peptidoglycan precursor biosynthesis (such as *glmS* to *murF*, *mraY*, *murG*), as well as major polymerization and cross-linking enzymes (such as class A/B PBPs). This suggests that *Serratia* Lt can independently synthesize a functional cell wall. However, some gene loss and pseudogenization were detected, especially in genes related to degradation and recycling of peptidoglycan precursors and peptidoglycan itself (*ldcA* [frameshift and internal stop codon], *ampH* [frameshift], *rlpA* [frameshift and an internal stop codon], and *amiC* [frameshift]). In addition to the pseudogenized *amiC*, *Serratia* Lt harbors two intact N-acetylmuramoyl-L-alanine amidases, although their specific identities (which *ami* genes they correspond to) could not be determined. There is another N-acetylmuramoyl-l-alanine amidase that is pseudogenized owing to a frameshift. In total, *Serratia* Lt carries five amidase homologs, three of which are pseudogenes. These patterns suggest that efficient recycling and specific cell separation mechanisms may be impaired compared to free-living bacteria. This likely represents an adaptation to the symbiotic environment, where reduced recycling demands or controlled suspension of excessive cell wall synthesis may be advantageous.

**Table S10**. Peptidoglycan pathway genes in *Buchnera* Lt, *Serratia* Lt, and *Escherichia coli*

| Category | Gene Name | *Buchnera* Lt | *Serratia* Lt | *Escherichia coli* |
| --- | --- | --- | --- | --- |
| **Precursor Biosynthesis** |  |  |  |  |
| Initial Sugar Phosphate Precursors | *glmS* | absent | present | present |
|  | *glmM* | absent | present | present |
|  | *glmU* | absent | present | present |
| Mur Peptide Synthesis | *murA* | absent | present | present |
|  | *murB* | absent | present | present |
|  | *murC* | absent | present | present |
|  | *murD* | absent | present | present |
|  | *murE* | absent | present | present |
|  | *murF* | absent | present | present |
| D-Ala-D-Ala Synthesis/Racemase | *ddlA* | absent | absent | present |
|  | *ddlB* | absent | absent | present |
|  | *dadX* | absent | absent | present |
|  | *alr* | absent | present | present |
|  | *murI* | absent | present | present |
| Transfer to Lipid Carrier | *mraY* | absent | present | present |
|  | *murG* | absent | present | present |
| **Membrane Transport** |  |  |  |  |
| Lipid II Flippase | *murJ* | present | present | present |
| **Peptidoglycan Polymerization & Cross-linking** |  |  |  |  |
| Class A PBP (Polymerization/Cross-linking) | *mrcA* | absent | present | present |
|  | *mrcB* | absent | present | present |
|  | *pbpC* | absent | absent | present |
| Class B PBP (Cross-linking) | *mrdA* | absent | present | present |
|  | *ftsI* | absent | present | present |
| Membrane-associated Glycosyltransferase (Polymerization) | *mtgA* | absent | absent | present |
|  | *ftsW* | absent | present | present |
|  | *mrdB* | absent | present | present |
| L, D-Transpeptidase (Cross-linking) | *ldtA* | absent | absent | present |
|  | *ldtB* | absent | present | present |
|  | *ldtC* | absent | absent | present |
|  | *ldtD* | absent | present | present |
|  | *ldtE* | absent | absent | present |
| **Cell Wall Remodeling & Degradation** |  |  |  |  |
| Low MW PBP (Remodeling) | *dacA* | absent | present | present |
|  | *dacC* | absent | absent | present |
|  | *dacD* | absent | absent | present |
|  | *dacD* | absent | absent | present |
|  | *dacB* | absent | present | present |
|  | *ampH* | absent | pseudo | present |
|  | *yfeW* | absent | absent | present |
|  | *pbpG* | absent | absent | present |
| Endopeptidase (Degeneration) | *mepA* | absent | present | present |
|  | *mepH* | absent | absent | present |
|  | *mepS* | absent | absent | present |
|  | *mepM* | absent | present | present |
| Lytic Transglycosylase (Degradation/Remodeling) | *slt (sltY)* | absent | present | present |
|  | *mltA* | absent | present | present |
|  | *mltB* | absent | present | present |
|  | *mltC* | absent | present | present |
|  | *mltD* | absent | present | present |
|  | *emtA* | absent | present | present |
|  | *mltF* | absent | present | present |
|  | *mltG* | absent | present | present |
|  | *rlpA* | absent | pseudo | present |
| Amidase (Degeneration) | *amiA* | absent | absent | present |
|  | *amiC* | absent | pseudo | present |
|  | *amiB* | absent | present | present |
|  | *amiD* | absent | absent | present |
| **Peptidoglycan Recycling** |  |  |  |  |
| Degradation Product Processing | *ddpX* | absent | absent | present |
|  | *ldcA* | absent | pseudo | present |
|  | *ampD* | absent | absent | present |
| Lipid Carrier Recycling | *bacA* | absent | absent | present |
|  | *ybjG* | absent | present | present |
| **Cell Wall Structure** |  |  |  |  |
| Structural Protein | *lpp* | absent | absent | present |

***Cell division, DNA segregation, and cell shape***

Beyond cell-wall biosynthesis, the integrity of bacterial life fundamentally relies on the precise control of cell division, accurate DNA segregation, maintenance of cell morphology, and robust global regulatory networks. The distinct morphologies of *Buchnera* Lt (coccoid) and *Serratia* Lt (rod-shaped) in *L. tropicalis* (Figure 3) prompted us to investigate their gene complements related to these essential cellular processes. As shown in Table S11, comparative genomic analyses revealed striking differences between these key processes.

*Buchnera* Lt exhibits a profound loss of genes within these functional categories, mirroring the extreme genome reduction characteristics of many obligate intracellular endosymbionts (42). With the exception of a few core components like *ftsZ*, *ftsA*, and *dnaA*, nearly all other genes for cell division, DNA segregation, and global regulation are absent (as reported in different strains; 12, 26). These extensive gene losses imply heavy reliance on the host for essential cellular processes, likely resulting in inefficient or host-dependent cell division and DNA segregation. Such functional streamlining in *Buchnera* Lt is particularly intriguing in light of reports of asymmetric division in other genome-reduced cockroach symbionts, *Blattabacterium* (genome size approximately 0.6 Mb, comparable to pea-aphid *Buchnera*) (43). Further detailed observations of *Buchnera* Lt cell division are therefore of great interest.

*Serratia* Lt, in contrast to *Buchnera* Lt, largely retains a comprehensive gene repertoire essential for cell division, DNA segregation, cell shape maintenance, and regulatory functions, similar to those in free-living *E. coli*. This suggests that *Serratia* Lt maintains a high degree of autonomy in fundamental cellular processes. However, certain gene losses and pseudogenization indicate adaptation to its symbiotic lifestyle. Specifically, while many Z-ring components are present, *zapC* and *zapE* are pseudogenes owing to frameshift mutations. Furthermore, *Serratia* Lt possesses two intact copies of *xerD*, which are involved in DNA topology and segregation, along with a third *xerD* copy that is a pseudogene due to a frameshift and an internal stop codon. Impairment of these auxiliary Z-ring factors could lead to subtle defects in cell division efficiency or accuracy, potentially resulting in altered cell morphology (such as cell chaining or uneven division) compared to *E. coli*. Notably, *ftsK*, critical for chromosome segregation, was annotated as “partial” by DFAST, although its counterpart in *Serratia symbiotica* (MBF1995254.1) suggests a potentially functional status. In addition, *Serratia* Lt lacks the DNA-binding proteins *cbpM* and *cbpA*, as well as the cell division-related peptidase *acrE*, which might affect specific aspects of gene expression regulation and daughter cell separation, respectively. The presence of two intact copies of *fic* is also noteworthy, potentially indicating a specialized regulatory role. These genomic features suggest that *Serratia* Lt, while largely self-sufficient, has undergone some streamlining, possibly favoring resource conservation within the stable host environment by reducing the need for certain redundant or energetically costly precise regulatory and cell separation mechanisms found in free-living bacteria. Future studies examining the actual division process and subtle morphological features of *Serratia* Lt could reveal this symbiont, as a relatively recent acquisition, to be a valuable illustration of the early processes of symbiotic evolution.

**Table S11**. Genes related to cell division, DNA segregation, and cell shape in *Buchnera* Lt, *Serratia* Lt, and *Escherichia coli*

| Category | Gene Name | *Buchnera* Lt | *Serratia* Lt | *Escherichia coli* |
| --- | --- | --- | --- | --- |
| **Cell Division (Z-ring Formation & Septum Formation)** |  |  |  |  |
| Z-ring Formation & Regulation | *ftsZ* | present | present | present |
|  | *zipA* | absent | present | present |
|  | *zapA* | present | present | present |
|  | *zapB* | absent | present | present |
|  | *zapC* | absent | pseudo | present |
|  | *zapD* | absent | present | present |
|  | *zapE* | absent | pseudo | present |
| Divisome Assembly | *ftsA* | present | present | present |
|  | *ftsE* | absent | present | present |
|  | *ftsX* | absent | present | present |
|  | *ftsI* | absent | present | present |
|  | *ftsQ* | absent | present | present |
|  | *ftsL* | absent | present | present |
|  | *ftsB* | absent | present | present |
|  | *ftsN* | absent | present | present |
| Cell Division Site Selection | *minC* | present | present | present |
|  | *minD* | present | present | present |
|  | *minE* | present | present | present |
| DNA Replication & Cell Division Link | *sulA* | absent | present | present |
|  | *slmA* | absent | present | present |
| **DNA Segregation & Structure** |  |  |  |  |
| Chromosome Segregation (Condensation/Decatenation) | *mukB* | absent | present | present |
|  | *mukE* | absent | present | present |
|  | *mukF* | absent | present | present |
|  | *ftsK* | absent | present | present |
|  | *parC* | absent | present | present |
|  | *parE* | absent | present | present |
| DNA Topology & Segregation | *xerC* | absent | present | present |
|  | *xerD* | absent | present (2) | present |
| DNA Replication Initiation | *dnaA* | present | present | present |
| DNA Binding & Structural Maintenance | *seqA* | absent | present | present |
|  | *ihfA* | absent | present | present |
|  | *ihfB* | present | present | present |
| **Cytoskeleton & Cell Shape Determination** |  |  |  |  |
| MreB System (Rod Shape) | *mreB* | absent | present | present |
|  | *mreC* | absent | present | present |
|  | *mreD* | absent | present | present |
|  | *mrdB* | absent | present | present |
| **Post-Translational Modification & Protein Quality Control** |  |  |  |  |
| rRNA Methylation | *mnmG* | present | present | present |
|  | *rsmG* | absent | present | present |
| Chaperone & Protease | *hspQ* | absent | present | present |
|  | *hejK* | absent | present | present |
| **Global Regulators & Transcriptional Regulation** |  |  |  |  |
| DNA-binding Proteins | *lrp* | absent | present | present |
|  | *fis* | absent | present | present |
|  | *diaA* | absent | present | present |
|  | *argP* | absent | present | present |
|  | *cbpM* | absent | absent | present |
|  | *cbpA* | absent | absent | present |
| Other Regulators | *acrA* | absent | present | present |
|  | *acrE* | absent | absent | present |
|  | *mrp* | absent | present | present |
|  | *fic* | absent | present (2) ^*^ | present |
|  | *yihA* | present | present | present |
|  | *hfq* | absent | present | present |

^*^ Identified as a Fic family protein gene.

***Cell motility; flagellar assembly***

Motility is crucial for environmental exploration and resource acquisition by free-living bacteria. As an obligate intracellular endosymbiont with an extremely reduced genome, *Buchnera* Lt is expected to be non-motile because of the extensive loss of most genes involved in flagellar assembly and the complete absence of all chemotaxis genes (Table S12). Although a few core components, such as *fliP* and *fliQ,* are retained and *fliF* is present as a nearly full-length gene, *fliN* is pseudogenized (severely truncated). The absence of the vast majority of other essential flagellar components indicates that *Buchnera* Lt cannot form a functional flagellum. The retention of these flagellar genes likely represents vestigial remnants of a motile ancestor or may reflect the acquisition of new, non-motility-related functions within its symbiotic lifestyle.

*Serratia* Lt retained a substantial number of flagellar assembly genes (Table S12). However, a detailed examination revealed critical losses and pseudogenization that strongly suggest that *Serratia* Lt is non-motile or has severely impaired motility. Key components such as *fliI* (internal stop codon), *flhA* (frameshift), *fliF* (internal stop codon), *flgK* (internal stop codon), and *flhC* (frameshift) are pseudogenized, while *fliT* is absent. Most notably, the essential stator component *motA* is absent, which is critical for generating flagellar torque and effectively renders a functional flagellar motor impossible. Furthermore, *Serratia* Lt has lost all chemotaxis genes, indicating a complete inability to move toward or away from stimuli. Therefore, despite retaining many flagellar genes, the absence of crucial components (particularly *motA* and the entire chemotaxis system) indicates that *Serratia* Lt is non-motile. Its current genetic state likely reflects an ongoing process of gene degradation typical of the early stages of genome streamlining in newly acquired symbionts, where selective pressure for motility is considerably reduced within a stable host environment (42).

**Table S12**. Genes related to the flagellar system in *Buchnera* Lt, *Serratia* Lt, and *Escherichia coli*

| Category | Gene Name | *Buchnera* Lt | *Serratia* Lt | *Escherichia coli* |
| --- | --- | --- | --- | --- |
| **Flagellar Assembly Proteins** |  |  |  |  |
| Type-III Secretion | *fliH* | absent | present | present |
|  | *fliI* | absent | pseudo | present |
|  | *fliJ* | absent | present | present |
|  | *fliO* | absent | present | present |
|  | *fliP* | present | present | present |
|  | *fliQ* | present | present | present |
|  | *fliR* | absent | present | present |
|  | *flhA* | absent | pseudo | present |
|  | *flhB* | absent | present^*1^ | present |
|  | *flhE* | absent | present | present |
| C Ring | *fliG* | absent | present | present |
|  | *fliM* | absent | present | present |
|  | *fliN* | pseudo^*2^ | present | present |
| M, S, P, and L Rings | *fliF* | present^*3^ | pseudo | present |
|  | *flgI* | absent | present | present |
|  | *flgA* | absent | present | present |
|  | *flgH* | absent | present | present |
|  | *fliL* | absent | present | present |
| Rod and Hook | *fliE* | absent | present | present |
|  | *fliK* | absent | present | present |
|  | *flgB* | absent | present | present |
|  | *flgC* | absent | present | present |
|  | *flgD* | absent | present | present |
|  | *flgF* | absent | present | present |
|  | *flgG* | absent | present | present |
|  | *flgJ* | absent | present | present |
|  | *flgE* | absent | present | present |
|  | *flgK* | absent | pseudo | present |
|  | *flgL* | absent | present | present |
|  | *flgN* | absent | present | present |
| Filament | *fliC* | absent | present^*4^ | present |
|  | *fliD* | absent | present | present |
|  | *fliS* | absent | present | present |
|  | *fliT* | absent | absent | present |
| Stator | *flhD* | absent | present | present |
|  | *flhC* | absent | pseudo | present |
|  | *fliA* | absent | present | present |
|  | *motA* | absent | absent | present |
|  | *motB* | absent | present | present |
| Others | *fliZ* | absent | present | present |
|  | *flgM* | absent | present | present |
| **Chemotaxis Proteins** |  |  |  |  |
| Two-Component System Proteins | *cheA* | absent | absent | present |
|  | *cheW* | absent | absent | present |
|  | *cheR* | absent | absent | present |
|  | *cheB* | absent | absent | present |
|  | *cheY* | absent | absent | present |
|  | *cheZ* | absent | absent | present |
| MCPs | *tsr* | absent | absent | present |
|  | *tar* | absent | absent | present |
|  | *trg* | absent | absent | present |
|  | *tap* | absent | absent | present |
|  | *aer* | absent | absent | present |

*1 *Serratia* Lt’s *flhB* gene was annotated as a pseudogene (partial sequence) by DFAST. However, a BLAST search revealed that it was a full-length gene of *Serratia symbiotica*, leading us to classify it as intact.

*2 *Buchnera* Lt’s *fliN* is annotated as a pseudogene (partial sequence). Indeed, a BLAST search revealed that the sequence contained 55% of the length of an intact FliM/FliN family flagellar motor switch protein from a related *Buchnera* species (a significant N-terminal truncation).

*3 *Buchnera* Lt *fliF* gene was annotated as a pseudogene (partial sequence) using DFAST. However, a BLAST search confirmed that the *fliF* gene was nearly full-length, leading to its intact classification.

*4 Identified as a FliC/FljB family flagellin.

**Chapter 5: Detailed protocols and observations for *L. tropicalis* symbionts**

**General structure of the aphid bacteriome (symbiotic organ)**

We conducted a literature survey on the bacteriome structure of aphids harboring *Serratia symbiotica*. Based on these descriptions, we summarized the general structure of the aphid bacteriome and provided schematic illustrations of bacteriome organization and the localization of both *Buchnera* and *Serratia* in well-described cases. Examples include *Aphis fabae* (Clade A, gut symbiont; 44), *Acyrthosiphone pisum* (Clade A, facultative and intracellular symbiont; 45, 46), *Periphyllus lyropictus* and *Sipha maydis* (Clade A, obligate and intra/extracellular symbiont; 39), *Cinara cedri* (Clade B, obligate and intracellular symbiont; 47, 48), and *Lachnus roboris* (Clade B, possibly obligate and intracellular symbiont; 49). Details are provided in the caption of Figure S9.

**Detailed methods for imaging**

To obtain detailed information on the cellular features of *L. tropicalis* bacteriome, we performed morphological observations of dissected bacteriomes. Late-instar viviparous nymphs and young viviparous adults collected from the NIBB campus were dissected in phosphate-buffered saline (PBS; 33 mM KH_2_PO_4_ and 33 mM Na_2_HPO_4_, pH 7.2) under a stereomicroscope (SZ61; Olympus, Japan) using fine forceps. Dissections were performed immediately after collection. Dissected tissues were fixed in 4% paraformaldehyde (PFA) in PBS for approximately 3 h and then washed three times with PBS-Tx (0.3% Triton X-100 in PBS). Finally, these tissues were stained with 4,6-diamidino-2-phenylindole (DAPI) (1 μg/mL; Dojindo, Japan) for nuclei and Alexa Fluor 488 phalloidin (66 nM; Thermo Fisher Scientific, USA) for F-actin, respectively. Samples were mounted on glass slides with VECTASHIELD (Vector Laboratories, USA), covered with cover slips, and observed with a confocal laser scanning microscope (FV1000; Olympus, Japan).

To visualize tissue localization and vertical transmission of symbionts in *L. tropicalis*, we conducted fluorescent in situ hybridization (FISH) on dissected bacteriomes and viviparous embryos. Symbiont-specific probes targeted the 16S rRNA gene sequences: *Buchnera aphidicola* (5’-Cy5- CCTCATCTAGGTAGATCCCC-3’) and *Serratia symbiotica* (PASSisR: 5’-Cy3- CCCGACTTTATCGCTGGC-3’ [50]). The *Buchnera* probes were custom-designed based on sequence data, targeting the same region frequently hybridized by the *A. pisum* *Buchnera* probe Apis2a (50). Viviparous individuals were dissected immediately in PBS. Bacteriomes, embryos, and gut tissues from late-instar nymphs and young adults were fixed in 4% PFA in PBS for approximately 3 h, then transferred to Carnoy’s solution (ethanol: chloroform: glacial acetic acid = 6:3:1 [v/v]) for more rigid fixation. Fixed samples were treated overnight with alcoholic 6% H_2_O_2_ to reduce insect tissue autofluorescence (52). Tissues were washed three times with PBS-Tx, followed by hybridization buffer (20 mM Tris-HCl, pH 8.0, 0.9 M NaCl, 0.01% sodium dodecyl sulfate, 30% [v/v] formamide) before hybridization. Samples were incubated overnight at 25–28 ℃ in hybridization buffer containing both probes (final concentration 100 nM each) and DAPI (1 μg/mL; Dojindo, Japan). After incubation, samples were washed three times with PBS-Tx, mounted, and observed as described above. Control samples processed identically but without FISH probes confirmed that autofluorescence was not significant.

**Viviparous embryonic development and symbiont transmission in *L. tropicalis***

We categorized *L. tropicalis* embryos into eight developmental stages: oocyte, syncytial blastoderm, cellular blastoderm stages I and II, invagination, segmentation, flipping, and final growth (Figures S10 and S11). This categorization is based primarily on descriptions of pea aphid, embryogenesis (53) and *Ceratovacuna japonica* (12), but informed by recent observations of embryogenesis and symbiont transmission in the Lachnine aphid *Cinara cedri* (48).

Figures S10 (up to the flip stage) and S11 (final growth stages) present our findings on the localization of symbiotic bacteria at each embryonic stage, vertical transmission patterns, and allocation to developing bacteriome cells (bacteriocytes and sheath cells), as determined by FISH. These data show that *Buchnera* is incorporated into bacteriocytes via cellularization, while *Serratia* infects and migrates into sheath cells following their formation. To more precisely capture symbiont migration, especially that of *Serratia* into bacteriocytes during the final growth stage, we subdivided this stage into three substages (final growth stages I, II, and III).

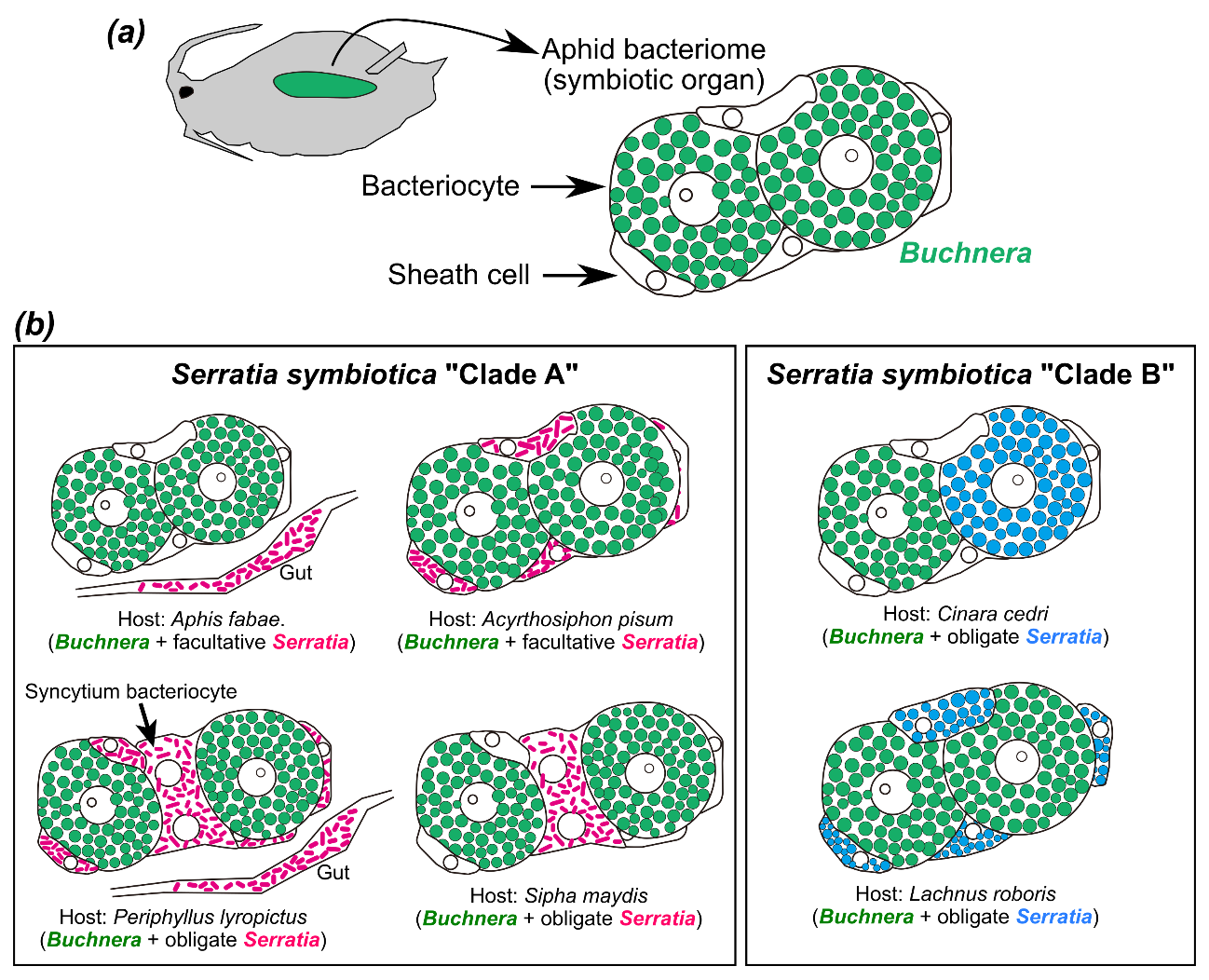
**Figure S9.** General structure of the aphid bacteriome and diverse *Serratia symbiotica* localization patterns. **(a)** Schematic illustration of an aphid body and its bacteriome (symbiotic organ). The aphid bacteriome generally consists of bacteriocytes and sheath cells, though cell types and their distribution can vary (49, 54). *Buchnera* is always housed within the large, polyploid bacteriocytes. In contrast, sheath cells are flattened, relatively small, and never contain *Buchnera*. **(b)** Localization patterns of *Serratia symbiotica* in well-documented cases. Clade A *S. symbiotica* encompasses culturable, facultative, and obligate strains, exhibiting diverse tissue localization. For instance, *Serratia* is a gut symbiont in *Aphis fabae* (44). In *Acyrthosiphon pisum*, *Serratia* is facultative and primarily localizes in sheath cells (45). In *Peryphyllus lyropictus* and *Sipha maydis*, *Serratia* is considered a newly integrated essential symbiont, localizing to a novel type of bacteriocyte (syncytium), and also infecting gut and sheath cells in *P. lyropictus* (39). Clade B *S. symbiotica* consists of obligate symbionts with reduced genomes. In *Cinara cedri*, *Buchnera* and *Serratia* are housed in distinct bacteriocytes (48). In *Lachnus roboris*, *Serratia* is harbored in sheath cells (49).

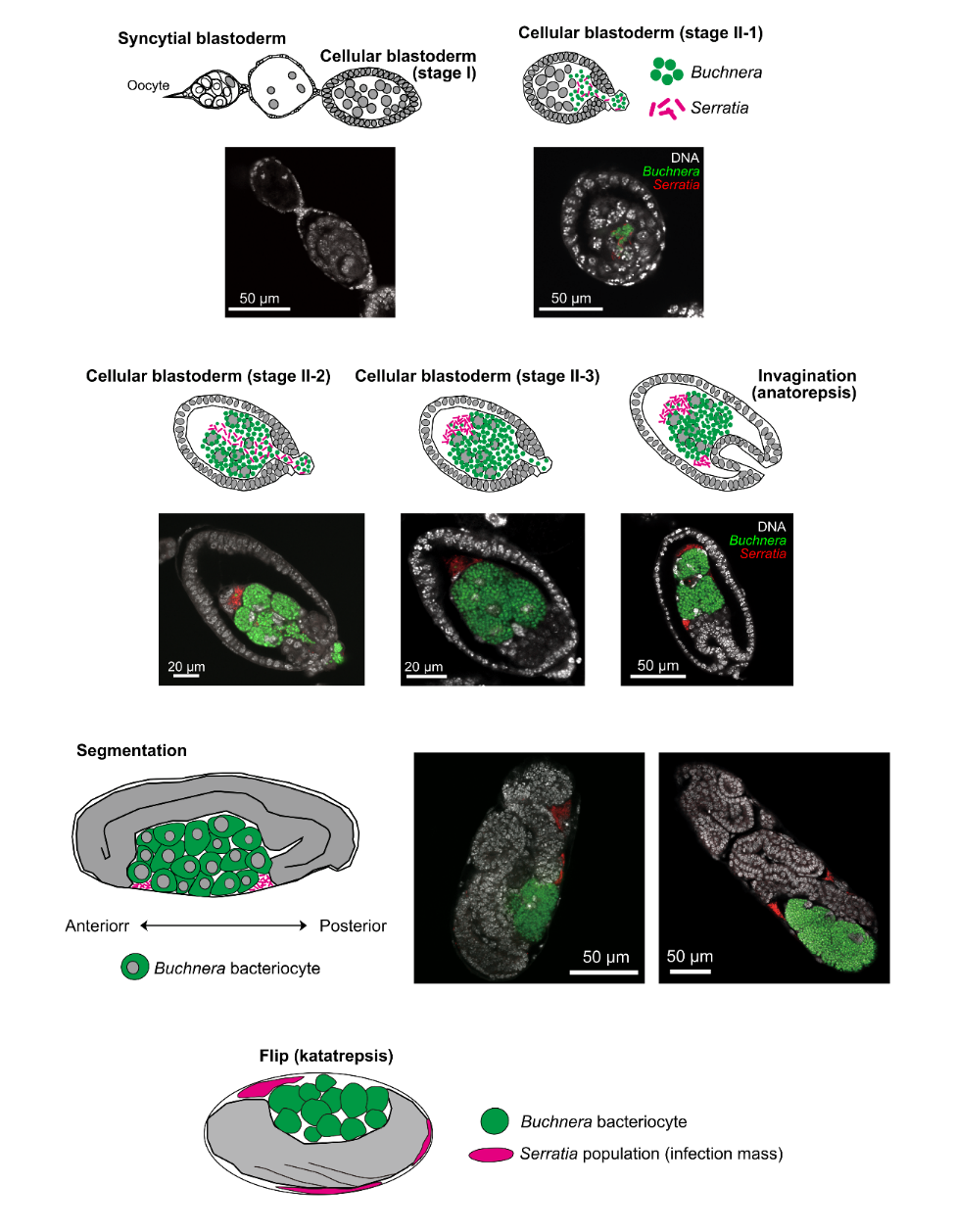
**Figure S10.** Vertical transmission of *Buchnera* and *Serratia*, and early stages of viviparous embryonic development in *Lachnus tropicalis*. Illustrations and representative microscopic images of early embryonic stages, detailing the localization of symbiotic bacteria. Following Nozaki et al. (48), embryos were categorized into several stages, including oocyte, syncytial blastoderm, cellular blastoderm, invagination (anatrepsis), segmentation, and flip (katatrepsis) stages, leading up to the final growth stage (Figure S11). Briefly, the vertical transmission of both *Buchnera* and *Serratia* was observed during the cellular blastoderm stage II. Initially, *Buchnera* colonized and formed clusters surrounding the nuclei (presumably future bacteriocyte nuclei) (stage II-2). Subsequently, *Serratia* cells penetrate these *Buchnera* clusters and form an anteriorly positioned “infection mass” (stage II-3). During invagination, segmentation, and flip stages, *Serratia* infection masses remained in a non-cellular state within peripheral zones or gaps between forming organs and tissues.

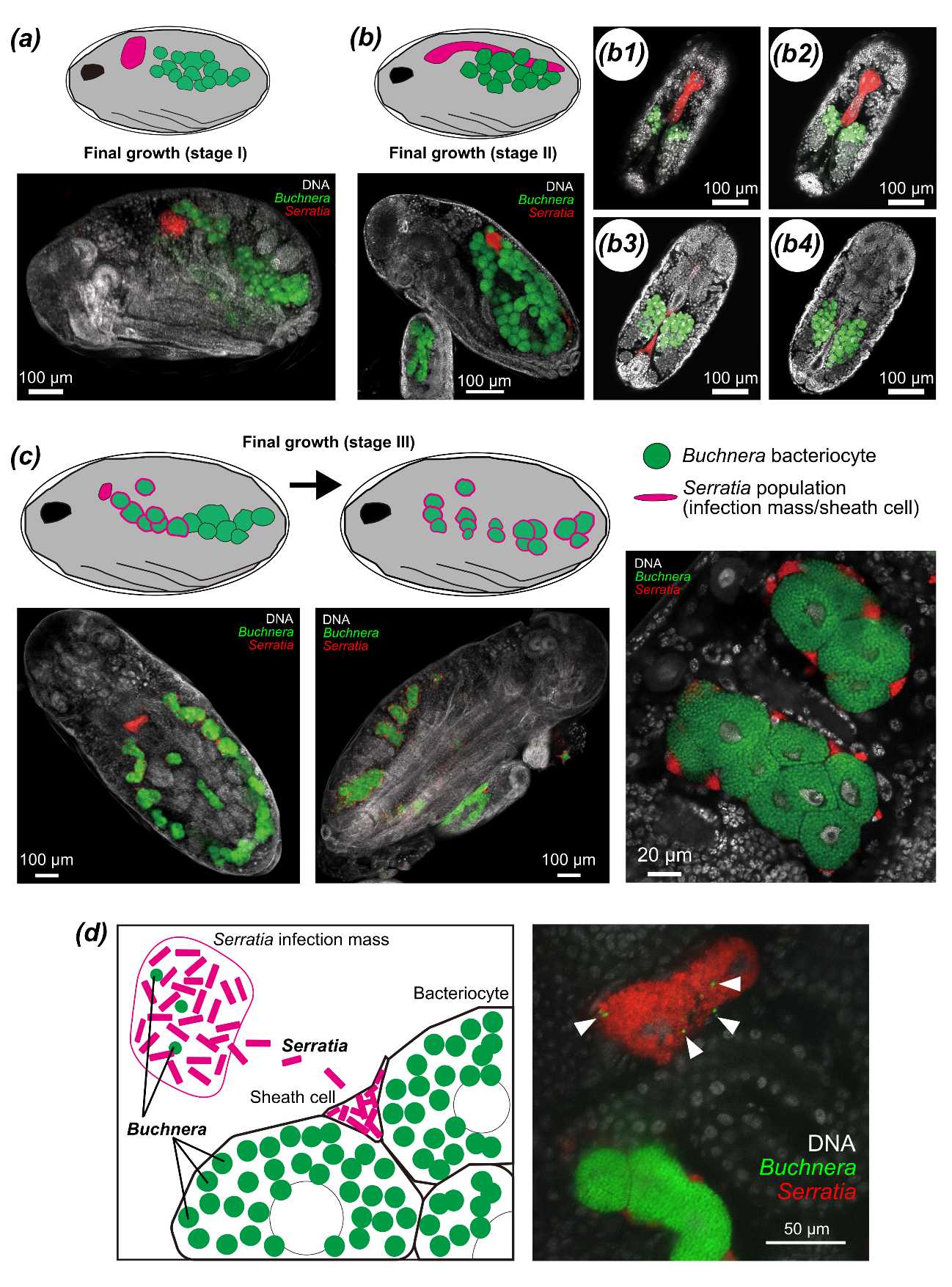
**Figure S11.** Late stages of viviparous embryonic development and bacterial distribution in *Lachnus tropicalis*. Illustrations and representative microscopic images showing the localization of symbiotic bacteria are presented. **(a)** In the first substage, *Buchnera* is housed within bacteriocytes, while *Serratia* forms a single large injection mass. **(b)** In the second substage, *Buchnera*-containing bacteriocytes actively proliferate. The *Serratia* infection mass is located dorsally, extending posteriorly from just behind the thorax towards the end of the abdomen. **(c)** In the third substage of the final growth stage, *Serratia* cells begin infecting sheath cells, first in anteriorly located cells and subsequently sequentially in posterior sheath cells. Ultimately, *Serratia* becomes localized in all sheath cells. Approximately 10 *Buchnera*-containing bacteriocytes and about twice as many *Serratia*-containing sheath cells form a single bacteriome. These bacteriomes are positioned laterally on the body, likely with two pairs per abdominal segment, and this arrangement is maintained throughout aphid life. **(d)** Schematic illustration (left) and microscopic image (right) showing the distribution of *Serratia* from the infection mass into sheath cells. Note the consistent presence of a few *Buchnera* cells within the *Serratia* infection mass, regardless of the developmental stage (white arrowheads in the right panel indicate *Buchnera* signals).

**Chapter 6: Review of the symbiotic system in *Lachnus roboris***

**Endosymbionts of *L. roboris***

We selected *Lachnus roboris* as a representative species that retained the ancestral *Serratia* for comparison with *L. tropicalis*. As a relatively well-studied European species within the genus *Lachnus*, existing data provide an excellent reference point for inferring the symbiotic state before the *new Serratia* acquisition in the genus. Therefore, we summarized the known information for *L. roboris*.

*L. roboris* belongs to a clade phylogenetically distinct from the Asian *Lachnus* species. Together with *L. pallipes* and *L. quercihabitans*, it forms a basal clade within *Lachnus* compared to the Asian *L. tropicalis* group (*L. tropicalis*, *L. shiicola*, *L. takahashii*, *L. siniquelcus*) (8, 55).

Previous studies (1, 2, 3) have suggested that *Serratia* Clade B is a co-obligate symbiont in *L. roboris*, as confirmed by 16S rRNA gene sequencing. Furthermore, amplicon sequencing by Manzano-Marín et al. (4) revealed the presence of *Buchnera*, *Serratia*, and *Wolbachia* in *L. roboris* and its closely related species, *L. pallipes*. Based on indirect evidence, including sequencing data and detailed histological observations by Klevenhusen (49), *Serratia* Clade B was inferred to be coccoid in morphology. Given that *Wolbachia* are generally rod-shaped, the rod-shaped symbionts described by Klevenhusen (49) likely correspond to guest symbionts such as *Wolbachia*.

Moreover, a draft genome of *Buchnera* from *L. roboris* has recently become available (6). This genome is 0.42 Mb in size as a chromosome, and no plasmid was registered. However, re-annotation using DFAST identified functional genes for the entire tryptophan synthesis pathway, except for *trpE* and *trpG*. Given that *Buchnera* commonly harbors a tryptophan plasmid (pTrp) containing *trpE* and *trpG*, probably, pTrp was not assembled because of technical issues. Similarly, given that the *Buchnera* genome of *L. tropicalis* contains pLeu, it is highly likely that *Buchnera* of *L. roboris*, a closely related and ancestral symbiotic species, also possesses pLeu, even if it was not detected in this particular assembly. This misassembly often occurs, especially in short-read-only assemblies (27). Therefore, in the discussion presented in the main text, we considered *Buchnera* from *L. roboris* to possess *trpE, trpG,* and *leuA–D*, which is a reasonable inference based on the above evidence.

**Summary of *L. roboris* symbiosis from Klevenhusen (1927, pp. 127-133)**

Klevenhusen (49) described three distinct symbionts in *Pterochlorus roboris* (a synonym of *L. roboris*): a large “aphid common” round symbiont (=*Buchnera*), a smaller coccoid symbiont (approximately 1 μm diameter) (=*Serratia*), and short rods (average approximately 4 μm) (=*Wolbachia* or other facultative symbionts). This study details the intricate spatial organization and transmission of symbionts within aphids.

***Spatial organization in adult aphids***

Coccoid symbionts (*Serratia*) are primarily localized within cells that appear to be infected sheath cells of the main bacteriome, positioned between typical bacteriocytes. Their distribution and cellular features support this interpretation. Fat cells are frequently found in close association with bacteriomes. Free coccoid forms also occur in the body cavity, where they accumulate near tracheae and interact with leukocyte-like cells, suggesting possible symbiont absorption. Rod-shaped symbionts (facultative symbionts such as *Wolbachia*) primarily colonize similar cells in the fat body and body cavity. Then often co-occur with coccoid forms in the same cells, although typically one type predominates. Rods are not consistently found within the bacteriome; when they are, they form a small, loosely arranged cluster between bacteriocytes, from which they can exit into the body cavity and subsequently aggregate in the gut.

***Embryonic infection and transmission***

During embryonic infection, a marked initial segregation occurs: the large primary symbionts (*Buchnea*) remain distinct from a mixed “jumble” of coccoid and rod-shaped forms. The primary symbionts first colonize the embryo, followed by coccoid and rod forms, which generate a “secondary infection mass” derived from maternal body cavity cells. Guided by diverging primary symbionts, this secondary mass migrates toward the anterior pole of the embryo. Some primary symbionts are incorporated laterally, whereas others degenerate within the secondary mass. Cocci and rods subsequently cap the primary mass.

As embryogenesis progresses, the secondary infectious mass undergoes temporary division and later reunification during gastrulation. Rearrangement of coccoid symbionts continues throughout development. Coccoid symbionts colonize bacterial sheath cells originating from the secondary mass and later extend into adjacent fat cells and other transient cells. They also become segregated into peripheral groups that are rod-free. Rod-shaped symbionts either migrate away from the infectious mass or are internalized by tissue cells, sometimes alongside coccoid forms. Ultimately, the infectious mass may completely divide or persist as a rod-enriched remnant.
